# Stereoretentive Post-Translational Protein Editing

**DOI:** 10.1101/2022.08.22.504816

**Authors:** Xia-Ping Fu, Yizhi Yuan, Ajay Jha, Nikita Levin, Andrew M. Giltrap, Jack Ren, Dimitrios Mamalis, Shabaz Mohammed, Benjamin G. Davis

**Author notes:** These authors contributed equally (Xia-Ping Fu, Yizhi Yuan).

## Abstract

Chemical post-translational methods now allow convergent side-chain editing of proteins as a form of direct chemical mutagenesis without needing to resort to genetic intervention. Current approaches that allow the creation of constitutionally native side-chains via C–C formation using off-protein carbon-centred C• radicals added to unnatural amino acid radical acceptor SOMOphile ‘tags’ such as dehydroalanine are benign and wide-ranging. However, they also typically create epimeric mixtures of D-/L-residues. Here we describe a light-mediated desulfurative method that, through the creation and reaction of stereoretained *on-protein* L-alanyl C_β_• radicals, allows C_β_–H_γ_, C_β_–O_γ_, C_β_–Se_γ_, C_β_–B_γ_ and C_β_–C_γ_ bond formation to flexibly generate site-selectively edited proteins with full retention of native stereochemistry under mild conditions from a natural amino acid. This methodology shows great potential to explore protein side-chain diversity and construct useful bioconjugates.

**Table of Contents Image:** 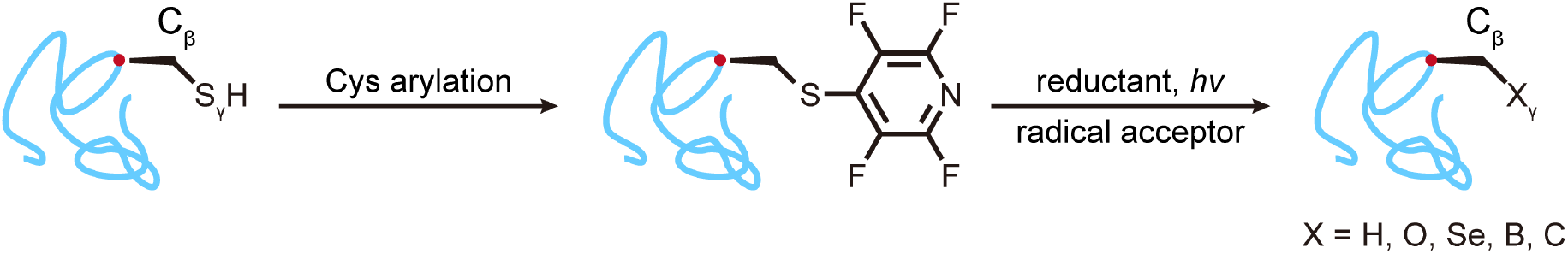

## Introduction

In nature, post-translational protein modification enables and mediates various essential biological processes.^1^ For example, glycosylation enables immune response,^2^ phosphorylation activates enzymes^3^ and ubiquitination triggers protein degradation.^4^ Whilst such natural post-translational modifications (PTMs) therefore extend the complexity of protein structures greatly and so increase the diversity of gene product / protein function, they do not cover all possible chemical space and so in principle useful unnatural functionality in biology remains undiscovered. The editing of proteins to create such ‘chemical PTMs’ could efficiently bridge that gap.^5^ Moreover, the recapitulation of PTM (and other protein) function through precise structural design allows causal links to be established to putative mechanisms.^6^

Classical strategies for protein modification often feature bonds to heteroatoms (non-carbon) made at the γ (Cys S_γ_, Thr O_γ_, Ser O_γ_) or ω (Lys N_ω_, Tyr O_ω_) positions of side chains,^7, 8^ These have valuably allowed technological and translational development of novel diagnostic and medical tools as well as the interrogation and the manipulation of biological processes.^5, 9, 10^ However, constructing C_β_–X bonds *via* C_β_, which is present in all amino acid side chains, is a rare but potentially far-reaching disconnection in synthetic and chemical biology (**Figure 1a**).^11^ From a retrosynthetic viewpoint,^12^ the major difficulty of utilizing C_β_ to catenate is that appropriate synthetic equivalents of a protein synthon at C_β_ are difficult to generate under mild conditions using classical heterolytic / 2e-chemistries as the C+ or C-equivalents (**Figure 1a**) that are required are typically quenched.

**Figure 1.**
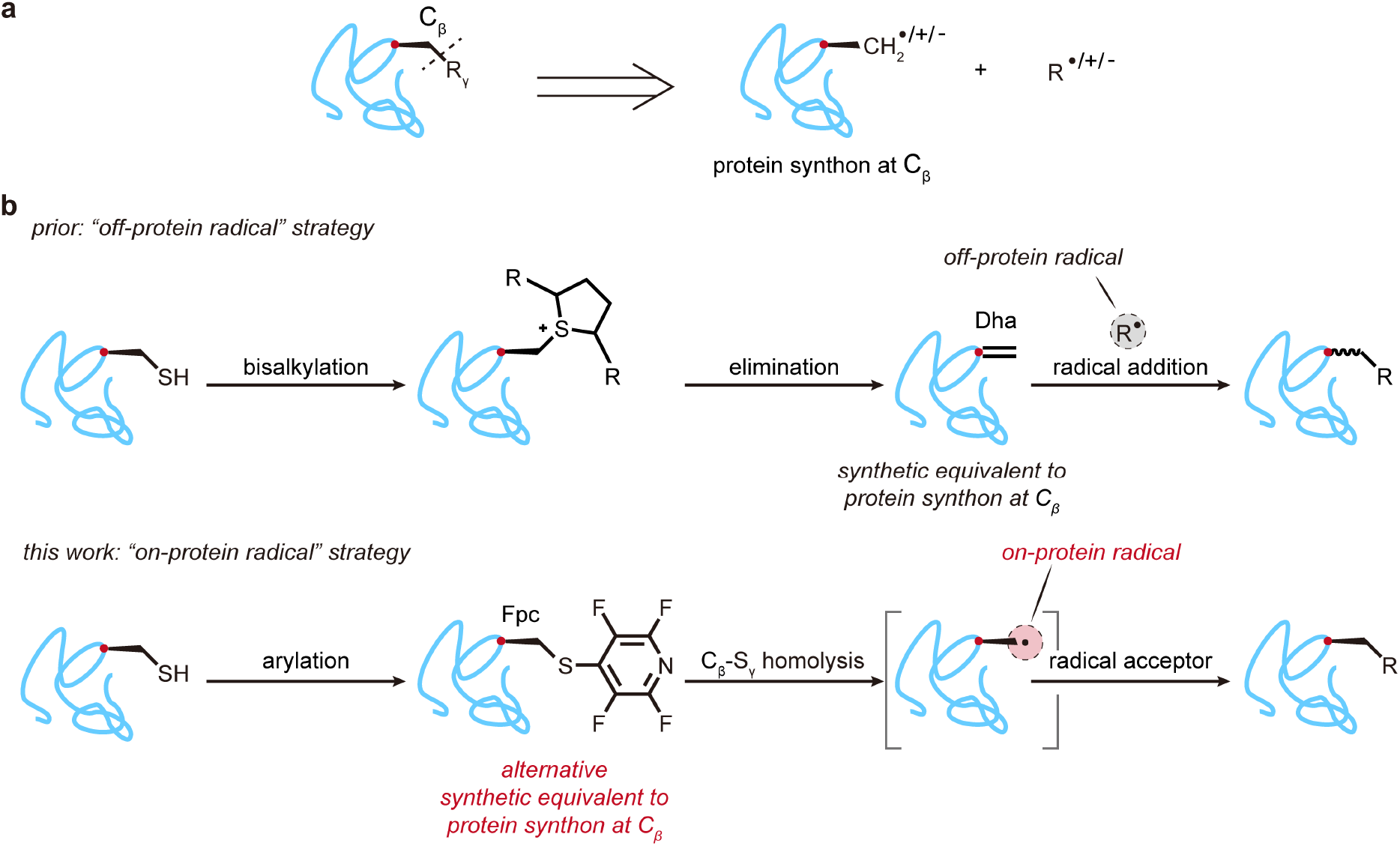
Strategies for C_β_-X_γ_ bond formation on Proteins. (**a**) Retrosynthetic analysis of *in situ* side chain C_β_-X_γ_ bond formation. (**b**) Contrasting methods for use of synthetic equivalents to a C_β_ protein synthon using, prior, off-protein or, this work, on-protein carbon-centred C• radicals. The use of on-protein radicals allows the potential for retention of stereochemistry via a configurationally defined L-alanyl radical intermediate, arbitrarily denoted in protein image ‘cartoons’ throughout this manuscript via the use of a ‘wedged’ bond.

To address this problem, we^11–14^ and others^15^ have considered the use of homolytic / 1e-chemistries that may be used in combination with readily-generated^15–19^ dehydroalanine (Dha) residues on proteins. In this way, Dha, by acting as singly occupied molecular orbital (SOMO) acceptor (‘radical acceptor’ or ‘SOMOphile’), can serve as a synthetic equivalent of a protein synthon at C_β_ by reacting with off-protein carbon-centred C• radical species. This has allowed selective C_β_–C_γ_ bond formation to introduce a wide variety of side chains into several protein scaffolds.^12–14^ Although this ‘off-protein radical’ strategy – radical acceptor (Dha) on-protein reacts with C• radical species generated off-protein – allows ready exploration of protein side-chain diversity, modification state and consequent function, the native L-stereochemistry at the modified residue is erased (**Figure 1b, top**) and typically regenerated with low diastereoselectivity. This results in the creation of a mixture of D-/L-epimers in d.r.s of typically ~1:1 (and < 3:1). Moreover, whilst function can often be reliably inferred from such epimeric mixtures,^12, 14, 20^ access to a stereodefined chemical method would undoubtedly be advantageous in removing the ambiguity of the potential role of the D-epimer in such analyses and in considering synthetic chemical routes to pure protein products. Here, we describe a ready stereoretentive method for achieving this through the strategic inversion of this homolytic / 1e-disconnection to allow efficient use of *on-protein* Cβ• radicals which retain their L-configuration (**Figure 1b, bottom**).

## Results

### *Design of a method for selectively creating on-protein* Cβ• *radicals*

In re-examining methods for the refunctionalization of Cβ in proteins to create putative synthons, we noted that for one synthetic equivalent, Dha, one of the most prevalent methods exploits the elimination of an activated sulfonium intermediate, generated *via* the chemoselective alkylation of a Cys residue.^16^ The strategic success of this method therefore relies in part on the ability to chemoselectively access a suitable synthetic equivalent (that itself can then be manipulated chemoselectively). For Dha formation via Cys this relies in part on the ability to react free Cys in the presence of unreacted cystinyl S–S bonds.

We considered that if a pre-activated (e.g. through suitable modification or ‘alkylation’) Cys-derived C_β_-S_γ_ bond underwent 1e-homolysis instead of an heterolytic 2e-E_1_cB process,^21^ then an L-alanyl-radical Cβ• might be generated without influencing the configuration of the stereogenic L-C_α_ centre. This L-alanyl-radical Cβ• might then be subsequently trapped by radical acceptors (SOMO-philes) to form new side chains. This ‘on-protein radical’ – radical acceptor is off the protein and radical is generated on the protein – strategy could theoretically avoid the loss of native stereochemistry (**Figure 1b, bottom**).

Notably, it has been proposed for over 60 years that alanyl radicals may be intermediates in Cys desulfurization reactions,^22, 23^ and such reactions are now commonly exploited in so-called ‘traceless native chemical ligation’^24–26^ to convert Cys to desulfurized Ala residues. In peptidic systems alanyl-radicals generated in this way have shown promise by taking advantage of phosphine to activate the C_β_-S_γ_ bond, limited to a peptide scaffold.^27, 28^ Such prior strategies for desulfurization at cysteine, cystine or selenenylcysteines proceed via a seemingly complex or likely multiple-manifold process^29^ involving the likely intermediate formation of thiophosphoranyl radical adducts as precursors to C• radicals formed upon β-scission.^22, 23^ The requirement in these systems for use of phosphines or other P(III) reagents, which are strongly reducing, effectively precludes more general use in typical protein systems since these are commonly used to disrupt disulfides (e.g. TCEP) at the concentrations required to effect desulfurization via thiophosphoranyl. This, and the sometime additional requirement of organic (co)solvent, have therefore prevented their efficient use in a general manner in protein systems beyond their use in ‘traceless’ ligation methods via C–H bond formation prior to refolding. We have shown that eliminative mechanisms to Dha may compete in some phosphine mediated desulfurization manifolds thereby raising the potential for loss of stereochemistry or side-reaction.^16, 30^

We have also shown that such on-protein C• radicals, when stabilised by α-fluoro-substitution as C(F)_n_•, allow reactivity that enables C–Se, C–O and C–C bond formation.^14^ Nonetheless, despite the promise suggested by all of these above methods they all require either the use of conditions that are not compatible with typical proteins or the creation of unnatural (e.g. fluorine substituted) sidechain precursors to access C• radicals.

Our generation of sidechain on-protein carbon-centred radicals from sulfonyl precursors through the cleavage of Cγ–S(O)_2_R bonds through reductive initiation highlighted the feasibility of C–S bond scission for radical generation in proteins. Whilst the redox potential and C–S bond strength to allow initiation at such sites may potentially be tuned^31^ our attempts to selectively generate suitable alkyl sulfonyl Cβ– S(O)_2_R sidechains at the β-carbon rather than γ-carbon directly from modified Cys residues proved challenging due, in part, to concomitant oxidation of other residues such as methionine.

Therefore, we turned to alternative methods for tuning the radical scission potential of the Cβ–S bond. The presence of electron-withdrawing substituents on S is known to enhance C–S bond cleavage via homolytic and mesolytic manifolds.^32, 33^ In reductive initiation this may stabilize appropriate radical anion intermediates formed upon single-electron transfer (SET) and/or thiolates in mesolysis. Such SET driven initiation may also be light-stimulated (either in the reductant e.g. photoredox catalyst or in the substrate).^34^

Initial^35,36^ *ab initio* DFT calculations (see **Supplementary Information**) suggested that the appropriate precursor MO energies and associated stability of a resulting radical anion might be effectively altered through the attachment of strongly electron withdrawing substituents on a simple sulfide, and especially those allowing *π*-acidic, conjugation effects. These could potentially be derived directly from the free thiol SH of Cys if an effective method for selective modification could be established. This therefore suggested that the installation of an electron withdrawing *π*-system on Sγ might create a suitable Cys-modified precursor for a Cβ• radical via Cβ–Sγ bond homolysis; DFT calculations also suggested an associated bond lengthening of the alkyl S–Csp^3^ and not the aryl S–Csp^2^. Whilst our calculations suggested different possible Cys-Sγ substituents might prove fruitful, we therefore turned our attention to the installation of electron-poor aryl moieties that could be readily achieved with chemoselectivity via direct protein arylation (**Figure 1b, bottom**).

Whilst we^37^ and others^8^ have shown that S-arylated motifs may be readily installed through metal-mediated arylation, we also considered more classical methods that allow S_N_2Ar type reactions for installation of such moieties.^38^ For example, elegant studies have shown that benzenoid systems may be installed in such a way into proteins, even exploiting selectivity by proposed exploitation of interactions of local residues (a so-called “pi-clamp”).^39^ More recently, the tuning of these benzenoids with electron-withdrawing substituents has further allowed enhanced selectivities for such benzenoid conjugations.^40^ However, computation (see **Supplementary Information**) suggested that certain aryl moieties might fail to stabilize required intermediates on the putative pathway required for reductive initiation.

S_N_2Ar reactions at *hetero*arenes are often more greatly favoured than their arene counterparts. One archetype, pentafluoropyridine (**pyF_5_**) possesses enhanced reactivity^41–43^ with respect to S_N_2Ar, attributed in part to the ability to delocalize developing negative charge in transition states around any Meisenheimer intermediate to the heteroatom.^42^ Such heteroatom-associated stabilization would likely also contribute on the pathway to initiation via a radical anion intermediate. This conclusion was not only supported by our computation (see **Supplementary Information**) but also further confirmed by experimental measurement of half potentials for the reduction of pyF_4_-sulfides^44^ (PyF-S) to radical anion intermediates that might potentially allow initiation via Cys-Cβ–Sγ bond homolysis.

We therefore reasoned that, although unprecedented, by combining a two-step process of chemoselective arylation (with such an electron poor heteroaryl) with reductively-driven initiation, we might create a ready and direct pathway to stereoretained, on-protein L-alanyl radical intermediates (via Cys-Cβ–Sγ cleavage) suitable for further addition reactions. Here, we demonstrate that either direct single electron transfer (SET) process or electron donor-electron acceptor (EDA) complexes in this way allow us to construct C_β_-H_γ_, C_β_-O_γ_, C_β_-Se_γ_, C_β_-C_γ,_ C_β_-B_γ_ bonds on proteins. (**Figure 2,3**).

**Figure 2.**
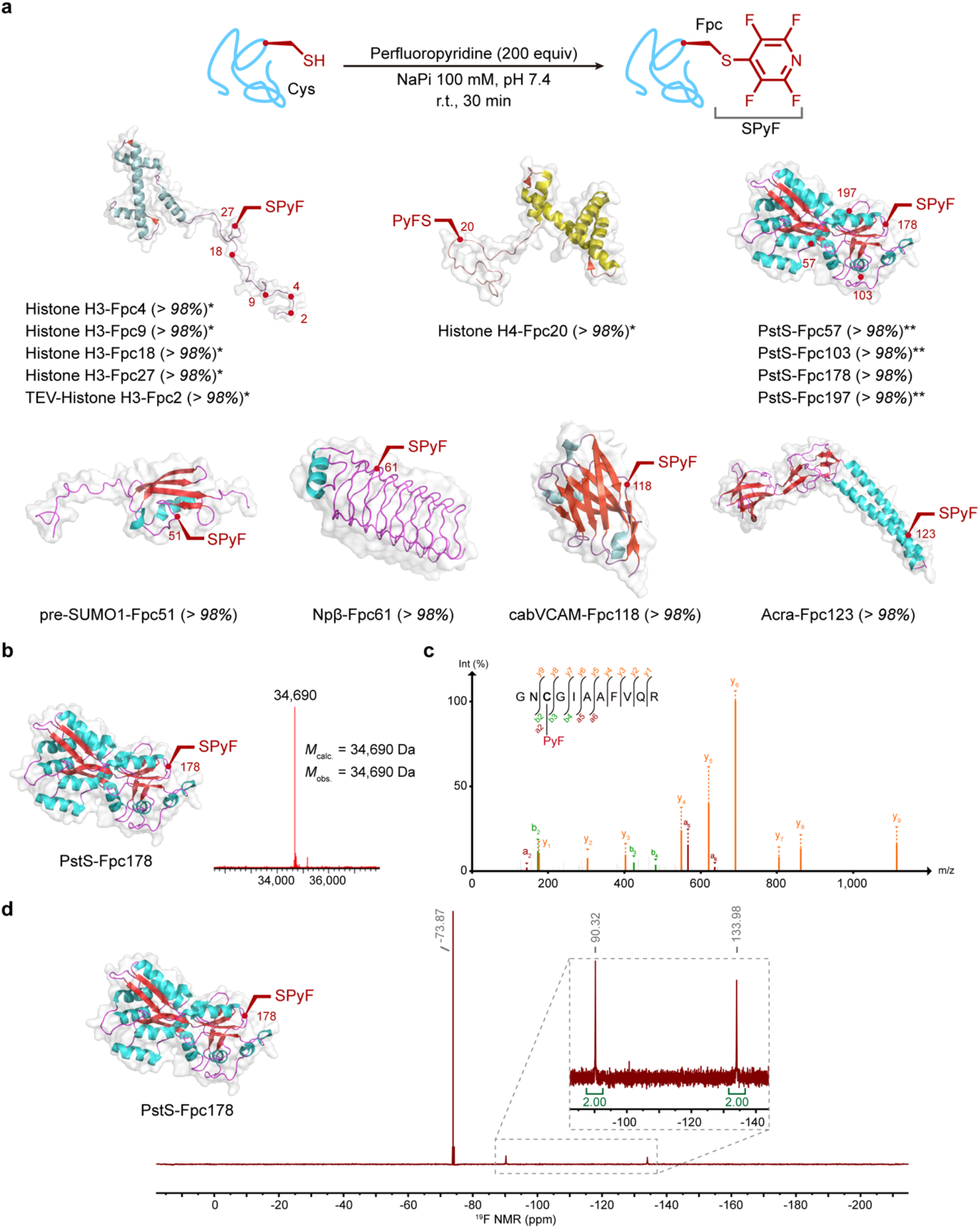
Site-selective chemical introduction of the Fpc sidechain into proteins. (**a**) The chemoselective modification of Cys allows site-selective introduction of Fpc at a range of sites in representative proteins with a varied scaffold type and at different secondary structure motifs with. (**b**) Representative intact protein MS confirms excellent conversion to install Fpc, shown here for PstS-Fpc178. (**c**) Site-selectivity was confirmed by tryptic-MSMS analyses, shown here for PstS-Fpc178^176–186^ peptide. (**d**) Advantageously, intact protein ^19^F NMR allows sensitive assessment in a ‘zero-background’ of reaction chemoselectivity. Characteristic chemical shifts (δ_F_1 ~ −90.3, δ_F_2 ~ −134 ppm)^45^ confirmed selective C–S product formation to generate Fpc, shown here for PstS-Fpc178. * = in Tricine buffer (100 mM pH 7.4); ** = in NaPi buffer (100 mM pH 7.4 with Gdn·HCl 3 M).

**Figure 3.**
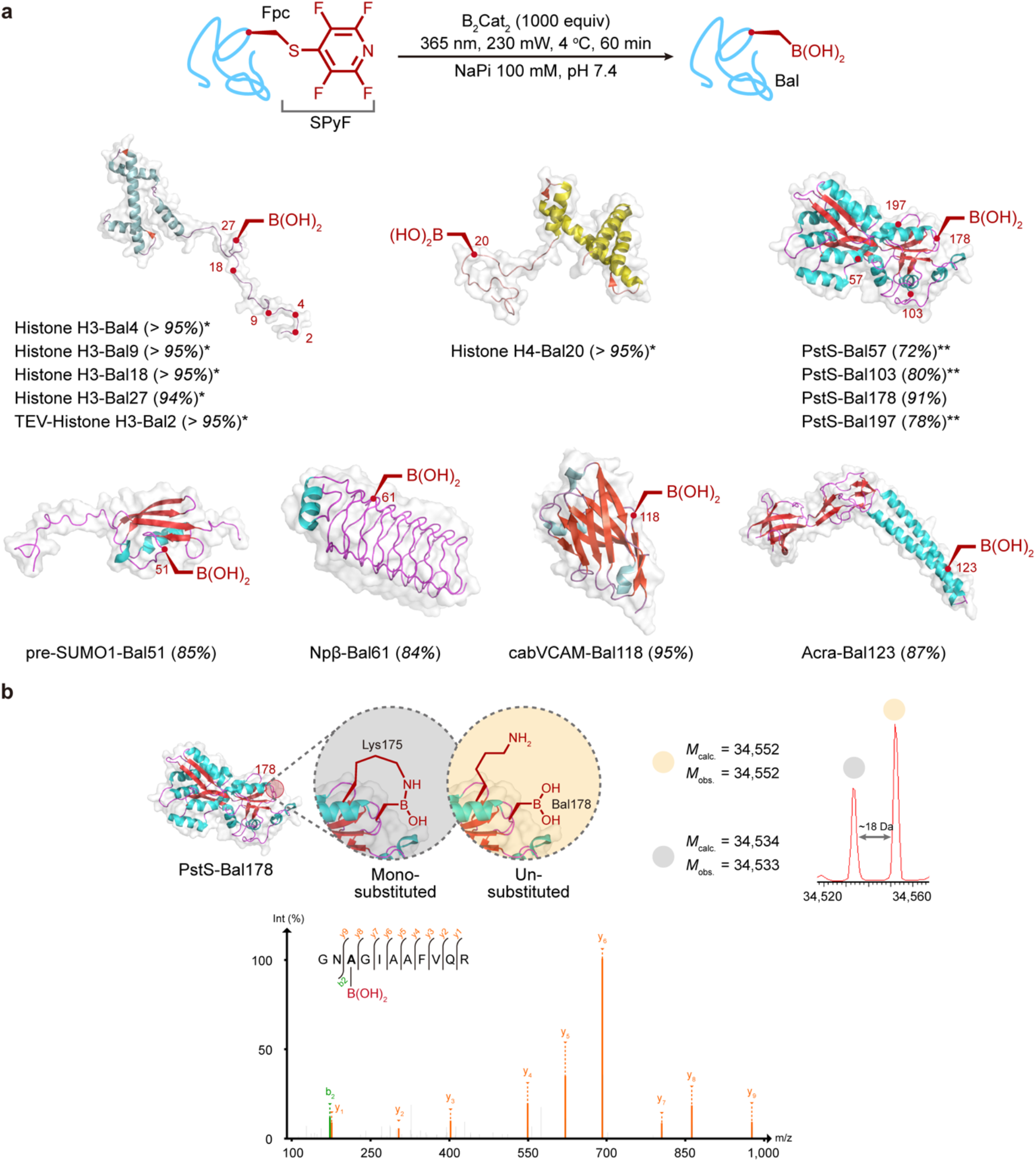
Chemical introduction of L-boronoalanine (L-Bal) into proteins. (**a**) Dual initiation and trapping from Fpc allows the introduction of Bal into diverse protein sites and scaffolds. (**b**) Consistent with prior observations,^58^ the insertion of Bal is observed in multiple states via intact protein MS concomitant with internal ligated boronates in intact protein (top), attributed here speculatively to nearby Lys175 as an illustrative example only. Bal can be observed directly via tryptic-MSMS (bottom). Data shown here for representative system PstS-Bal178. * = in Tricine buffer (100 mM pH 7.4); ** = in NaPi buffer (100 mM pH 7.4 with Gdn·HCl 3 M).

### Site-selective chemical introduction of tetra**<u>f</u>**luoro**<u>p</u>**yridyl-**<u>C</u>**ys (Cys-S-PyF or **Fpc**) into proteins

Perhaps due to its known enhanced S_N_2Ar reactivity in classical small molecule systems,^42^ **pyF_5_** has been perceived to be a non-selective modification reagent,^38^ in part based on exploration of peptidic systems in DMF.^45^ Whilst this is a reactivity that can tuned by use of protic solvent trifluoroethanol^46^ its use in aqueous systems has not been exploited. The creation of tetra**<u>f</u>**luoro**<u>p</u>**yridyl-**<u>C</u>**ys (Cys-S-PyF / **Fpc**) in proteins is therefore unexplored.

The reaction between a model protein containing a single cysteine, AcrA-Cys123 and pentafluoropyridine (**pyF_5_**) was evaluated as an initial model. AcrA is a challenging model substrate membrane protein that would allow us to test the limits of this method. Whilst initial attempts explored the use of lower temperature (4 °C) to control reactivity this ultimately proved unnecessary. Instead, pH proved an important determinant. Thus, whilst under neutral or even mildly acidic conditions (pH 6) reaction proceeded only slowly, strikingly perfluoroheteroarylation with **pyF_5_** could proceed efficiently in different kinds of buffers at pH > 7.0 (**Supplementary Table 1**). Optimized conditions (pH 7.4, 200 equiv. **pyF_5_**, 25 °C) were effective in a range of buffers (NaPi, tricine or Tris) and allowed full conversion to AcrA-Fpc123.

With a reliable method in hand, a variety of proteins were screened, resulting in all cases full conversion to tetrafluoropyridyl-cysteine containing proteins (**Figure 2**): histones H3 and H4, small α-helical nuclear proteins; PstS, a protein involved in bacterial phosphate transport;^47^ pre-SUMO1 (SUMO, small ubiquitin-like modifier), a small globular protein containing α-helices and β-sheets; cabVCAM, a cross-reactive nanobody against human and murine VCAM1;^48^ Npβ, a β-helical pentapeptide repeat;^49^ and AcrA, a membrane protein. Characterization (including intact protein mass spectrometry (MS), proteolytic/tandem MS (MS/MS): **Figure 2b,c,d** and **Supplementary Information**) confirmed that proteins were successfully and site-selectively modified with perfluoropyridine.

Notably, the strong dual ^19^F resonances in the Fpc sidechain also allowed unequivocal confirmation of the formation of C–S product formation via direct use of ^19^F protein NMR (δ_F_1 = −90.32, δ_F_2 = −133.98 ppm) wholly consistent with observations in peptidic systems (δ_F_1 = −90.37, δ_F_2 = −134.15 ppm)^45^ and highlighting a lack of modification of other putative protein nucleophiles (e.g. Lys: δ_F_1 = −98.17, δ_F_2 = −165.54; Tyr: δ_F_1 = −91.32, δ_F_2 = −155.98; Ser: δ_F_1 = −97.10, δ_F_2 = −165.92).^45^ The ability to use this ^19^F signal in the ‘zero-signal’ background of native proteins is a striking additional advantage of the Fpc sidechain as an intermediate protein ‘tag’ state (**Figure 2d**).

### Light-mediated C_β_-S_γ_ bond cleavage testing

With the establishment of a reliable method to install Fpc into a range of protein substrates, we then tested its potential in light-mediated C_β_–S_γ_ bond cleavage on a protein scaffold. Our prior generation of on-protein radicals had successfully exploited reductive initiation.^14^ Multiple reductants were screened to drive putative reductive initiation using protein PstS-Fpc178 as a test substrate (**Table 1** and **Supplementary Figure 1**).

**Table 1.**
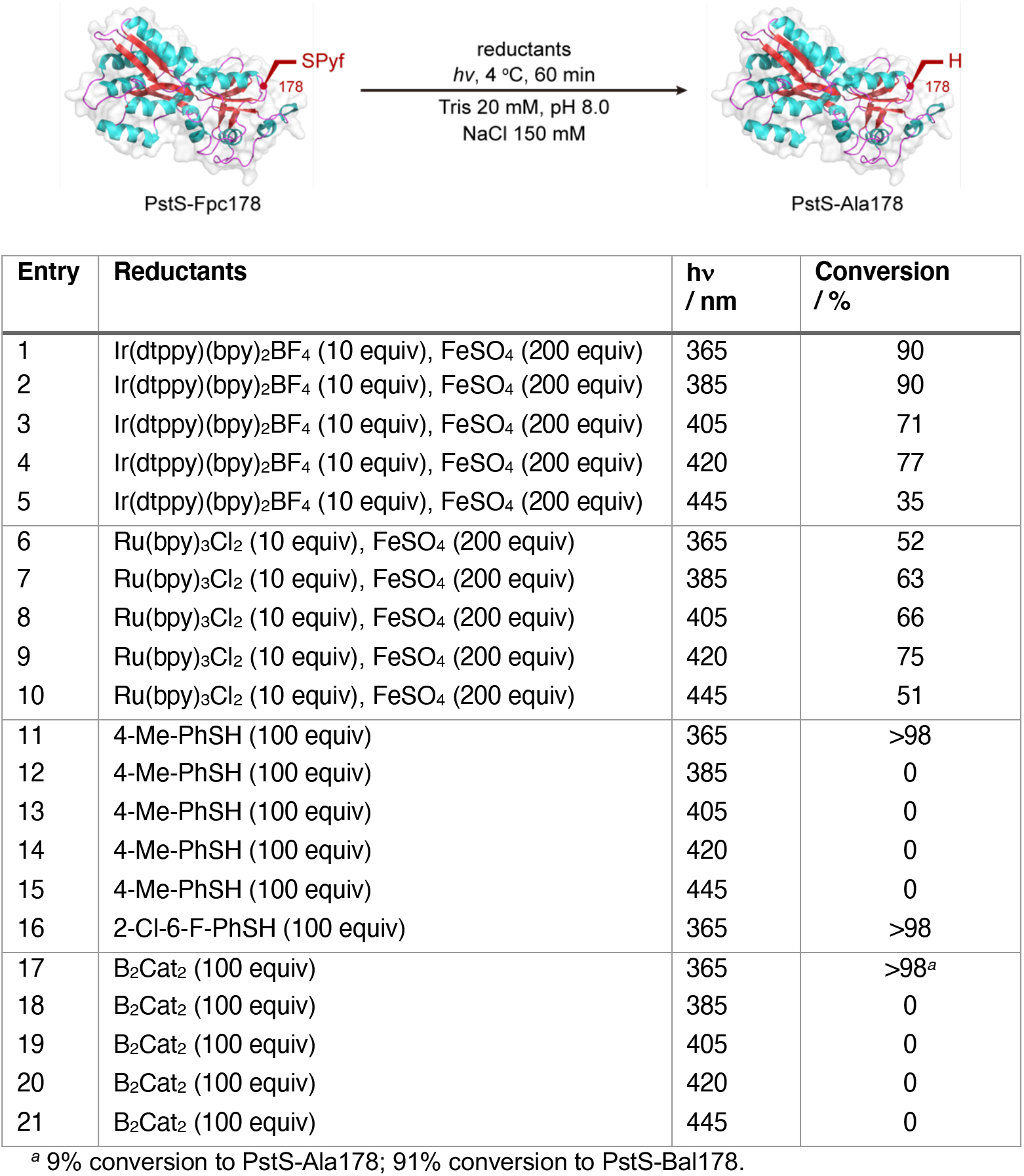
Light-mediated C_β_-S_γ_ Bond Cleavage in Fpc-containing Proteins. Exploration of varied SET reagents revealed differing modes of reductive initiation and optimal conditions. Conditions (see also scheme): in a glovebox at < 10 ppm O_2_, PstS-Fpc178 (15 μM, 50 μL), reductants were mixed and the reaction irradiated at 4 ° C for 60 min. The reaction mixture was then analysed by LC-MS.

Pleasingly, both strongly reducing [Ir(dtbbpy)(ppy)_2_]PF_6_ (E_ox_ −1.51 V vs SCE) and less reducing Ru(bpy)_3_Cl_2_ (E_ox_ −1.33 V vs SCE) photostimulated outer-sphere SET metal complexes (‘photoredox’ catalysts) drove reaction at a range of wavelengths (365 - 445 nm) in the presence of Fe(II) as co-reductant^14^ to give PstS-Ala178 (**Table 1**). Whilst the conversions with [Ir(dtbbpy)(ppy)_2_]PF_6_ proved typically higher (up to 90%) even the milder Ru(bpy)_3_Cl_2_ proved effective (up to 75% conversion); the former also generated some apparent oxidative damage in proteins. The implied breadth of the action spectrum for photoreaction is consistent with the broad absorption excitation region of both of these complexes.

Whilst these reactions proved successful, the associated issues of conversion and damage led us to consider alternative systems and, so, alternative reductants. Two non-metal chemical reductant types were therefore considered: aryl thiols and diboron(IV) compounds. Both classes importantly encompassed the use of reagents with the potential to act in a photostimulated manifold via putative charge-transfer complexes.^50, 51^

With both arylthiols, *para*-tolyl-SH (*p*Tol-SH) and 2-chloro-6-fluoro-phenyl-SH (ClFΦ-SH) gave excellent conversions under mild conditions of PstS-Fpc178 → PstS-Ala178. It should be noted that whilst these aryl thiols represent differently tuned acidities (pK_a_s *p*Tol-SH = 6.82 in water,^52^ ClFΦ-SH = 5.27 predicted in water^53^) neither are sufficiently potent nucleophilic reductants to disrupt protein S–S bonds.^54^ In all cases the activated C_β_–S_γ_ bond in the Fpc sidechain could be successfully cleaved under 365 nm light but not at other wavelengths (**Table 1**). Notably, this narrow implied action spectrum, in contrast with the observed outer-sphere metal complexes (see above) was consistent with the formation of a corresponding charge-transfer complex enabling donor-acceptor^55^ SET (see below) as well as the putative reductive capacity of, for example, the thiolate/thiyl half reaction.^56^

Finally, the aryl diboron(IV) compound *bis*(catecholato)diboron (B_2_Cat_2_) was also tested. This too proved effective under mild conditions (**Table 1**) in driving cleavage of the activated C_β_–S_γ_ bond in PstS-Fpc178, also with a narrow action spectrum (365 nm). Notably, however, this reaction of PstS-Fpc178 led to the formation not only of PstS-Ala178 but also concomitant formation of boronylated PstS-Bal178 as, indeed, the major product. Excitingly, this implied not only the potential for reductive initiation of PstS-Fpc178 but also the trapping of a putative intermediate on-protein Cβ• radical through coincident B–B bond cleavage by B_2_Cat_2_ allowing direct on-protein Cβ–Bγ bond formation.

### Stereoretentive introduction of L-boronoalanine (Bal) into proteins using dual initiation-trapping

This observed, dual, light-mediated reductive initiation and trapping using B_2_Cat_2_ led us to test the greater breadth of such concomitant on-protein L-alanyl trapping, here in Cβ–Bγ formation. Whilst organoboronic acids and their esters are vital building blocks that play a pivotal role in organic synthesis,^57^ there are few chemical methods to introduce boronic acid groups to proteins.^58^ Currently, the minimal borono amino acid, boronoalanine (Bal) cannot be introduced without dilution of homochirality at C_α_.^58, 59^ Since use of epimeric D-/L-Bal on proteins already exhibits the benefits of *de novo* binding function in expanding biological function,^58^ the ability to control such function at a homochiral residue could allow more precise dissection of associated mechanisms and ligand sequestration.

Strikingly, application of a 1000 equiv excess of B_2_Cat_2_ combined with photostimulated reductive initiation at 365 nm led, under a reduced oxygen atmosphere (< 10 ppm) to concomitant trapping and hydrolysis to form L-Bal (see also below) directly in proteins with excellent efficiency on seven different protein scaffolds at multiple pre-determined sites (**Figure 3** and **Supplementary Figure 2**).

### *Testing the breadth of* Cβ· *alanyl radical trapping*

This promising indication of C–B bond formation via reaction of an on-protein Cβ· radical led us to test the breadth and utility of L-alanyl C· trapping. Given the observed dual reduction-trapping activity of B_2_Cat_2_ we sought first to separate reductive initiation from trapping. To accomplish this, we tested the utility of the two aryl thiols, *p*Tol-SH and ClFΦ-SH, as reagents that could be varied in not only their pKa but also potentially their associated SET and hydrogen atom transfer (HAT) activities via tuning of their substituents^60^ in concert with several different kinds of representative radical acceptors that would allow formation of varied Cβ–Xγ bonds.

First, using *p*Tol-SH as reductant, we explored direct Cβ–Oγ formation using 2, 2,6,6-tetramethyl-1-piperidine-1-oxyl (TEMPO) as a persistent radical that might trap. Pleasingly, this combination enabled separation of reductive initiation (by *p*Tol-SH) from Cβ· trapping allowing conversion in 90% (**Figure 4a,b** and **Supplementary Figure 3**) from the alanyl radical with formation of a Cβ-Oγ bond.

**Figure 4.**
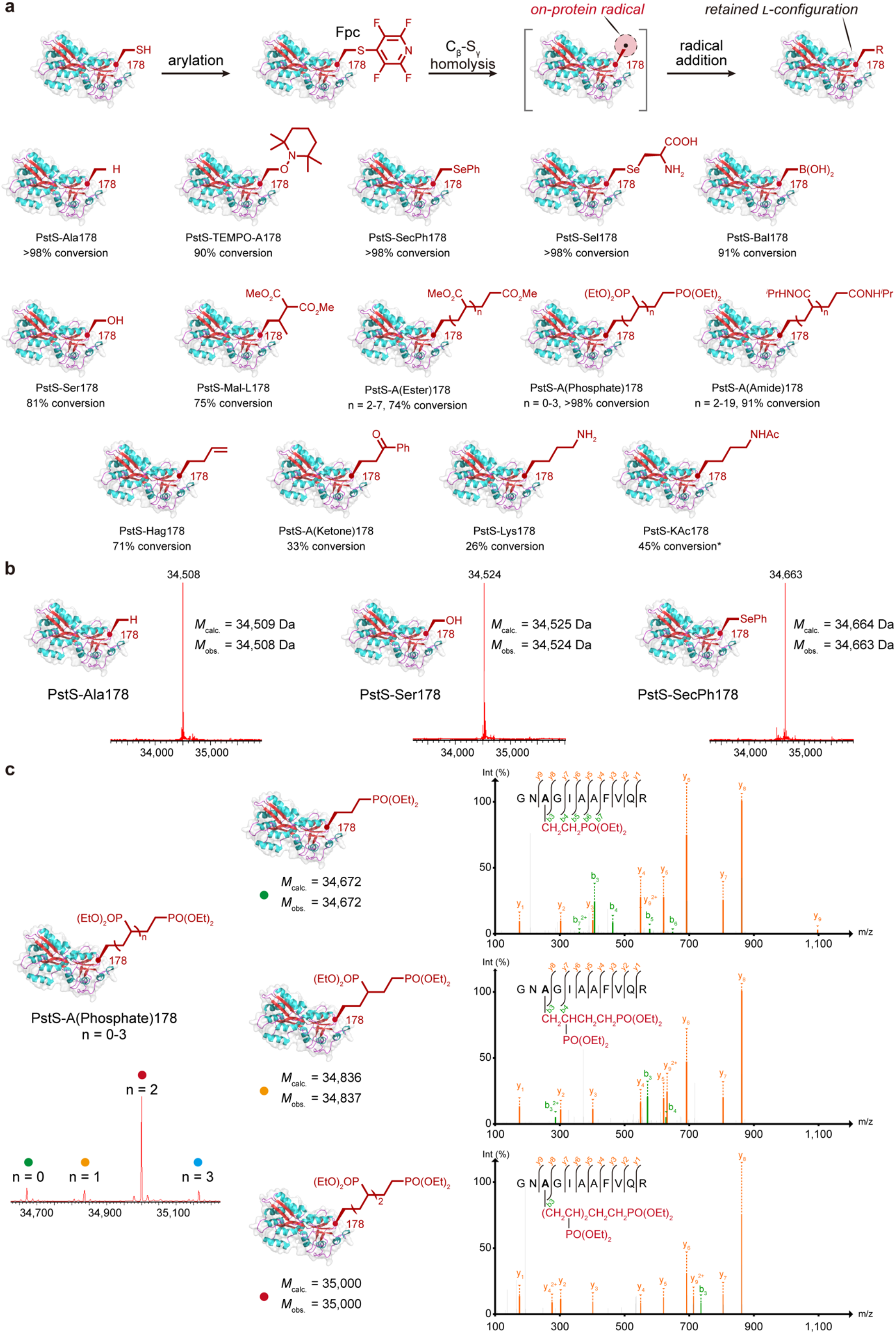
The Scope of L-Alanyl radical Trapping. (**a**) Diverse bond-forming proves possible through trapping of the on-protein radical generated from PstS-Fpc178 to generate varied sidechains. (**b**) Representative examples of Cβ–Hγ Cβ– Oγ and Cβ–Seγ bond formation proceed with excellent conversions. (**c**) Cβ–Cγ bond formation allows differing modes of bond formation. For example, on-protein C–C-polymerization where individual oligomer states can be observed by both intact protein MS (see **Supplementary Information**) and precisely mapped by tryptic-MSMS analyses (bottom). Direct Cβ–Cγ trapping without polymerization may also be achieved with differing substrates. In all cases, conversions are essentially full and the major side product formed in lower yielding reactions is the reduced Ala product. * = for KAc formation in PstS, formation of KAc includes 40% ‘dimer’ formation; interestingly, in histone H3 at site 18 only KAc product is observed (see **Figure 5**).

Diselenides could also trap the alanyl radical to form C_β_-Se_γ_ bond, creating modified selenocysteine residues; selenocysteines can exhibit relevant, typically redox, activities in natural systems^61^ (**Figure 4a,b**). Notably, despite the strong potential for direct nucleophilic heterolytic reduction of the diselenides by thiol, the use of ClFΦ-SH as a tuned SET reductant proved effective, allowing essentially complete conversion to both PstS-SecPh178 (using PhSe-SePh) and the selenolanthionine (Sel) adduct PstS-Sel178 (**Supplementary Figures 4,5**).

Most importantly, Cβ-Cγ bonds could be constructed, providing one of nature’s most important sidechain structural motifs (see above), when treated with appropriate polarized olefins as radical acceptor traps (**Figure 4a,b**). Notably, use again of the low nucleophilicity thiol ClFΦ-SH allowed reaction without apparent concomitant inhibitory side-reaction (e.g. Michael-type). The consequent reactivity was also determined by the nature of the radical acceptor. Thus, whilst expectedly^62^ vinyl phosphonate, acrylate and acrylamide, which give rise to stabilized adduct radical products that can react further via matched polarity allowed on-protein polymerization (n = 1-4, **Figure 4c**) acceptors that give rise to more fleeting intermediate radicals or ones with non-matched polarities or lower consequent addition allowed simple addition.

In this way, dimethyl ethylidenemalonate, 1-phenyl-1-trimethylsiloxyethylene, and phenyl allylsulfone also allowed ready formation of Cβ–Cγ bonds as direct or indirect adducts (**Supplementary Figures 6-8**). The indirect adducts thereby allowed the generation of side-chains with functional groups that may be further reacted as chemoselective ‘tags’ in protein chemistry. The former provides a reactive acetophenone carbonyl-containing moiety with the consequent potential for further application in diverse bioconjugation.^63^ The latter allows chemical generation of L-homoallylglycine (Hag), an archetypal ‘tag’ sidechain for thiyl-ene ligations^64^ The ability to introduce this chemically now complements its typical introduction via sense-codon reassignment.

Whilst typically poor electrophiles for protein modification, some of these Cβ– Cγ bond-forming reagents C acrylates and acrylamides nonetheless have the potential to non-specifically modify protein nucleophiles. Notably, in control reactions in the absence of thiophenol, protein substrate was not consumed. Moreover, when thiophenol alone was used to first reduce Pst-178Fpc to Pst-178Ala and then treated with further portions of thiophenol and acrylate or acrylamide no further reaction was is detected. These control experiments suggest that any such competing side reactions are negligible and that only radical-trapping reactivity is seen under these conditions that we describe here (see also Discussion).

Importantly, such Cβ–Cγ bond formation also allowed direct access also to fully native L-residues or their post-translationally modified variants. Thus, through the use of allylic amines (allylamine or its acetamide) fully-native L-lysine and L-*N*-acetyl-lysine residues were generated not only in a natural Lys site in histone H3 (H3-KAc18, see **Figure 5** and **Supplementary Information**) but also in an unnatural site in PstS (PstS-Lys178 and PstS-KAc178, **Supplementary Figures 9,10**), albeit in more modest yields. Interestingly, the reactivity of the acetamide appears to sit at a cusp under these conditions that leads to concomitant dimerization at some sites (e.g. unnatural site 178 in PstS but not at natural site 18 in H3).

**Figure 5.**
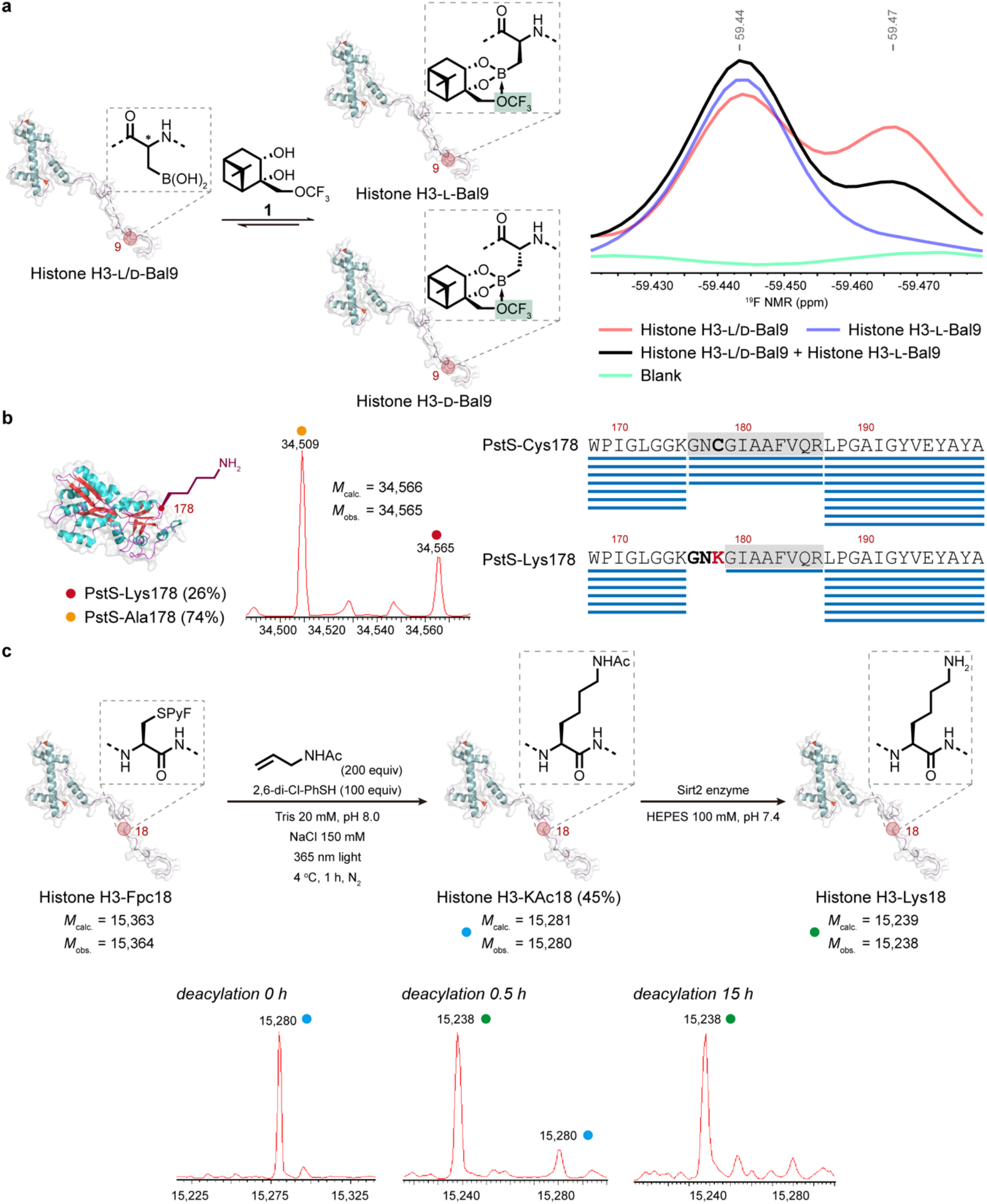
Functional and spectroscopic characterization of the retention of native L-stereochemistry. Different sites, proteins and sidechains were assessed for the configuration of the residues edited into corresponding proteins. (**a**) In NMR ‘chiral shift’ reagent diol **1** allowed use of intact protein ^19^F NMR to assess the configuration of Bal introduced into histone H3-Bal9. Comparison of that introduced with poorly stereoselective methods^58^ (mixture of D/L-, red) with that formed using the methods described here (blue), as well as the corresponding ‘mutual spike’ (black) suggest full formation of L-Bal using the editing methods described here within detection limits. (**b**) The observation of full cleavage of chemically-edited Lys178 in PstS-Lys178 by stereospecific enzyme trypsin through its binding of Lys178 as an S1 residue in proteolysis suggests the full formation of L-Lys using the editing methods described here within detection limits. Sequence coverage of the region of the precursor PstS-Cys178 is compared with that following editing to PstS-Lys178; this leads to tryptic cleavage C-terminal to the edited Lys178. Accordingly, the results of the tandem LCMS analysis of the modified protein were processed using the protein sequence database in which wild type PstS was changed to PstS-Lys178. (**c**) The observation of full deacetylation of chemically-edited KAc18in histone H3-KAc18 by stereospecific HDAC enzyme Sirt2 to generate H3-Lys18 suggests the full formation of L-KAc using the editing methods described here within detection limits.

Finally, the ability to install Bal through dual initiation-trapping using B_2_Cat_2_ (see above) was used to further extend the range of methods that would allow access to native residues. Thus, when conducted in open air L-Ser was formed as a consequence of a three-step, one-pot initiation-trapping-migratory oxidation pathway as another example of a Cβ–Oγ bond formation. In this way the original L-Cys residue was mutated (via L-Fpc and, without isolation, L-Bal) to L-Ser (**Figure 4a,b** and **Supplementary Figure 11**)) in 80% conversion as a further example of the use of post-translational editing in allowing chemical protein mutagenesis.

### Functional and spectroscopic characterization of the retention of native L-stereochemistry

To probe that the configuration of the residues formed from on-protein radical trapping, three differing systems were tested with complementary and orthogonal spectroscopic and functional methods. First, intact protein ^19^F NMR experiments were conducted on a histone H3 variant (H3-Bal9) in which Bal had been installed at site 9 (**Figure 5a**). We have previously shown that chiral shift reagent **1** is able to distinguish epimers of D- and L-Bal following installation of epimeric D-/L-Bal into proteins.^58^ The resulting fluorinated diol boronate ester adducts formed through binding to the Bal residue on proteins therefore reveals D/L configuration information *via* ^19^F NMR analysis.^58^ When an epimeric sample of H3–D/L-Bal9 (formed using Cu(II)-catalyzed boronylation^58^) was mixed with chiral shift reagent **1**, a ratio of approximately 1:1 was detected after integration of the corresponding CF_3_ resonances (**Figure 5a**, **red line**) consistent with the known poor diastereoselectivity. However, when a sample of putative H3–L-Bal9, prepared using the methods described here, was mixed with chiral shift reagent **1**, only a single resonance was detected (**Figure 5a**, **blue line**). When these two protein samples were combined in equal amounts (mutually ‘spiked’) a ratio of >2:1 was detected after integration following enhancement of only one of the resonances in the H3–D/L-Bal9 epimeric mixture (**Figure 5a**, **black line**), consistent with the enhancement of concentration of only one epimer. Together these data strongly indicated that the native L-stereochemistry at the modified residue was preserved during the reaction to form H3–L-Bal9 from H3–L-Cys9 via H3–L-Fpc9.

Next, we tested the stereospecific enzymatic reaction of residues in two other protein scaffolds and sites. Trypsin digests stereoselectively at L–Lys (rather than D-Lys) residues,^65^ including in polypeptides,^66^ thereby enabling tryptic digests for so-called peptide mapping in protein sequencing via MSMS methods. When PstS-Lys178, that had been formed using the methods described here, was subject to digestion, full and complete enzymatic digestion to the corresponding octapeptide PstS^179–186^ peptidic fragment was observed (**Figure 5b**). This arises from cleavage C-terminal of Lys178 through the known selectivity of trypsin for basic P1 residues in its S1 pocket and was further confirmed to be L-Lys specific through the notable absence of the PstS^179–186^ peptide in all other digests of the other PstS-178 variants formed in this study to the lack of digestion at this site, which were instead formed as native PstS^176–186^ undecapeptides.

Finally, after exploiting the stereospecificity of a protein-backbone-modifying enzyme we then exploited the complementary stereospecificity of a protein-sidechain-modifying enzyme in yet another protein scaffold and site, H3–L-KAc18. Acetylated lysine (KAc) is known to be stereospecifically deacetylated during the ‘erasing’ of the acetyl post-translational modification at L-Lys18 in histone H3 by the histone deacetylation (HDAC) enzyme Sirt2.^67, 68^ We have previously shown that this enzyme does not fully process mixed D-/L-epimers of acetylated L-lysine or its analogues, also consistent with this L-stereospecificity and with failed deacetylation of acetylated D-KAc.^14, 20^ Strikingly, when H3-KAc18, prepared using the methods described here, was treated with Sirt2 then complete deacetylation was observed, consistent with the presence only of an acetylated L-KAc residue (**Figure 5c**).

Together this testing of configuration in three different, representative protein scaffold, sites and residue systems (H3–L-Bal9, PstS–L-Lys178 and H3–L-KAc18) suggested that L-configuration is well preserved through diverse reactions and substrates using the reactions we describe here in a stereoretentive manner.

## Discussion

In summary, we have described an efficient on-protein free radical generation-and-trapping method via light-mediated C_β_-S_γ_ bond cleavage to now realize a general form of chemical mutagenesis via post translational editing. Most importantly we observe that the L-configuration of the stereogenic C_α_ at mutated residues is preserved during such editing.

Not all of the problems of this form of stereoretentive post-translational editing have been solved and some limitations remain. For example, whilst many reactions are efficient in the trapping of the on-protein radicals that are formed, in other cases lower conversions are observed for some of the proof-of-principle systems that we describe here. Notably, however, the secondary side products in these cases are the directly reduced inactive L-alanine variants (see above for examples with Sirt2 or trypsin). In this way, this form of editing even with lower conversions still allows ready chemical installation of altered residues for rapid functional scoping of protein activity in the background of a typically inactive (i.e. L-Ala) variant. In addition, in the case of phenyl vinyl sulfone non-specific modification of protein lysines was observed (**Supplementary Figure 15**) in addition to on-protein radical trapping, highlighting that in some cases competent radical acceptors may also have concomitant additional direct electrophilic reactivity that proves competitive.

Whilst the full details of the mechanism of the reactions that we describe here are the subject of current experiments and we cannot discount other mechanisms, our initial data suggests that a pathway consistent with the formation of charge transfer complexes is followed involving so-called electron acceptor-electron donor (EDA) species. Indeed, the observation of a weak charge transfer (CT) band lower in energy than the parent molecular transitions was observed not only in model small molecule systems but also in protein systems during spectroscopic probing (via absorption spectra) of the pre-irradiation reaction mixture of bis(catecholato)diboron (B_2_Cat_2_ (electron donor, 0.1 mM) and PstS-Fpc178 (0.02 mM, in Tris pH 8.0) (**Supplementary Figures 12,13**).

The breadth of bond-forming reactions at diverse sites and in differing representative protein scaffolds used here suggest a true generality for this L-residue specific method that could enable numerous applications in protein science. To pick just three, the installation of L-Bal as a minimal homochiral precursor boronic acid may open yet further synthetic avenues. Now that such L-Bal boronic acids can be installed into proteins with site selectivity, then it may now be possible to take advantage of well-developed boronic acid transformations for further post-translational editing. Moreover, the functional utility that we demonstrate here now of an on-protein L-alanyl radical, without the need for additional stabilization,^14^ opens the door to further radical processing methods including metal-relayed trapping and so-called ‘sorting’^69^ methods with further potential for editing of biological systems. It has also not escaped our attention that this strategy should readily enable complementary methods for the creation of epimeric series not only of L-residues but also D-residues in proteins and studies in this direction will be published in due course.

## Acknowledgments

The authors would like to thank Tim Mollner for invaluable discussions.

The authors are grateful to Nick Devoogdt for the provision of the cAbVCAM plasmid and Guy Salvesen for plasmids distributed via Addgene.

The Next Generation Chemistry theme at the Franklin Institute is supported by the EPSRC (V011359/1 (P)).

