## Supplementary Information for "Stereoretentive Post-Translational Protein Editing"

### Supplementary Figures

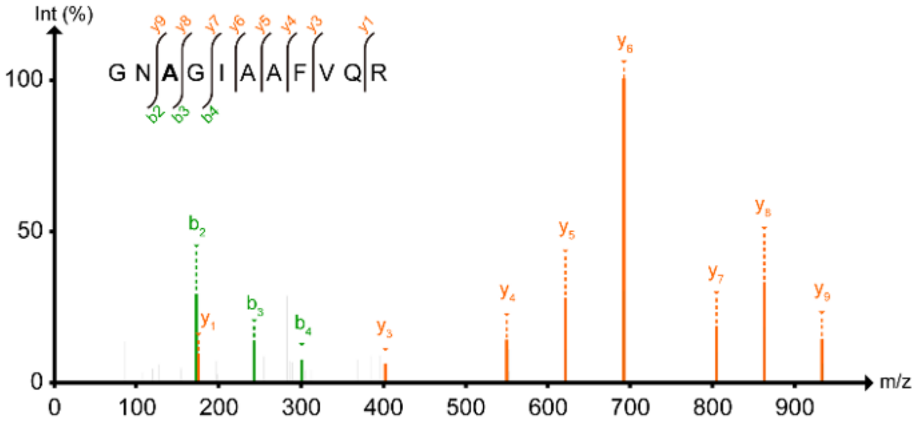

**Supplementary Figure 1: MS/MS analysis of PstS-Ala178**

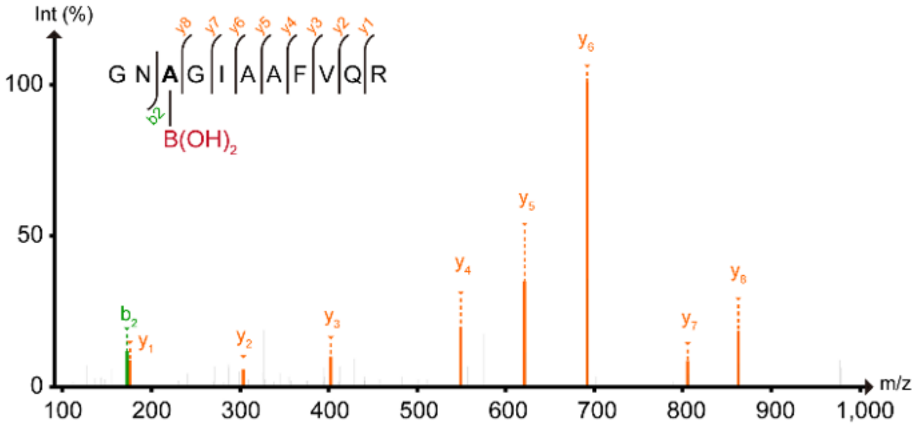

**Supplementary Figure 2: MS/MS analysis of PstS-Bal178**

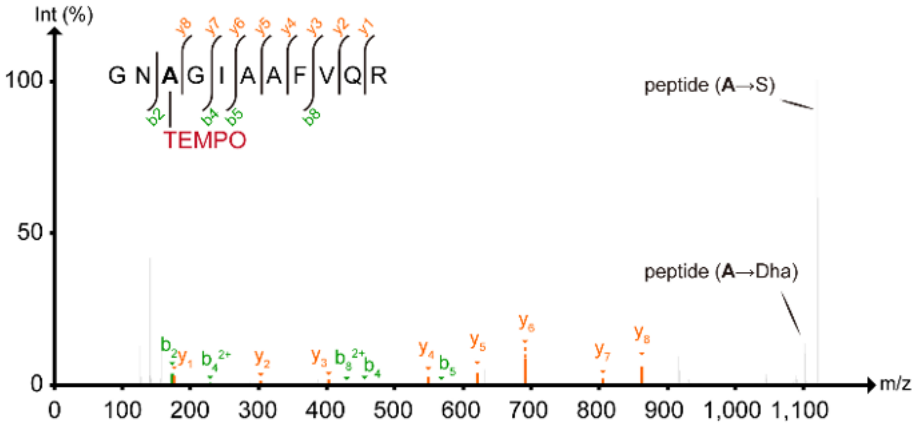

**Supplementary Figure 3: MS/MS analysis of PstS-TEMPO-A178**

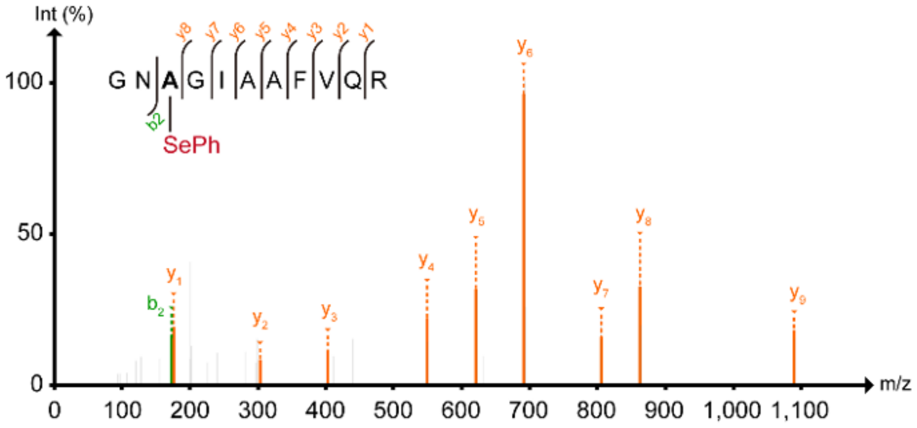

**Supplementary Figure 4: MS/MS analysis of PstS-SecPh178**

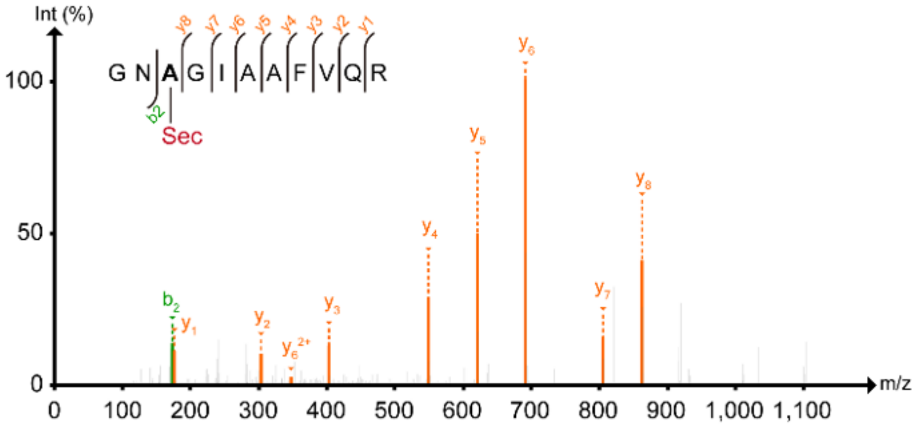

**Supplementary Figure 5: MS/MS analysis of PstS-Sel178**

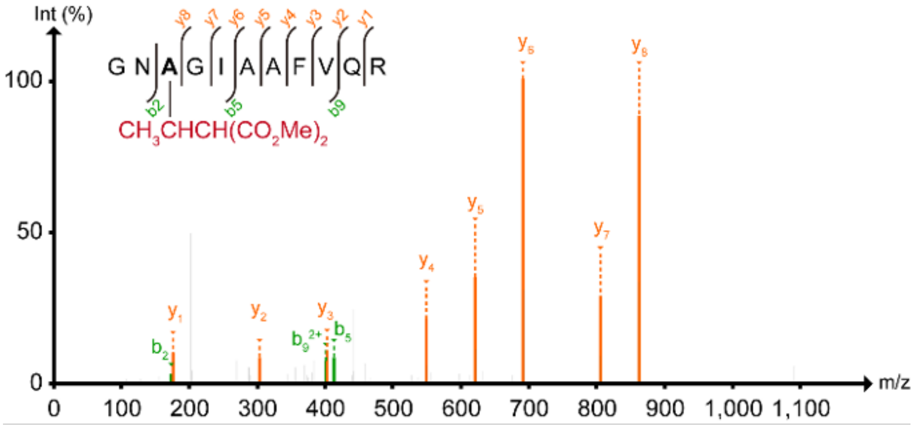

**Supplementary Figure 6: MS/MS analysis of PstS-Mal-L178**

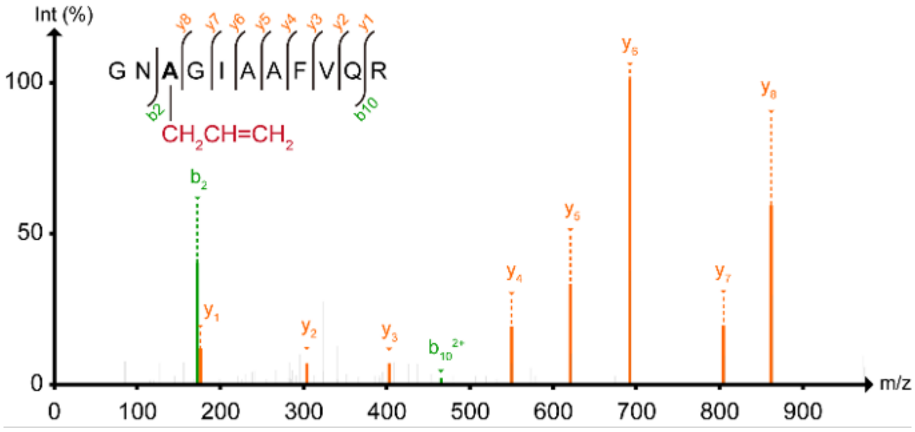

**Supplementary Figure 7: MS/MS analysis of PstS-Hag178**

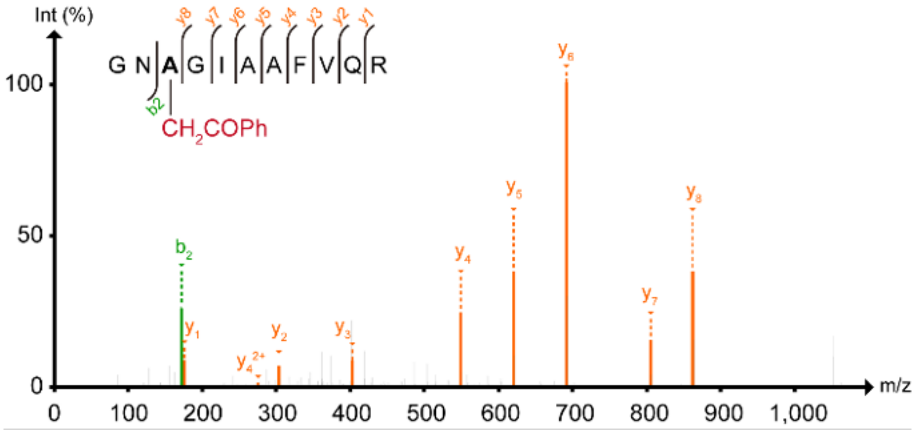

**Supplementary Figure 8: MS/MS analysis of PstS-A(Ketone)178**

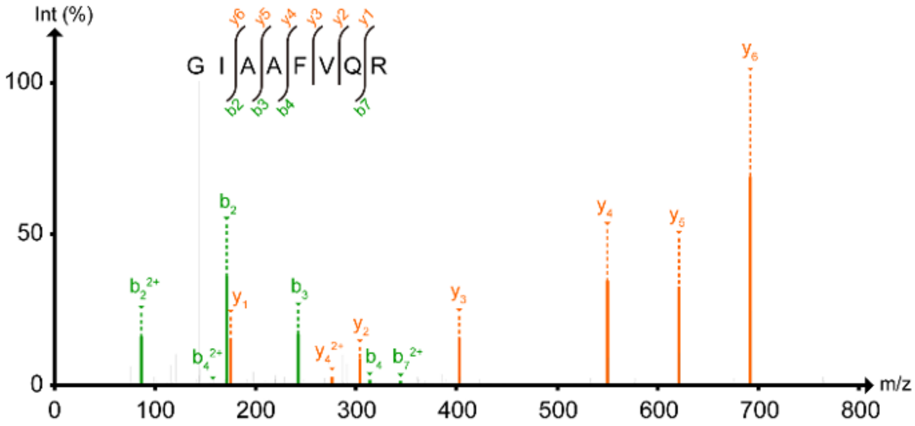

**Supplementary Figure 9: MS/MS analysis of PstS-Lys178**

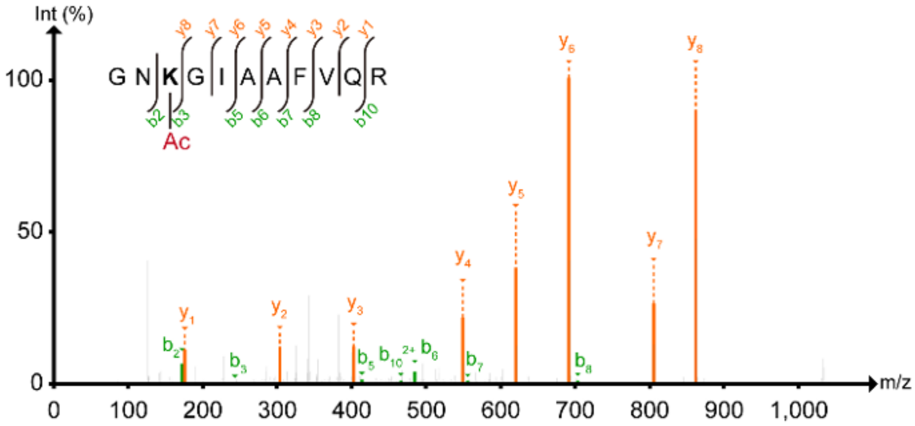

**Supplementary Figure 10: MS/MS analysis of PstS-KAc178**

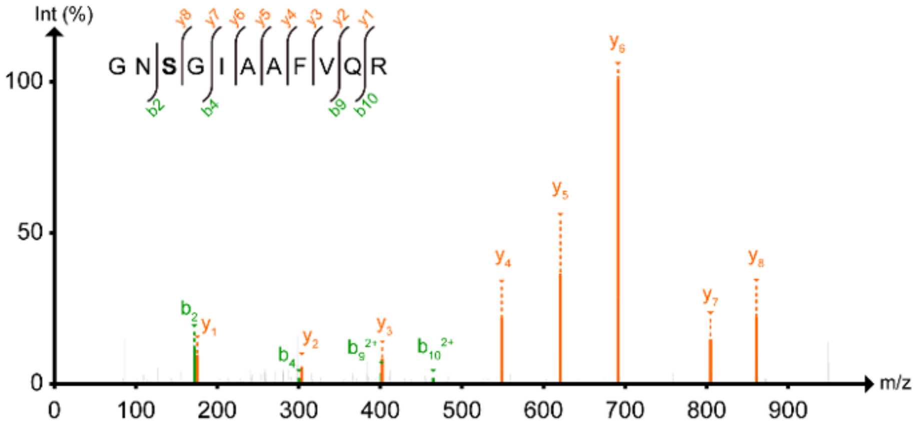

**Supplementary Figure 11: MS/MS analysis of Psts-Ser178**

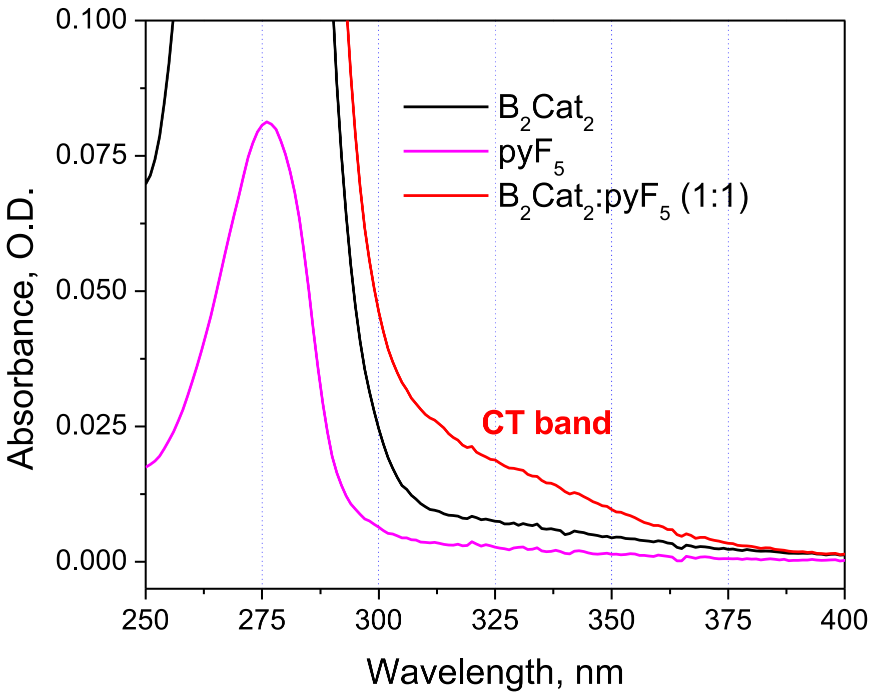

**Supplementary Figure 12: Spectral signature of EDA complexation**

Absorption spectra of bis(catecholato)diboron ($B_2Cat_2$, electron donor, 0.1 mM, in black) and pentafluoropyridine ($pyF_5$, electron acceptor, 0.1 mM, in magenta). The red trace represents 1:1 mixture of donor and acceptor in buffer solution (Tris, pH = 8.0). A weak charge transfer (CT) band lower in energy than the parent molecular transitions can be observed, which confirms EDA complexation.

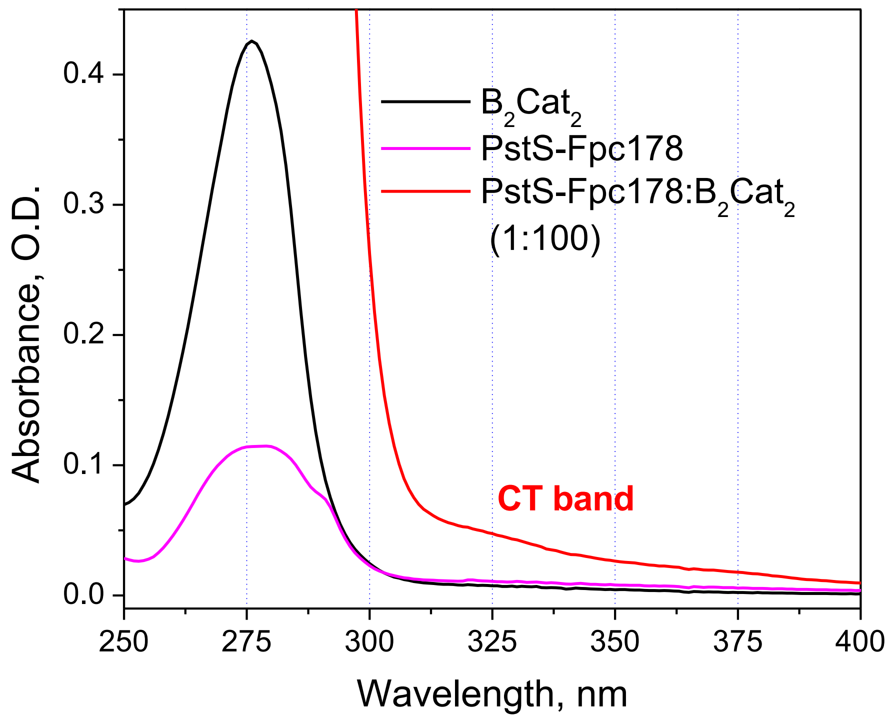

**Supplementary Figure 13: Spectral signature of EDA complexation on protein**

Absorption spectra of bis(catecholato)diboron ($B_2Cat_2$, electron donor, 0.1 mM, in black) and PstS-Fpc178 (protein with electron acceptor, 0.02 mM, in magenta). The red trace represents 1:100 mixture of donor and acceptor in buffer solution (Tris, pH = 8.0). A weak charge transfer (CT) band lower in energy than the parent molecular transitions can be observed, which confirms EDA complexation.

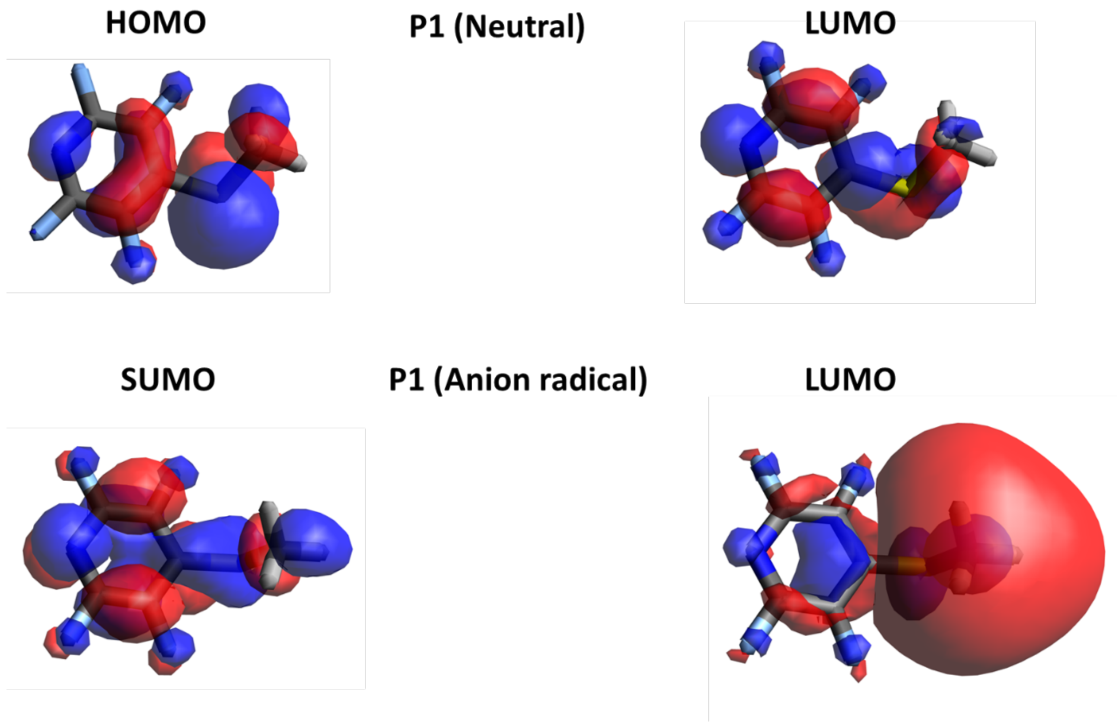

**Supplementary Figure 14: Frontier molecular orbitals from DFT Calculations**

The frontier molecular orbitals of P1 molecule is shown in its neutral and anion radical state. Molecular orbitals shown here are obtained by performing DFT calculations using B3LYP/6311G(d,p).

(a)

AAWKTNIK
DQKKPEQGTEVLK
DSSGKPLY
ETGNKVNYQGIGSSGGVK
FFDWAYKTGAK
GADWSKTFAQDLTNQK
GEDAWPITSTTFILIHK
GNA(178)GIAAFVQR
IKKWDDEAIAK
KPEQGTEVLK
KWDDEAIAK
LISADGKPVSPTEENFANAAK
LISADGKPVSPTEENFANAAKGADWSK

LNPGLKLPSQNIAVVR
LPGAIGYVEYAYAKQNNLAYTK
NNVGTGSTVKWPIGLGGK
QNNLAYTKLISADGKPVSPTEENFANAAK
SGELVLDGKTLGDIYLGK
TGAKQANDLDYASLPDSVVEQVR
TLGDIYLGKIK
TNIKDSSGKPLY
VNEEWKNNVGTGSTVK
VNEEWKNNVGTGSTVKWPIGLGGK
WADTYQKETGNK
WDDEAIAKLNPGLK

(b)

MEASLTGAGATFPAPVYAKWADTYQKETGNKVNYQGIGSSGGVKQII
ANTVDFGASDAPLSDEKLAQEGLFQFPTVIGGVVLAVNIPGLKSGELV
LDGKTLGDIYLGKIKKWDDEAIAKLNPGLKLPSQNIAVVRRADGSGTS
FVFTSYLAKVNEEWKNNVGTGSTVKWPIGLGGKGNAGIAAFVQRLP
GAIGYVEYAYAKQNNLAYTKLISADGKPVSPTEENFANAAKGADWSK
TFAQDLTNQKGEDAWPITSTTFILIHKDQKKPEQGTEVLKFFDWAYKT
GAKQANDLDYASLPDSVVEQVRAAWKTNIKDSSGKPLY

(c)

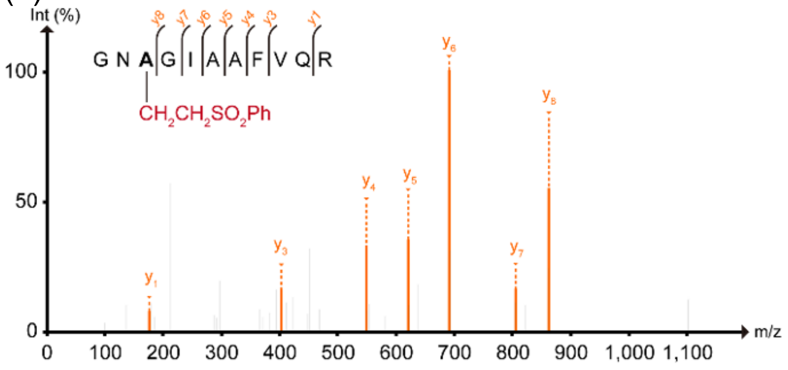

**Supplementary Figure 15: MS/MS analysis of Psts-A(Sulphone)178**

(**a**) A list of peptides with the mass shift corresponding (within the mass accuracy of 0.005 Da) to $CH_2CH_2SO_2Ph$ identified in the tryptic digest of PstS-A(Sulfone)178. Possible additional non-specific modification sites are labelled with the red colour. The modification of the lysine residues in these peptides led to missed cleavages by trypsin at these sites. (**b**) Coverage of the amino-acid sequence of PstS by peptides listed in (a). Modification sites are highlighted in red. (**c**) MS/MS spectrum for PstS$^{176\text{-}186}$ peptide in PstS-A(Sulfone)178.

### Supplementary Tables

**Supplementary Table 1: Optimisation of chemical introduction of tetrafluoropyridyl-cysteine (Fpc) into proteins**

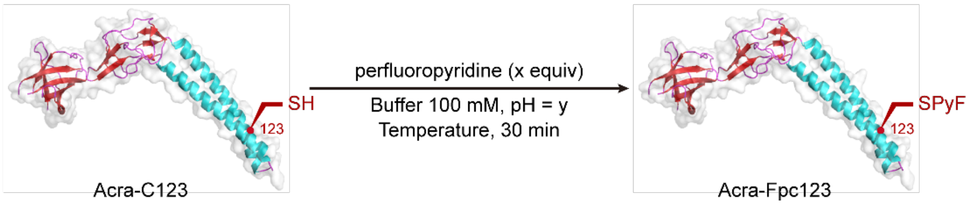

| entry | Buffer | x | y | Temperature (°C) | Conversion |
|---|---|---|---|---|---|
| 1 | NaPi | 100 | 8.0 | 4 | 75% |
| 2 | NaPi | 200 | 8.0 | 4 | 84% |
| 3 | NaPi | 400 | 8.0 | 4 | 90% |
| 4 | NaPi | 200 | 8.0 | 25 | >98% conversion |
| 5 | NaPi | 200 | 7.4 | 25 | >98% conversion |
| 6 | NaPi | 200 | 7.0 | 25 | 60% |
| 7 | NaPi | 200 | 6.0 | 25 | Trace |
| 8 | Tricine | 200 | 7.4 | 25 | >98% conversion |
| 9 | Tris | 200 | 7.4 | 25 | >98% conversion |

General procedure: AcrA-Cys123 (20 µM, 50 µL, in buffer), perfluoropyridine (x equiv, 100 mM, in DMSO), were mixed and incubated at 4 °C or 25 °C for 30 min. The reaction mixture was then analysed by LC-MS.

**Supplementary Table 2: Optimisation of light-mediated $C_\beta$-$S_\gamma$ bond cleavage**

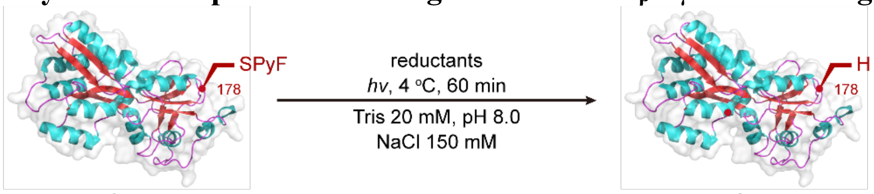

PstS-Fpc178 PstS-A178

| Entry | Reductants | hυ | Conversion |
|---|---|---|---|
| 1 | 4-Me-PhSH (100 equiv) | 365 nm | >98% conversion |
| 2 | 4-Me-PhSH (100 equiv) | 385 nm | 0% |
| 3 | 4-Me-PhSH (100 equiv) | 405 nm | 0% |
| 4 | 4-Me-PhSH (100 equiv) | 420 nm | 0% |
| 5 | 4-Me-PhSH (100 equiv) | 445 nm | 0% |
| 6 | 2-Cl-4-F-PhSH (100 equiv) | 365 nm | >98% conversion |
| 7 | Ir(dtppy)(bpy)$_2$BF$_4$ (10 equiv), FeSO$_4$ (200 equiv) | 365 nm | 90% |
| 8 | Ir(dtppy)(bpy)$_2$BF$_4$ (10 equiv), FeSO$_4$ (200 equiv) | 385 nm | 90% |
| 9 | Ir(dtppy)(bpy)$_2$BF$_4$ (10 equiv), FeSO$_4$ (200 equiv) | 405 nm | 71% |
| 10 | Ir(dtppy)(bpy)$_2$BF$_4$ (10 equiv), FeSO$_4$ (200 equiv) | 420 nm | 77% |
| 11 | Ir(dtppy)(bpy)$_2$BF$_4$ (10 equiv), FeSO$_4$ (200 equiv) | 445 nm | 35% |
| 12 | Ru(bpy)$_3$Cl$_2$ (10 equiv), FeSO$_4$ (200 equiv) | 365 nm | 52% |
| 13 | Ru(bpy)$_3$Cl$_2$ (10 equiv), FeSO$_4$ (200 equiv) | 385 nm | 63% |
| 14 | Ru(bpy)$_3$Cl$_2$ (10 equiv), FeSO$_4$ (200 equiv) | 405 nm | 66% |
| 15 | Ru(bpy)$_3$Cl$_2$ (10 equiv), FeSO$_4$ (200 equiv) | 420 nm | 75% |
| 16 | Ru(bpy)$_3$Cl$_2$ (10 equiv), FeSO$_4$ (200 equiv) | 445 nm | 51% |
| 17 | B$_2$Cat$_2$ (100 equiv) | 365 nm | >98% conversion[a] |
| 18 | B$_2$Cat$_2$ (100 equiv) | 385 nm | 0% |
| 19 | B$_2$Cat$_2$ (100 equiv) | 405 nm | 0% |

| | | | |
|---|---|---|---|
| 20 | $B_2Cat_2$ (100 equiv) | 420 nm | 0% |
| 21 | $B_2Cat_2$ (100 equiv) | 445 nm | 0% |

In glovebox PstS-Fpc178 (15 µM, 50 µL), reductants, were mixed and irradiated with *hυ* at 4 $^{o}$C for 60 min. The reaction mixture was then analysed by LC-MS.

[a] 9% conversion to PstS-A178; 91% conversion to PstS-Bal178.

### Supplementary Methods

#### General Experimental Procedures

Chemicals and solvents were purchased from Sigma-Aldrich UK, Acros UK, Alfa Aesar UK, Carbosynth, Fluorochem or Fischer UK and were used as delivered unless stated otherwise. Thin layer chromatography (TLC) was carried out using plastic 0.20 mm Polygram® Silg/UV254 plates that were dried using a heat gun and visualised under UV (λmax 254 nm or 366 nm) or by use of anisaldehyde, potassium permanganate, sulfuric acid or vanillin dip. Flash column chromatography was performed using Geduran® Si 60.8.2 or a Teldyne Flash Purification System with either Kinesis Telos or Biotage Snap columns.

NMR Spectroscopy General: Deuterated solvents were used as the lock and the residual protonated solvent as the internal reference peak. Spectra were analysed using MestReNova.

Glovebox Usage: Anaerobic atmosphere was achieved using a Belle Technology glovebox equipped with the BASF R3-11G catalyst. The oxygen level was measured below 6 ppm during all reactions.

Photobox usage: Reactions were performed on the Zinsser Analytic off-deck irradiation system with two reaction positions irradiated by Lumidox II 96-LED arrays. 96-position, open-bottom Desyre reaction blocks equipped with several 1.2 mL vials were placed in the irradiation box, agitated at 400 rpm, and irradiated with 365 nm light at 230 mW per well for 60 mins. Cooling of reaction positions to 4 °C was provided by an off-deck circulating cooler.

Raw data is deposited at doi: 10.5281/zenodo.7011026

#### Protein Mass Spectrometry

Protein samples were analysed on Waters Xevo G2-XS QTof mass spectrometers equipped with a Waters Acquity UPLC. Separation was achieved using a Thermo Scientific ProSwift RP-2H monolithic column (4.6 mm × 50 mm) using water + 0.1% formic acid (solvent A) and acetonitrile + 0.1% formic acid (solvent B) as mobile phase at a flow rate of 0.3 ml/min and running a 10-min linear gradient as follows: 5% solvent B for 1 min, 5 to 95% solvent B over 6 min, 95 to 5% solvent B over 1 min, and 5% solvent B for 2 min. Spectra were deconvoluted using MassLynx 4.1 (Waters) and the “MaxEnt1” deconvolution algorithm with the following settings: resolution: 1.0 Da per channel; damage model: uniform Gaussian; width at half height: 0.4 Da; minimum intensity ratios: 33% (left) and 33% (right); and iterate to convergence. Con-versions were calculated from peak intensities.

**Protein digestion and analysis by MS/MS**

For in-solution proteolytic digestion, samples were dissolved in 100 mM ammonium bicarbonate, reduced with 10 mM tris(2-carboxyethyl)phosphine (Thermo Fisher) at 56 °C for 30 min and alkylated with 30 mM 2-chloroacetamide (Sigma Aldrich) at room temperature for 30 min in the dark. Trypsin (Pierce) was added to each sample for an overnight incubation at 37 °C with 1:25 trypsin:protein ratio (w/w). The samples were desalted by Oasis HLB cartridges (Waters), dried, and reconstituted in water containing 5% formic acid, 5% DMSO right before the LC-MS analysis.

For in-gel proteolytic digestion (PstS-Bal178 and PstS-TEMPO-A178), samples were resolved via SDS-PAGE and stained with InstantBlue® (Abcam). After destaining in MilliQ® $H_2O$ for 20 min, the corresponding bands were excised and cut into small cubes. The gels were extensively destained twice with 50 mM $NH_4HCO_3$/MeCN (1:1, v/v) at room temperature for 10 min. After washing with 50 mM $NH_4HCO_3$ for 10 min, samples were reduced with 10 mM tris(2-carboxyethyl)phosphine (Thermo Fisher) at room temperature for 60 min and alkylated with 30 mM 2-chloroacetamide (Sigma Aldrich) at room temperature for 30 min in the dark. The gels were washed twice with 50 mM $NH_4HCO_3$/MeCN (1:1, v/v) at room temperature for 10 min. In turn, the gels were dehydrated with MeCN and dried under air for 10 min before overnight trypsin (Pierce) digestion in 50 mM $NH_4HCO_3$ at 37 °C with 1:25 trypsin:protein ratio (w/w).

Samples were subjected to LC-MS/MS using a UltiMate 3000 nanoUHPLC system (Thermo Fisher Scientific) coupled to an Orbitrap Fusion Lumos (Thermo Fisher Scientific). The peptides were trapped on a C18 PepMap100 pre-column (300 µm i.d. x 5 mm, 100 Å, Thermo Fisher Scientific) using solvent A (0.1% formic acid in water), then separated on an in-house packed analytical column (75 µm i.d. x 50 cm in-house packed with ReproSil Gold 120 C18, 1.9 µm, Dr. Maisch GmbH) with a gradient of 12% to 40% B (0.1% formic acid in acetonitrile) over 15 min at a flow rate of 200 nL/min. Full scan MS spectra were acquired in the Orbitrap (scan range 350-1400 m/z, resolution 60000, AGC target 1200000). The 20 most intense peaks were selected for HCD fragmentation at 30% of normalised collision energy and with a resolution 7500, AGC target 20000.

Spectra were searched using FragPipe (v18.0) MSFragger 3.5[1] with standard 'open' search settings against database (PDB ID 2abh with mutation D178A). Data was filtered using the inbuilt tools within FragPipe to an FDR of below 1%. Modified peptides were discerned by filtering the resulting dataset using the expected changes in mass caused by each modification.

**General procedure 1: Cys-tetrafluoropyridylsulfide (Fpc) formation in proteins**

To a solution of Protein (1.0 mL, 1 mg/mL, 1.00 equiv) NaPi buffer (100 mM, pH 7.4)[a] was added prepared stock solutions of perfluoropyridine (1 M in DMSO, 100 equiv). The mixture was shaken for 30 min at room temperature. The protein was desalted by passing through a GE MiniTrap G-25 column pre-equilibrated with Tris buffer (20 mM, NaCl 150 mM, pH 8.0) according to the manufacturer's instructions.

[a]For protein PstS-D57C, PstS-D103C, PstS-A197C, NaPi buffer (100 mM, 3 M Gdn·HCl, pH 7.4) was used; for histone proteins, tricine buffer (100 mM, 3 M Gdn·HCl, pH 7.4) was used.

**General procedure 2: Alanyl radical trapped by HAT**

In glovebox PstS-Fpc178 (15 μM, 50 μL) in Tris buffer (20 mM, NaCl 150 mM, pH 8.0), 4-Me-PhSH (50 mM in DMSO, 100 equiv, 1.5 μL) were mixed and irradiated with 365 nm light at 4 °C for 60 min. The reaction mixture was then analysed by LC-MS.

**General procedure 3: Alanyl radical trapped by TEMPO**

In glovebox PstS-Fpc178 (15 μM, 50 μL) in Tris buffer (20 mM, NaCl 150 mM, pH 8.0), 4-Me-PhSH (50 mM in DMSO, 100 equiv, 1.5 μL), TEMPO (100 mM in DMSO, 200 equiv, 1.5 μL) were mixed and irradiated with 365 nm light at 4 °C for 60 min. The reaction mixture was then analysed by LC-MS.

**General procedure 4: Alanyl radical trapped by diselenides**

In glovebox PstS-Fpc178 (15 μM, 50 μL) in Tris buffer (20 mM, NaCl 150 mM, pH 8.0), 2,6-di-Cl-PhSH[b] (50 mM in DMSO, 100 equiv, 1.5 μL), diselenides (100 mM in DMSO, 200 equiv, 1.5 μL) were mixed and irradiated with 365 nm light at 4 °C for 60 min. The reaction mixture was then analysed by LC-MS.

[b]For PhSeSePh, 2,6-di-Cl-PhSH (50 mM in DMSO, 200 equiv, 3.0 equiv) was used.

**General procedure 5: Alanyl radical trapped by alkenes**

In glovebox PstS-Fpc178 (15 μM, 50 μL) in Tris buffer (20 mM, NaCl 150 mM, pH 8.0), 2-Cl-6-F-PhSH (50 mM in DMSO, 100 equiv, 1.5 μL), alkenes[c] (100 mM in DMSO, 200 equiv,

1.5 μL) were mixed and irradiated with 365 nm light at 4 °C for 60 min. The reaction mixture was then analysed by LC-MS.

[c]For allylic amine and 1-phenyl-1-trimethylsiloxyethylene, 2,6-di-Cl-PhSH (50 equiv) was used. For allylic phenyl sulfone (50 mM in DMSO, 100 equiv, 1.5 μL), 2-Cl-6-F-PhSH (50 mM in DMSO, 200 equiv, 3.0 μL) was used.

**General procedure 6: Alanyl radical trapped by $B_2Cat_2$.**

In glovebox PstS-Fpc178 (15 μM, 50 μL) in Tris buffer (20 mM, NaCl 150 mM, pH 8.0), $B_2Cat_2$ (100 mM in $H_2O$, 1000 equiv, 7.5 μL), were mixed and irradiated with 365 nm light at 4 °C for 60 min. The reaction mixture was then analysed by LC-MS.

**General procedure 7: Alanyl radical trapped by $O_2$.**

In air PstS-Fpc178 (15 μM, 50 μL) in Tris buffer (20 mM, NaCl 150 mM, pH 8.0), $B_2Cat_2$ (100 mM in $H_2O$, 1000 equiv, 7.5 μL), were mixed and irradiated with 365 nm light at 4 °C for 60 min. The reaction mixture was then analysed by LC-MS.

**General procedure 8: Chemical introduction of boronoalanine (Bal) into proteins.**

Cys-tetrafluoropyridylsulfide containing proteins were prepared *via* general procedure 1 without desalting. The mixture was directly used for the following borylation. In glovebox Cys-tetrafluoropyridylsulfide containing proteins (15 μM, 50 μL) in NaPi buffer (100 mM, pH 7.4), $B_2Cat_2$ (100 mM in $H_2O$, 1000 equiv, 7.5 μL), were mixed and irradiated with 365 nm light at 4 °C for 60 min. The reaction mixture was then analysed by LC-MS.

**Histone H3-Fpc2, -Fpc4, -Fpc9, -Fpc18, -Fpc27, Histone H4-Fpc20: protein expression, purification and Fpc generation**

Histone proteins expression and purification

Histone proteins were expressed and purified following a previously published procedure.[2]

**Histone H3-K4C**

ARTCQTARKSTGGKAPRKQLATKAARKSAPATGGVKKPHRYRPGTVALREIRRYQK STELLIRKLPFQRLVREIAQDFKTDLRFQSSAVMALQEASEAYLVALFEDTNLAAIHA KRVTIM PKDIQLARRIRGERA

Calculated mass = 15214

Observed mass = 15214

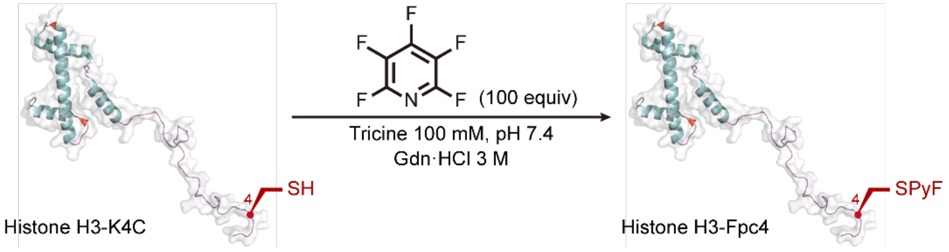

The Cys-tetrafluoropyridylsulfide-containing histone H3-Fpc4 was prepared according to general procedure 1.

ESI-MS spectrum for the modified histone is shown below.

Calculated mass = 15363

Observed mass = 15363

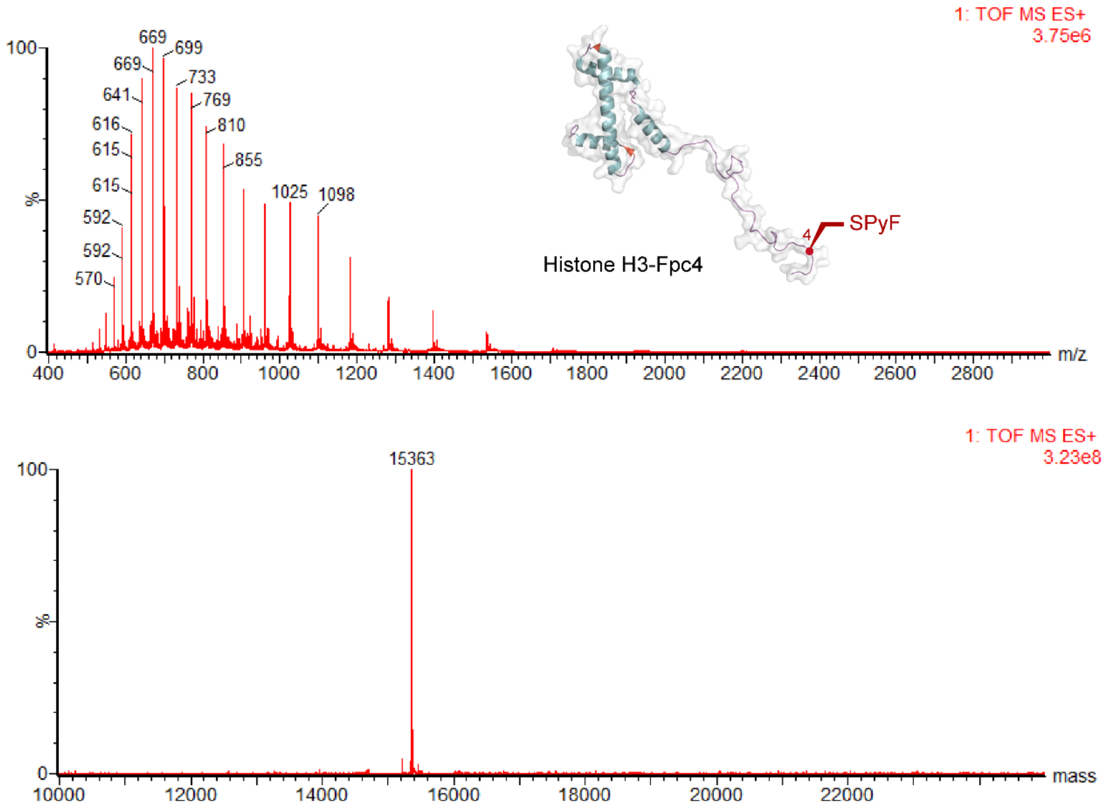

**Histone H3-K9C**

ARTKQTARCSTGGKAPRKQLATKAARKSAPATGGVKKPHRYRPGTVALREIRRYQK STELLIRKLPFQRLVREIAQDFKTDLRFQSSAVMALQEASEAYLVALFEDTNLAAIHA KRVTIM PKDIQLARRIRGERA

Calculated mass = 15214

Observed mass = 15214

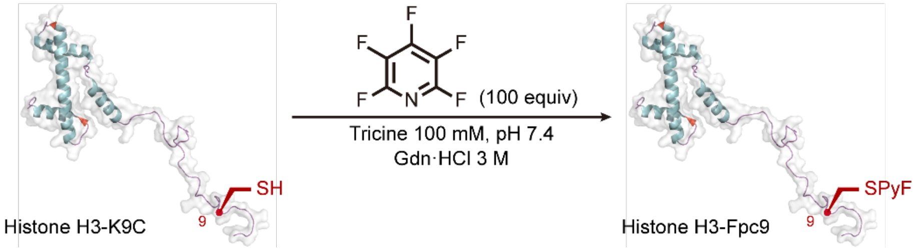

The Cys-tetrafluoropyridylsulfide-containing histone H3-Fpc9 was prepared according to general procedure 1.

ESI-MS spectrum for the modified histone is shown below.

Calculated mass = 15363

Observed mass = 15363

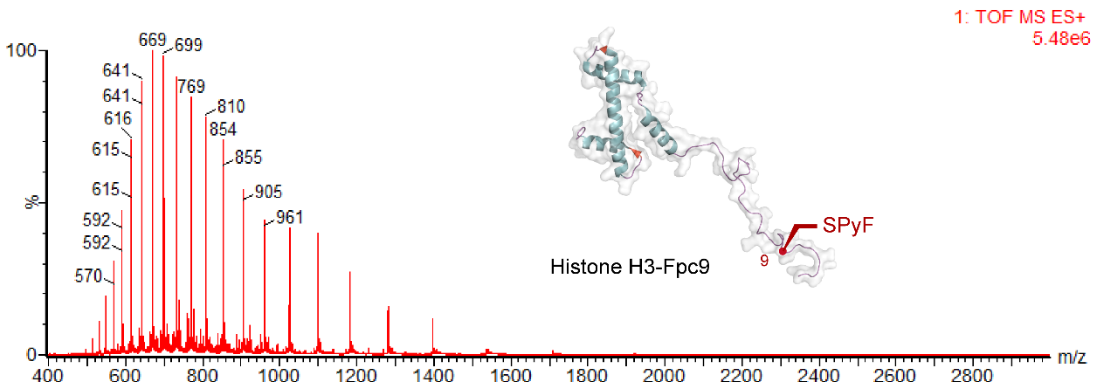

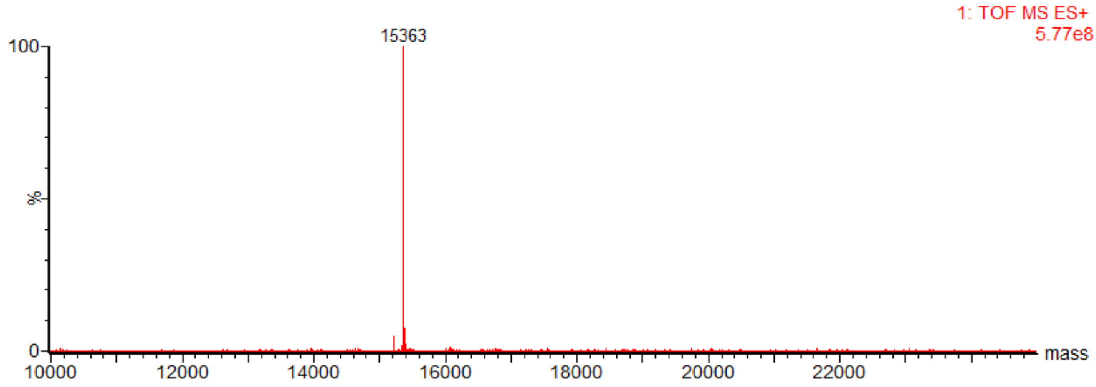

**Histone H3-K18C**

ARTKQTARKSTGGKAPRCQLATKAARKSAPATGGVKKPHRYRPGTVALREIRRYQK STELLIRKLPFQRLVREIAQDFKTDLRFQSSAVMALQEASEAYLVALFEDTNLAAIHA KRVTIM PKDIQLARRIRGERA

Calculated mass = 15214

Observed mass = 15214

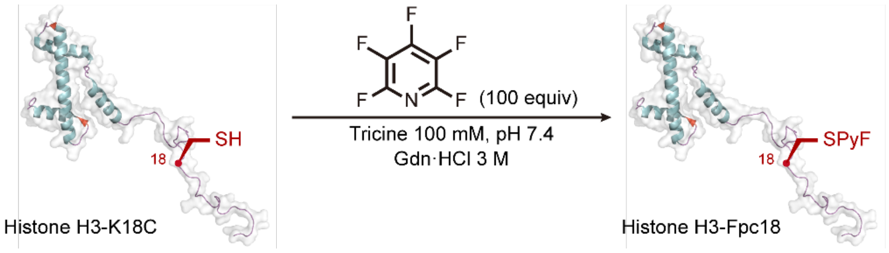

The Cys-tetrafluoropyridylsulfide-containing histone H3-Fpc18 was prepared according to general procedure 1.

ESI-MS spectrum for the modified histone is shown below.

Calculated mass = 15363

Observed mass = 15364

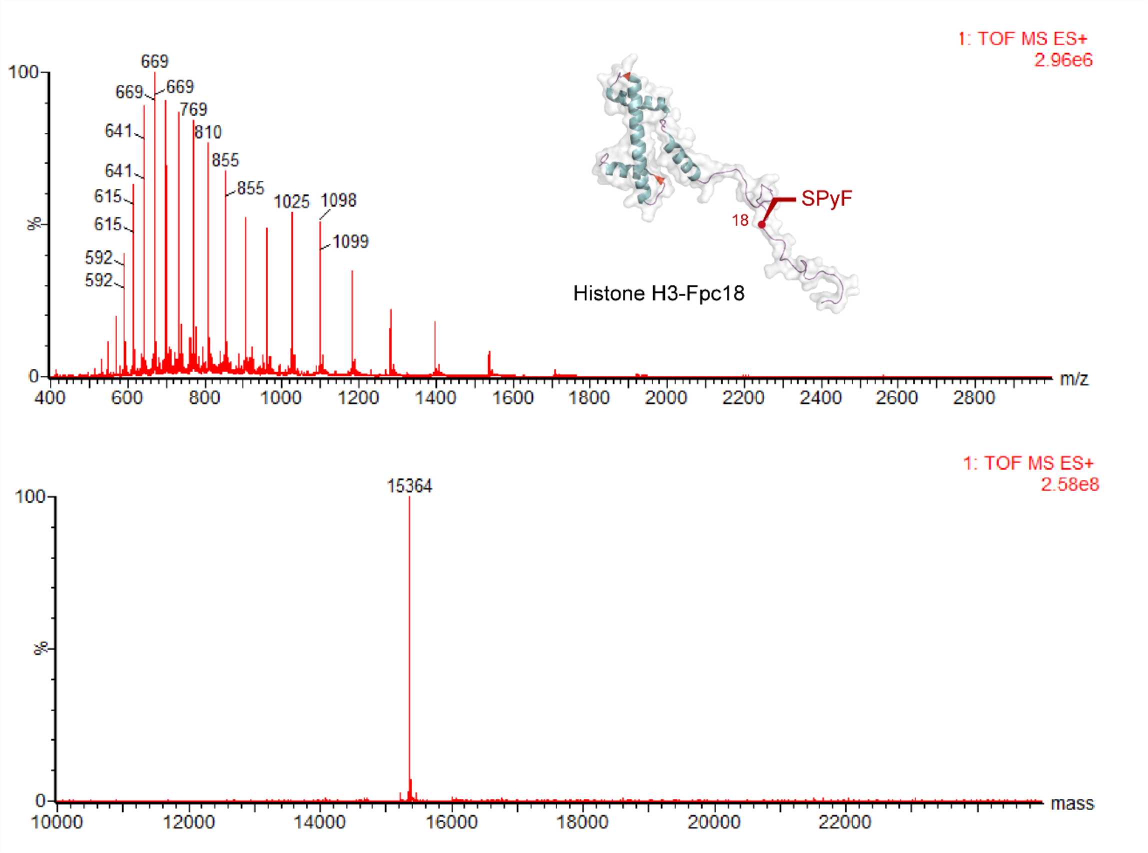

**Histone H3-K27C**

ARTKQTARKSTGGKAPRKQLATKAARCSAPATGGVKKPHRYRPGTVALREIRRYQK STELLIRKLPFQRLVREIAQDFKTDLRFQSSAVMALQEASEAYLVALFEDTNLAAIHA KRVTIM PKDIQLARRIRGERA

Calculated mass = 15214

Observed mass = 15214

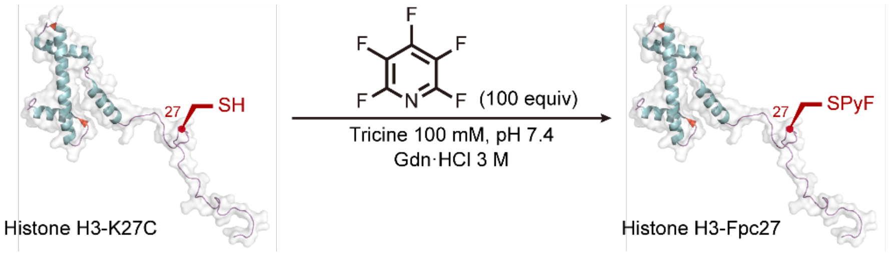

The Cys-tetrafluoropyridylsulfide-containing histone H3-Fpc27 was prepared according to general procedure 1.

ESI-MS spectrum for the modified histone is shown below.

Calculated mass = 15363

Observed mass = 15363

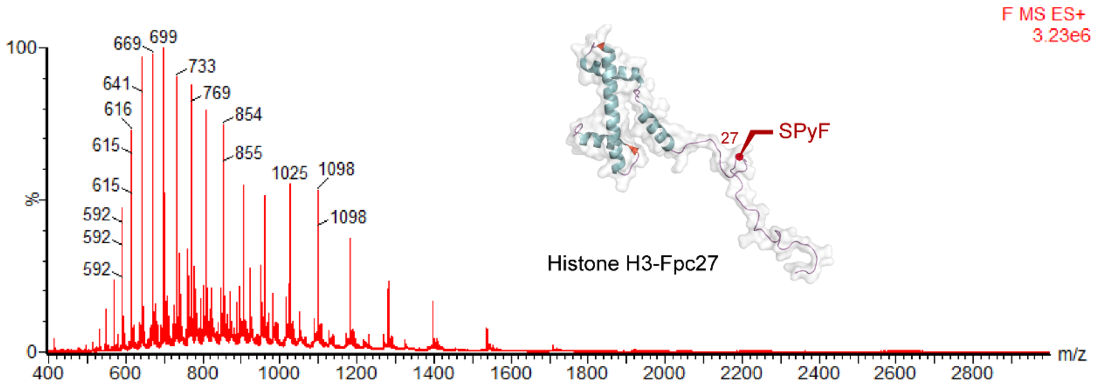

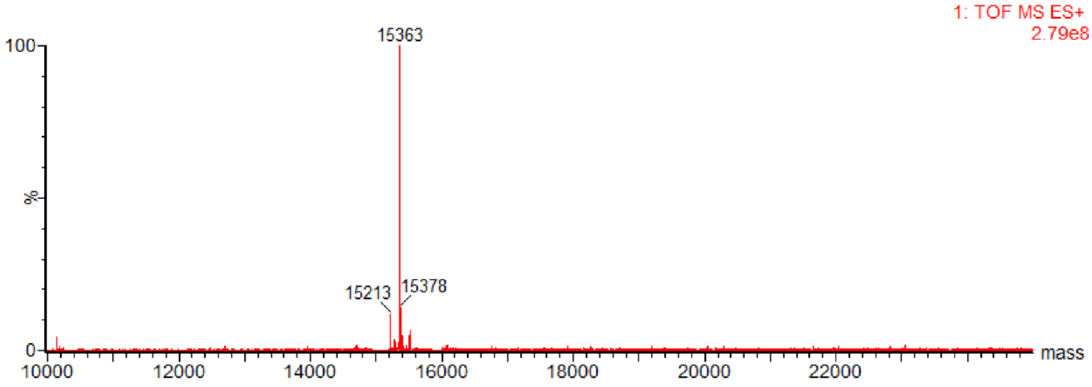

**Histone H3-R2C**

ACENLYFQGTKQTARKSTGGKAPRKQLATKAARKSAPATGGVKKPHRYRPGTVAL
REIRRYQKSTELLIRKLPFQRLVREIAQDFKTDLRFQSSAVMALQEASEAYLVALFED
TNLAAIHAKRVTIMPKDIQLARRIRGERA

Calculated mass = 16038

Observed mass = 16038

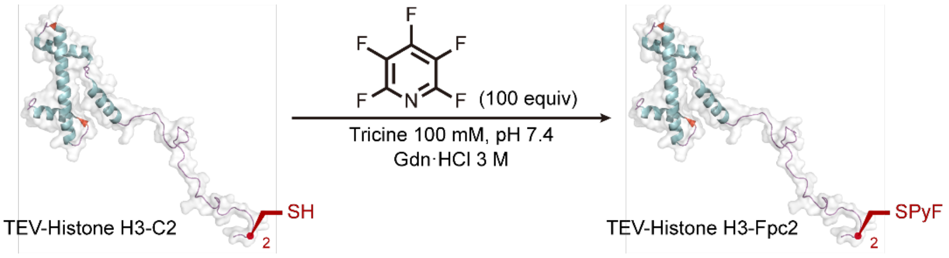

The Cys-tetrafluoropyridylsulfide-containing TEV-Histone H3-Fpc2 was prepared according to general procedure 1.

ESI-MS spectrum for the modified histone is shown below.

Calculated mass = 16186

Observed mass = 16187

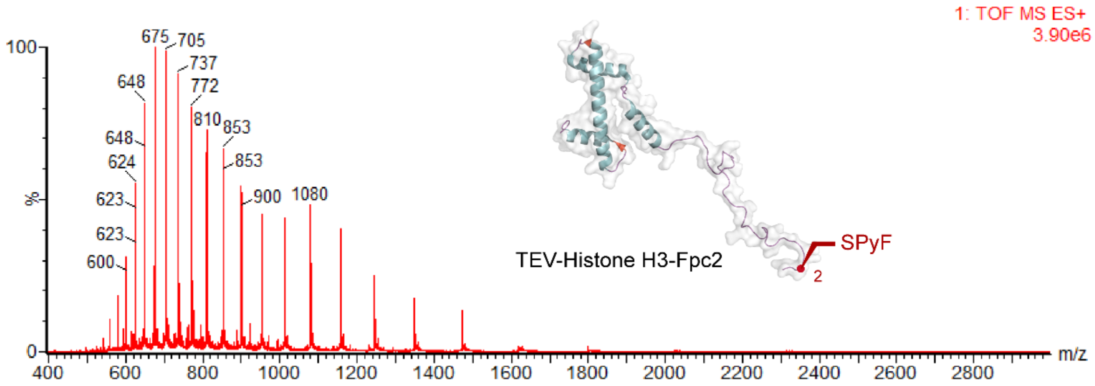

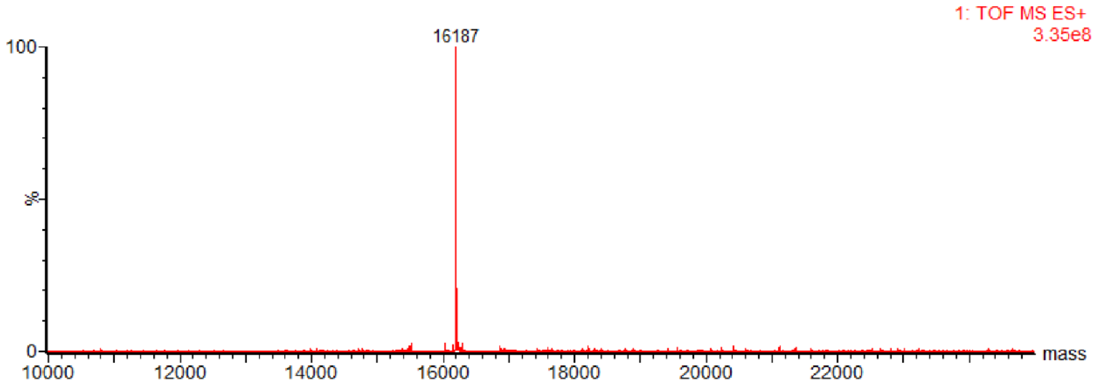

**H4-Fpc20: protein expression, purification and Fpc generation**

SGRGKGGKGLGKGGAKRHRCVLRDNIQGITKPAIRRLARRGGVKRISGLIYEETRGV LKVFLENVIRDAVTYTEHAKRKTVTAMDVVYALKRQGRTLYGFGG

Calculated mass = 11211;

Observed mass = 11212

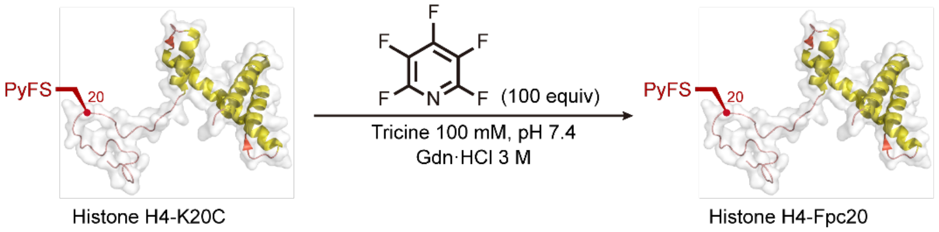

The Cys-tetrafluoropyridylsulfide-containing histone H4-Fpc20 was prepared according to general procedure 1.

ESI-MS spectrum for the modified histone is shown below.

Calculated mass = 11360;

Observed mass = 11360

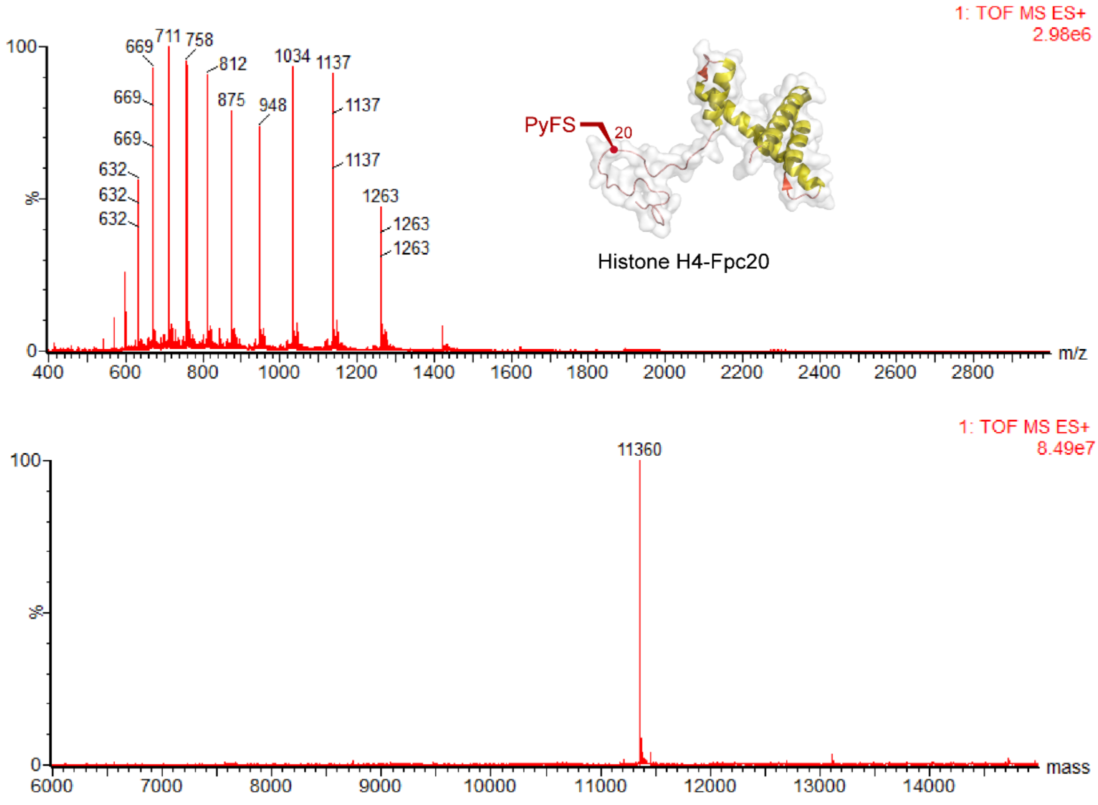

**PstS -Fpc57, -Fpc103, -Fpc178, -Fpc197: protein expression, purification and Fpc generation**

*E. coli* DH5α carrying the plasmid pET22b-PstS-A197C was kindly donated by Martin Webb and distributed by Addgene (Addgene plasmid #78198) as an agar stab. Site-directed mutagenesis to introduce Cys mutations was performed using QuikChange II (Agilent), according to manufacturer's instructions.

The appropriate PstS encoding plasmid was transformed into BL21 (DE3) cells, with ampicillin added. A single colony was selected after overnight growth and used to inoculate 20 mL LB medium with the same antibiotics. This culture was grown at 37 °C overnight. 20 mL of starter culture was then added to 1 L LB medium containing the same antibiotics and grown at 37 °C until $OD_{600}$ = 0.6-0.8. IPTG was added, to a final concentration of 1 mM and the flask was shaken at 37 °C for 2-4 h. The cells were harvested by centrifugation (9,000 rpm, 20 min, 4 °C), suspended in 12.5 mL of lysis buffer each (10 mM Tris base, 1 mM DTT, pH 8.6, one tablet cOmplete™ Mini EDTA-free Protease Inhibitor Cocktail (Roche)) and frozen in liquid nitrogen, and stored at -80 °C.

Frozen cells were thawed in a water bath, when the solution was viscous DNase I was added, sonicated (10 cycles, 40% amplitude, 15 sec. on, 60 sec. off) and the cell debris was removed by centrifugation (30,000 rpm, 30 min, 4 °C). The solution was loaded onto a HiTrap Q HP column (5 mL) (GE Healthcare) and washed with 10 CV of binding buffer (10 mM Tris base, 1 mM DTT, pH 8.6) and eluted with a 20 CV gradient 0−100% elution buffer (10 mM Tris base, 200 mM NaCl, 1 mM DTT, pH 8.6). The fractions were analysed by SDS-PAGE and clean fractions were pooled together. The protein concentration was determined using an A280 spectrophotometer. The expression yield was determined to be 20 mg (30 mg/L expression volume). The protein solution was divided into aliquots, frozen in liquid nitrogen and stored at −80 °C until needed.

**PstS-D57C**

MEASLTGAGATFPAPVYAKWADTYQKETGNKVNYQGIGSSGGVKQIIANTVDFGAS
CAPLSDEKLAQEGLFQFPTVIGGVVLAVNIPGLKSGELVLDGKTLGDIYLGKIKKWD
DEAIAKLNPGLKLPSQNIAVVRRADGSGTSFVFTSYLAKVNEEWKNNVGTGSTVKW
PIGLGGKGNDGIAAFVQRLPGAIGYVEYAYAKQNNLAYTKLISADGKPVSPTEENFA
NAAKGADWSKTFAQDLTNQKGEDAWPITSTTFILIHKDQKKPEQGTEVLKFFDWAY
KTGAKQAN DLDYASLPDSVVEQVRAAWKTNIKDSSGKPLY

Calculated mass = 34541

Observed mass = 34541

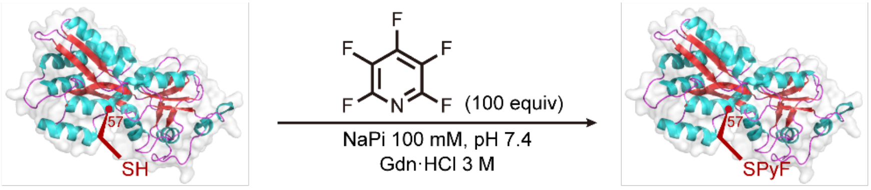

PstS-D57C PstS-Fpc57

The Cys-tetrafluoropyridylsulfide-containing PstS-Fpc57 was prepared according to general procedure 1.

ESI-MS spectrum for the modified PstS is shown below.

Calculated mass = 34690

Observed mass = 34690

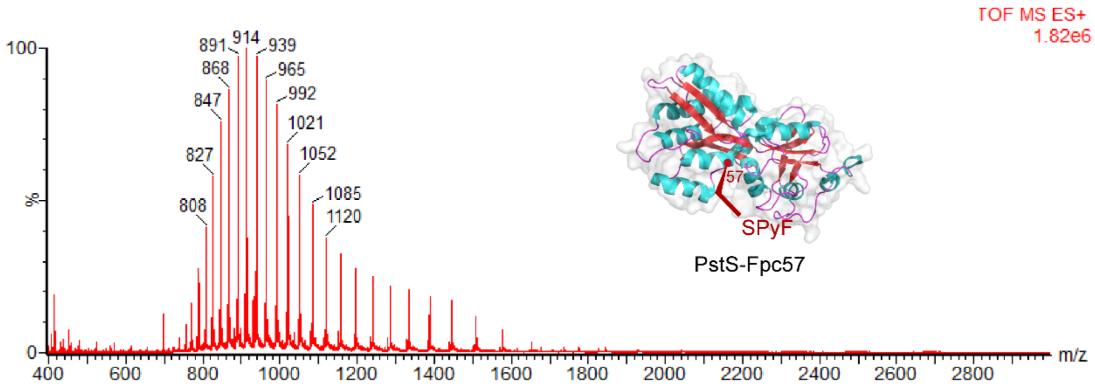

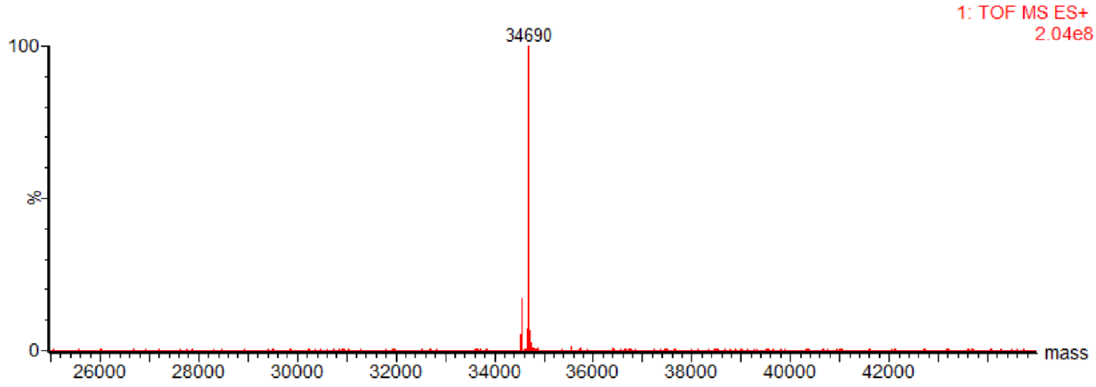

**PstS-D103C**

MEASLTGAGATFPAPVYAKWADTYQKETGNKVNYQGIGSSGGVKQIIANTVDFGAS
DAPLSDEKLAQEGLFQFPTVIGGVVLAVNIPGLKSGELVLDGKTLGCIYLGKIKKWD
DEAIAKLNPGLKLPSQNIAVVRRADGSGTSFVFTSYLAKVNEEWKNNVGTGSTVKW
PIGLGGKGNDGIAAFVQRLPGAIGYVEYAYAKQNNLAYTKLISADGKPVSPTEENFA
NAAKGADWSKTFAQDLTNQKGEDAWPITSTTFILIHKDQKKPEQGTEVLKFFDWAY
KTGAKQANDLDYASLPDSVVEQVRAAWKTNIKDSSGKPLY

Calculated mass = 34541
Observed mass = 34541

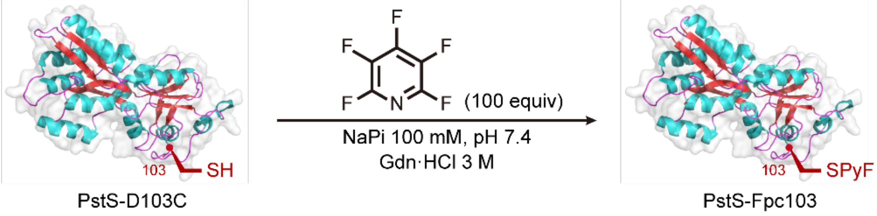

The Cys-tetrafluoropyridylsulfide-containing PstS-Fpc103 was prepared according to general procedure 1.

ESI-MS spectrum for the modified PstS is shown below.

Calculated mass = 34690

Observed mass = 34690

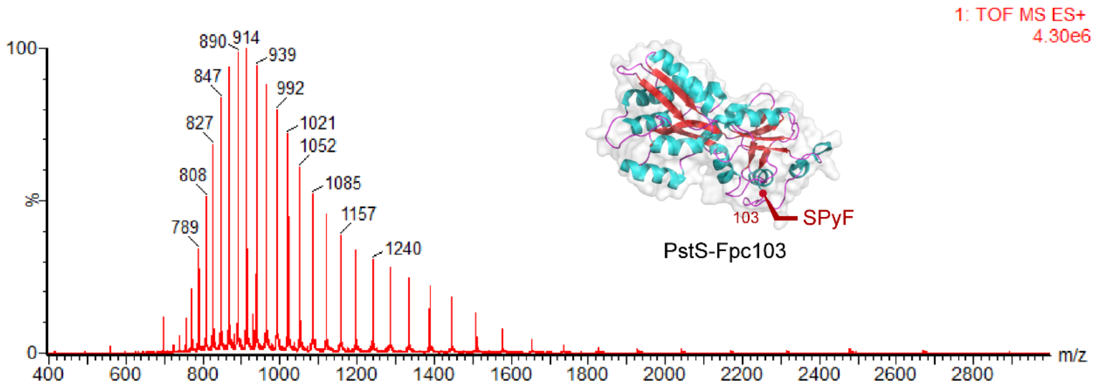

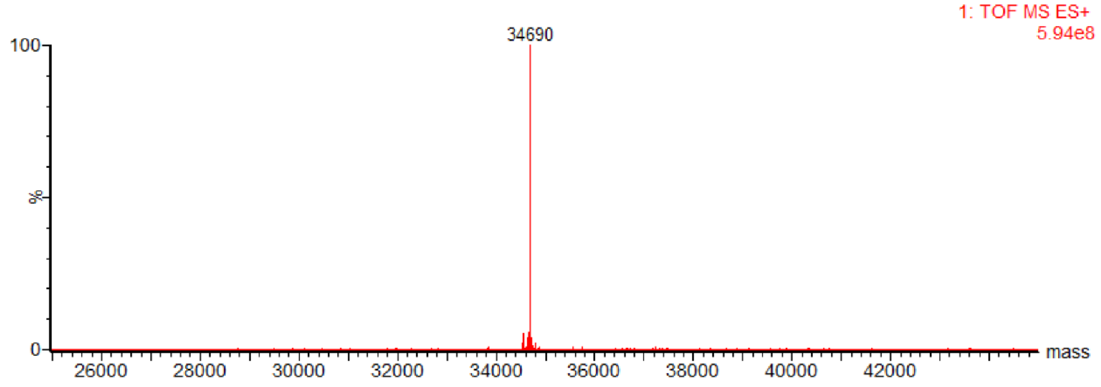

**PstS-D178C**

MEASLTGAGATFPAPVYAKWADTYQKETGNKVNYQGIGSSGGVKQIIANTVDFGAS

DAPLSDEKLAQEGLFQFPTVIGGVVLAVNIPGLKSGELVLDGKTLGDIYLGKIKKWD

DEAIAKLNPGLKLPSQNIAVVRRADGSGTSFVFTSYLAKVNEEWKNNVGTGSTVKW

PIGLGGKGNCGIAAFVQRLPGAIGYVEYAYAKQNNLAYTKLISADGKPVSPTEENFA

NAAKGADWSKTFAQDLTNQKGEDAWPITSTTFILIHKDQKKPEQGTEVLKFFDWAY

KTGAKQANDLDYASLPDSVVEQVRAAWKTNIKDSSGKPLY

Calculated mass = 34541

Observed mass = 34541

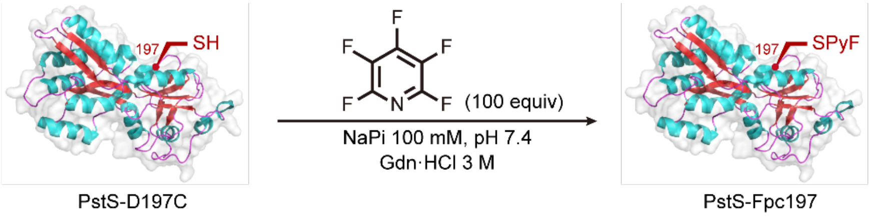

The Cys-tetrafluoropyridylsulfide-containing PstS_178PyfS was prepared according to general procedure 1.

ESI-MS spectrum for the modified PstS is shown below.

Calculated mass = 34690

Observed mass = 34690

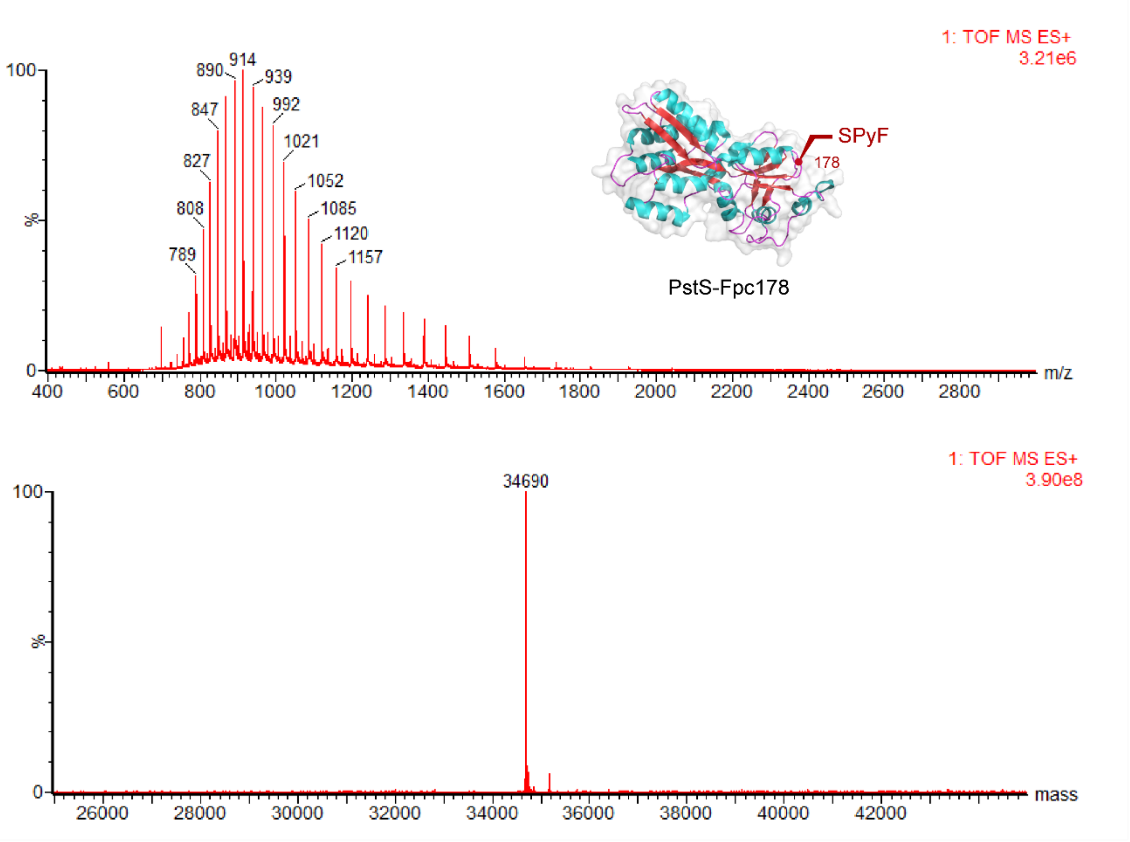

**PstS-A197C**

MEASLTGAGATFPAPVYAKWADTYQKETGNKVNYQGIGSSGGVKQIIANTVDFGAS
DAPLSDEKLAQEGLFQFPTVIGGVVLAVNIPGLKSGELVLDGKTLGDIYLGKIKKWD
DEAIAKLNPGLKLPSQNIAVVRRADGSGTSFVFTSYLAKVNEEWKNNVGTGSTVKW
PIGLGGKGNDGIAAFVQRLPGAIGYVEYCYAKQNNLAYTKLISADGKPVSPTEENFA
NAAKGADWSKTFAQDLTNQKGEDAWPITSTTFILIHKDQKKPEQGTEVLKFFDWAY
KTGAKQANDLDYASLPDSVVEQVRAAWKTNIKDSSGKPLY

Calculated mass = 34585

Observed mass = 34585

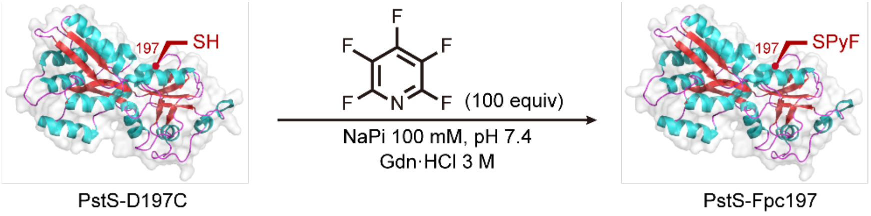

The Cys-tetrafluoropyridylsulfide-containing PstS-Fpc197 was prepared according to general procedure 1.

ESI-MS spectrum for the modified PstS is shown below.

Calculated mass = 34734

Observed mass = 34734

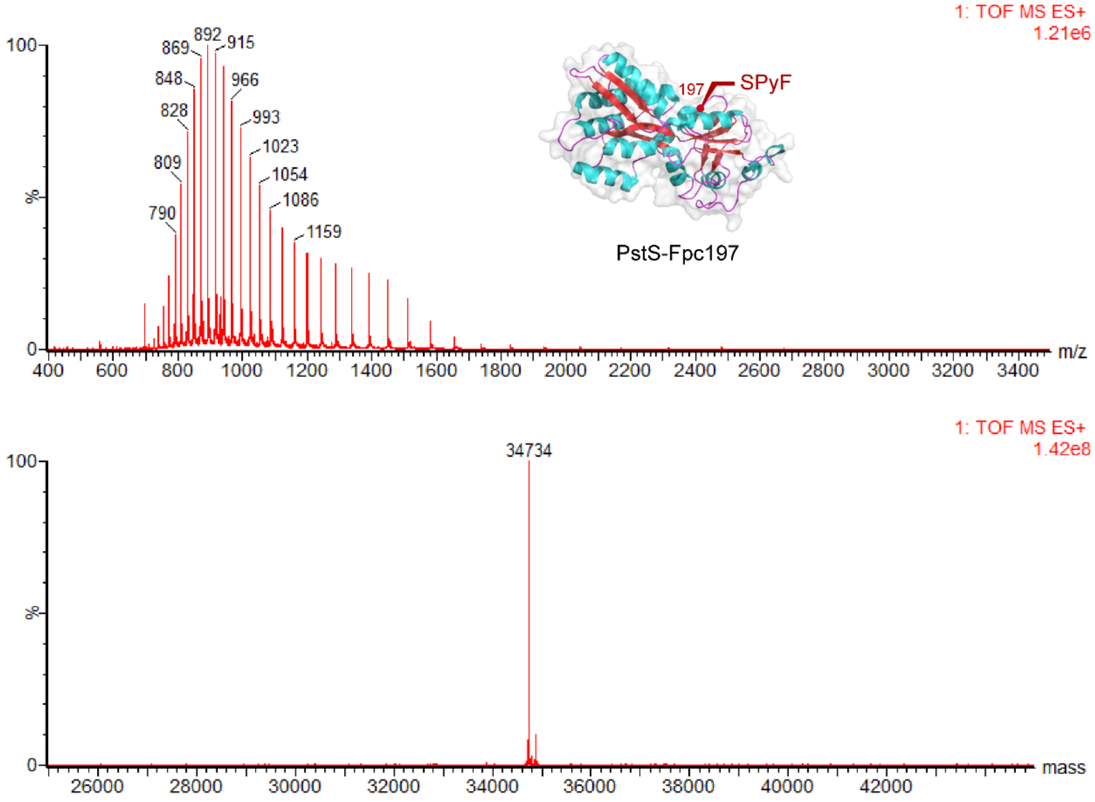

**pre-SUMO1-Fpc51: protein expression, purification and Fpc generation**

pre-SUMO1-Cys51 protein was expressed and purified following a previously published procedure.[3]

**pre-SUMO1-Cys51**

ADQEAKPSTEDLGDKKEGEYIKLKVIGQDSSEIHFKVKMTTHLKKLKESYCQRQGVPMNSLRFLFEGQRIADNHTPKELGMEEEDVIEVYQEQTGGHSTVLEHHHHHH

Calculated mass = 12475

Observed mass = 12474

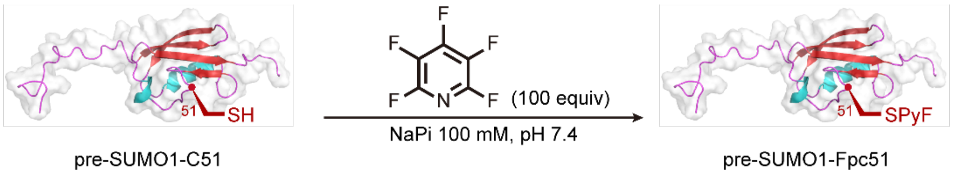

The Cys-tetrafluoropyridylsulfide-containing pre-SUMO1-Fpc51 was prepared according to general procedure 1.

ESI-MS spectrum for the modified pre-SUMO1 is shown below.

Calculated mass = 12623

Observed mass = 12623

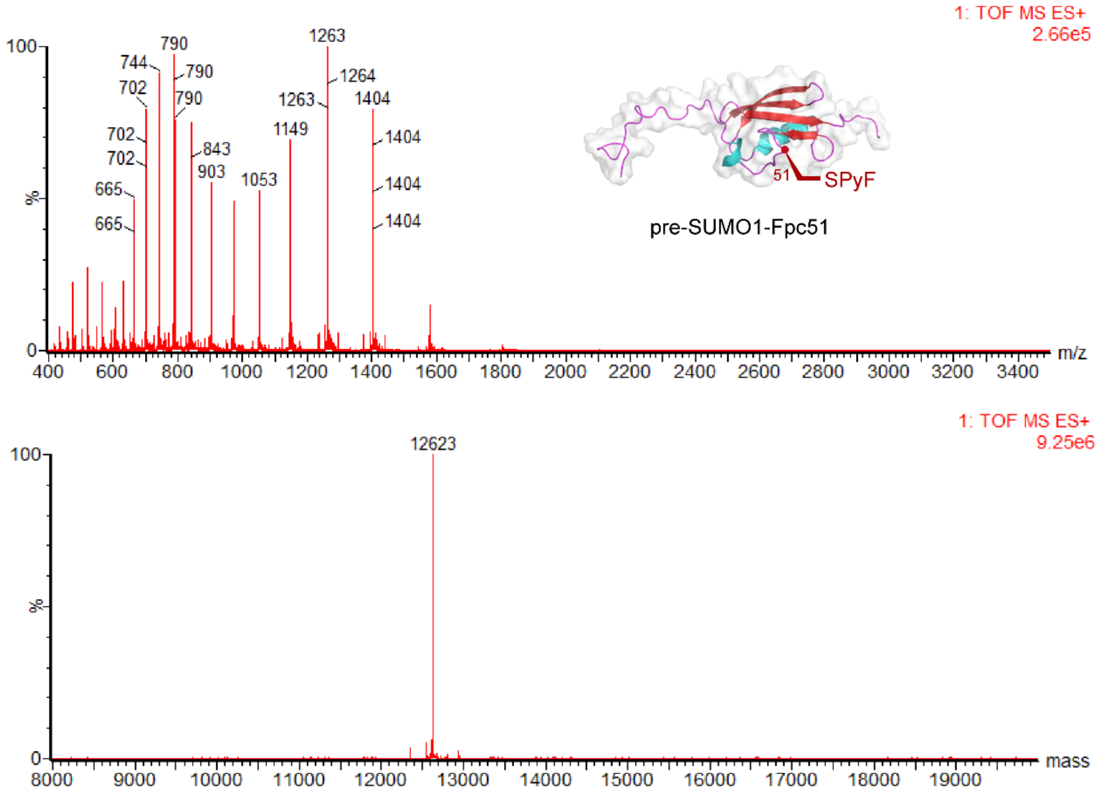

**Npβ-Fpc61: protein expression, purification and Fpc generation**

Npβ-C61 was expressed and purified following a previously published procedure.[3]

**Npβ-Cys61**

MFSSHHHHHHSSGLVPRGSHIDVGKLRQLYAAGERDFSIVDLRGAVLENINLSGAIL HGACLDEANLQQANLSRADLSGATLNGADLRGANLSKADLSDAILDNAILEGAILDE AVLNQANLKAANLEQAILSHANIREADLSEANLEAADLSGADLAIADLHQANLHQA ALERANLTGANLEDANLEGT ILEGGNNNLAT

Calculated mass = 21031

Observed mass = 21031

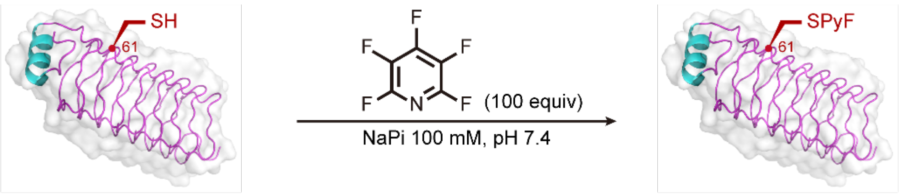

Npβ-C61 Npβ-Fpc61

The Cys-tetrafluoropyridylsulfide-containing Npβ-Fpc61 was prepared according to general procedure 1.

ESI-MS spectrum for the modified Npβ is shown below.

Calculated mass = 21180

Observed mass = 21180

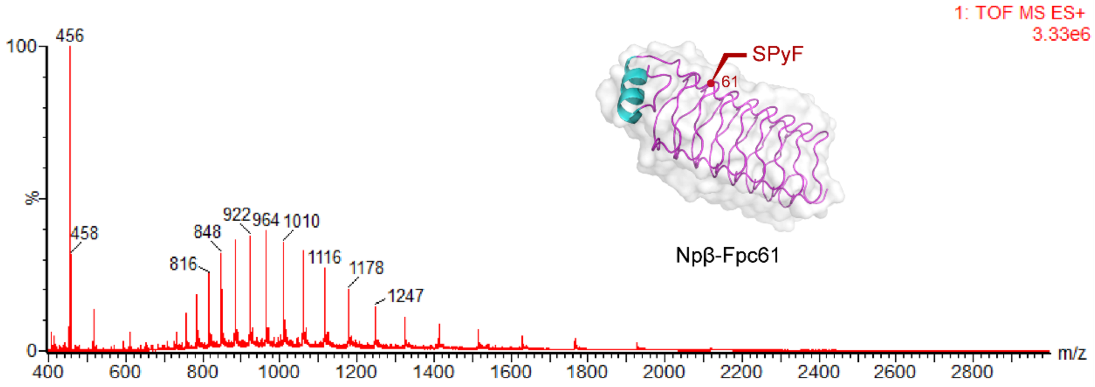

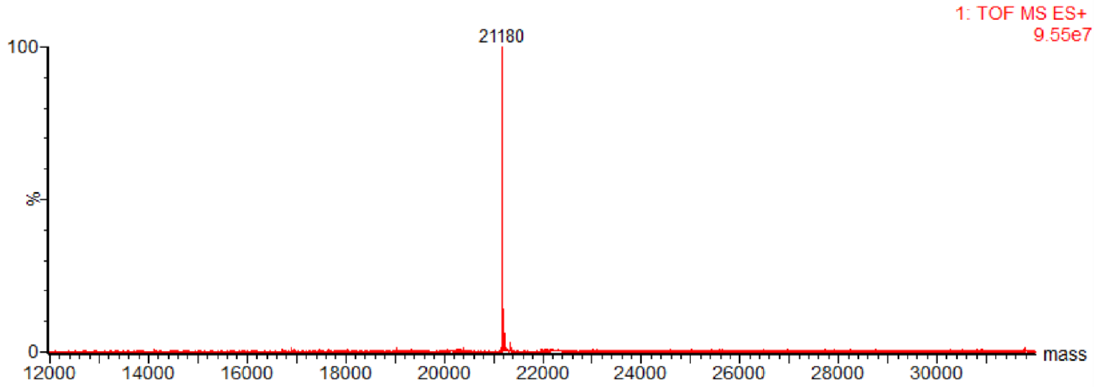

**cAbVCAM1-Fpc118: protein expression, purification and Fpc generation**

A fresh stock of WK6 competent *E. coli* cells (received as a gift from Prof. Ray Owens) was amplified according to the manufacturer's instructions

Single colonies were transferred to 15 mL of LB media supplemented with ampicillin (100 µg/mL) and incubated at 37 °C for 16 hours. The resulting suspension was immediately used to inoculate 1 L Terrific Broth (TB) media supplemented with ampicillin (100 µg/mL, 0.1 % glucose, and 2 mM $MgCl_2$. The cultures were incubated at 37 °C (180 rpm), for approximately 4 hours, until an $OD_{600}$ between 0.9 and 1.1 was reached. Protein overexpression was induced by the addition of IPTG (final conc. 1 mM) and the cultures were incubated for 16 more hours at 27 °C (180 rpm). The cells were harvested by centrifugation (12,000 x g, 10 minutes, 4 °C) to afford cell pellets (25 g wet weight per liter of culture). The cell pellets were kept at -80 °C until further manipulation.

A single cell pellet was thawed on ice and mixed with 40 mL of TES lysis buffer (0.2 M Tris pH 7.8, 0.5 mM EDTA, 0.5 M sucrose) containing one pre-dissolved cOmplete protease inhibitor mix tablet (EDTA free, Roche). The cell pellet was vortexed until bacterial clumps were not visible and then shaken via end-over-end mixing for 30 minutes at 4 °C. 2 mg of DNase I were added, and the mixture was further shaken via end-over-end mixing for 2 hours at 4 °C. The lysate was centrifuged at 4 °C and 22,000× g for 15 min.

Supernatant was filtered through 0.2 µm syringe filter and loaded to a pre-equilibrated HisTrap HP 5 mL column (GE Healthcare, 2.5 mL/min), using a 50 mL superloop (GE Healthcare). The protein was eluted running a stepwise gradient of 30 CV to 100% buffer B (10 CV Buffer A, 2 CV 5% Buffer B, 2 CV 7% Buffer B, 2 CV 10% Buffer B, 2 CV 20% Buffer B, 2 CV 35% Buffer B, 8 CV 100% Buffer B). The fractions were analysed by SDS-PAGE and clean fractions containing protein were combined. The protein fractions were buffer exchanged to 50 mM NaPi, pH 8 using an Amicon® Ultra-15 Centrifugal Filter Unit and samples were flash-frozen in liquid nitrogen and stored at -80 °C. Protein expression yield was measured after the final buffer exchange and was measured at 11.2 mg/L.

Buffer A - 20 mM Tris-HCl, 15 mM imidazole, 500 mM NaCl, 1 mM DTT, 0.05% (v/v) β-mercaptoethanol, pH 7.8

Buffer B - 20 mM Tris-HCl, 500 mM imidazole, 500 mM NaCl, 1 mM DTT, 0.05% (v/v) β-mercaptoethanol, pH 7.8

**cAbVCAM-Cys118**

NVQLQESGGGSVQTGGSLRLSCAASGYTNSIMYMAWFRQAPGKKREGVAAIRFPDD
SAYYAGSVKGRFTISHDNAKNTVYLQMNNLNPEDTAMYYCAARSSPYSFAWNDPN
YNYWGCGTQVTVSSHHHHHH

Calculated mass = 14619

Observed mass = 14619

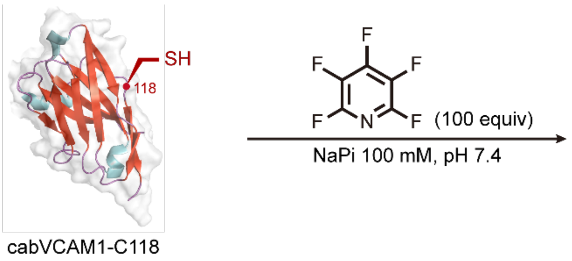

The Cys-tetrafluoropyridylsulfide-containing cabVCAM-Fpc118 was prepared according to general procedure 1.

ESI-MS spectrum for the modified cAbVCAM is shown below.

Calculated mass = 14768

Observed mass = 14767

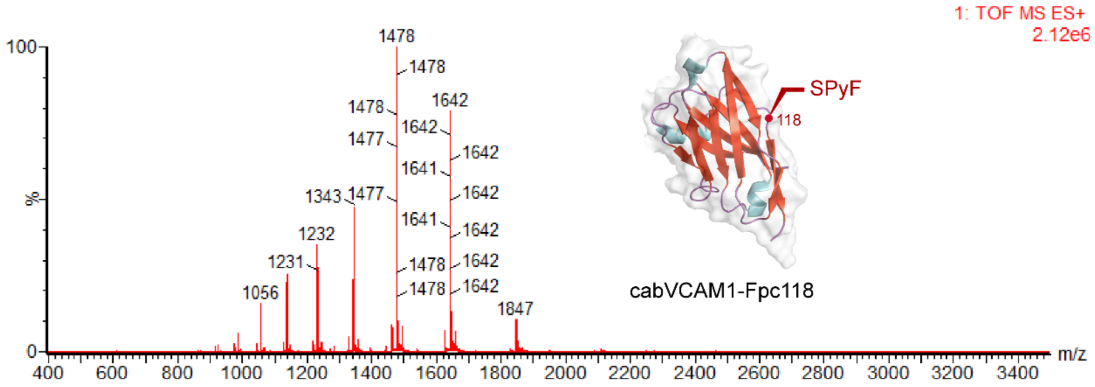

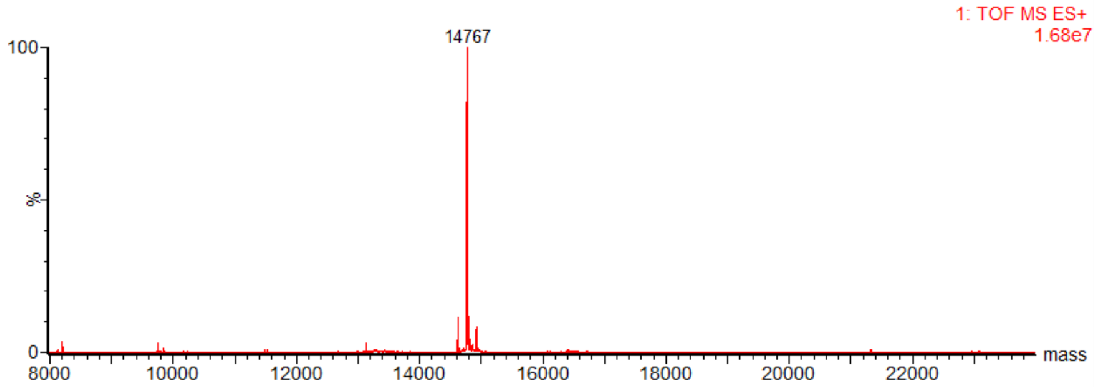

**AcrA-Fpc123: protein expression, purification and Fpc generation**

AcrA-C123 was expressed and purified following a previously published procedure.[2]

**AcrA-Cys123**

SKEEAPKIQMPPQPVTTMSAKSEDLPLSFTYPAKLVSDYDVIIKPQVSGVIVNKLFKA
GDKVKKGQTLFIIEQDKFKASVDSAYGQALMAKATFENASKDFCRSKALFSKSAISQ
KEYDSSLATFNNSKASLASARAQLANARIDLDHTEIKAPFDGTIGDALVNIGDYVSAS
TTELVRVTNLNPIYADFFISDTDKLNLVRNTQSGKWDLDSIHANLNLNGETVQGKLY
FIDSVIDANSGTVKAKAVFDNNNSTLLPGAFATITSEGFIQKNGFKVPQIGVKQDQND
VYVLLVKNGKVEKSSVHISYQNNEYAIIDKGLQNGDKIILDNFKKIQVGSEVKEIGAQ
LEHHHHHH

Calculated mass = 38817

Observed mass = 38817

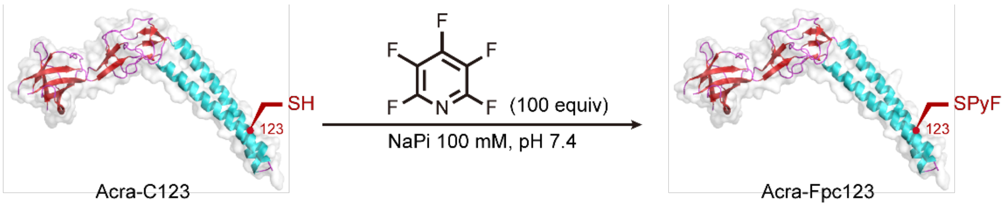

The Cys-tetrafluoropyridylsulfide-containing Acra-Fpc123 was prepared according to general procedure 1.

ESI-MS spectrum for the modified Acra is shown below.

Calculated mass = 38966

Observed mass = 38966

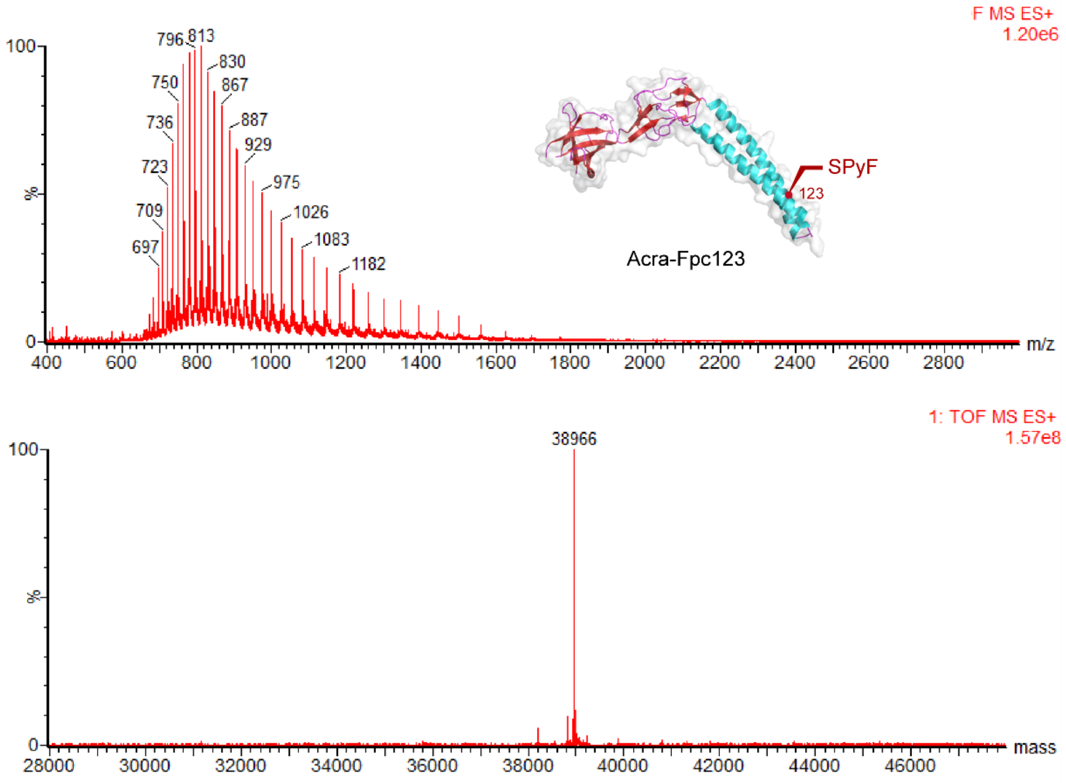

### On-protein radical trapping from PstS-Fpc178

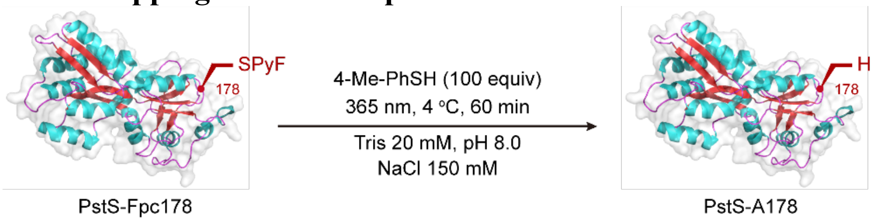

The PstS-A178 was prepared according to general procedure 2.

ESI-MS spectrum for the modified PstS is shown below.

Calculated mass = 34509

Observed mass = 34509

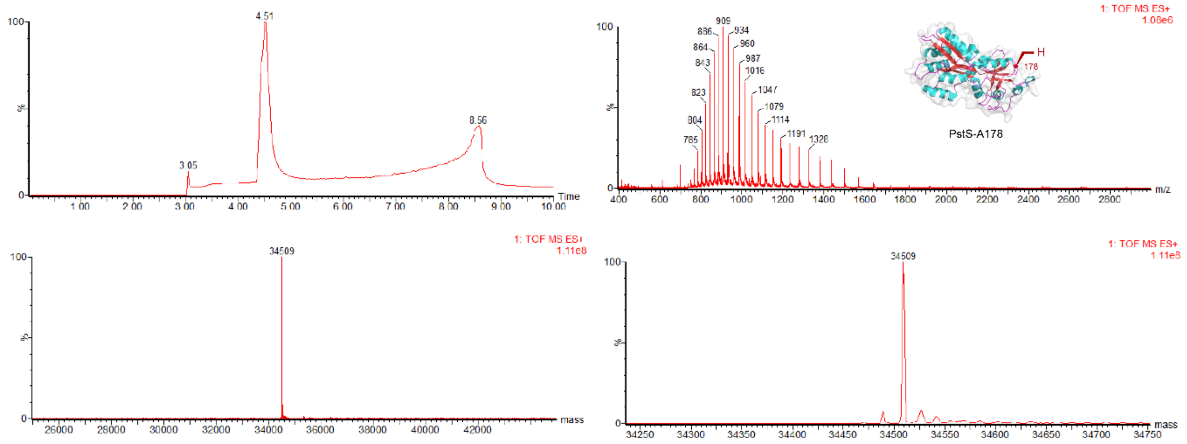

Conversion

| PstS-Fpc178 | PstS-Ala178 |
|---|---|
| 0% | >98% conversion |

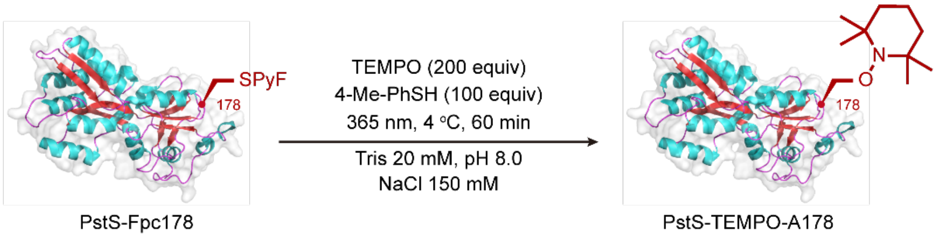

The PstS-TEMPO-A178 was prepared according to general procedure 3.

ESI-MS spectrum for the modified PstS is shown below.

Calculated mass = 34664

Observed mass = 34663

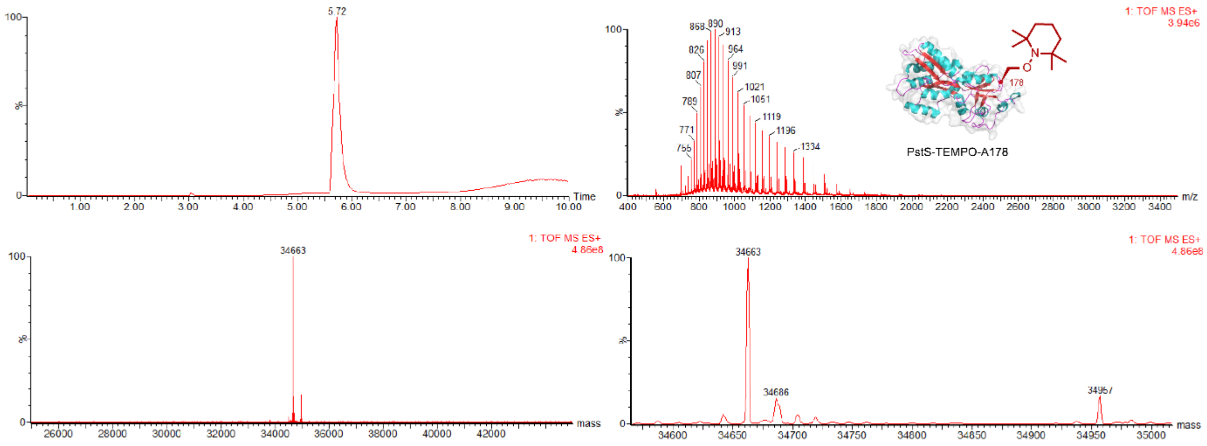

Conversion

| PstS-Fpc178 | PstS-Ala178 | PstS-TEMPO-A178 |
|---|---|---|
| 10% | 0% | 90% |

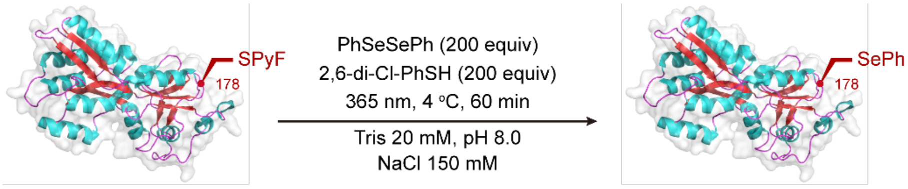

PstS-Fpc178 PstS-SecPh178

The PstS-SecPh178 was prepared according to general procedure 4.

ESI-MS spectrum for the modified PstS is shown below.

Calculated mass = 34664

Observed mass = 34664

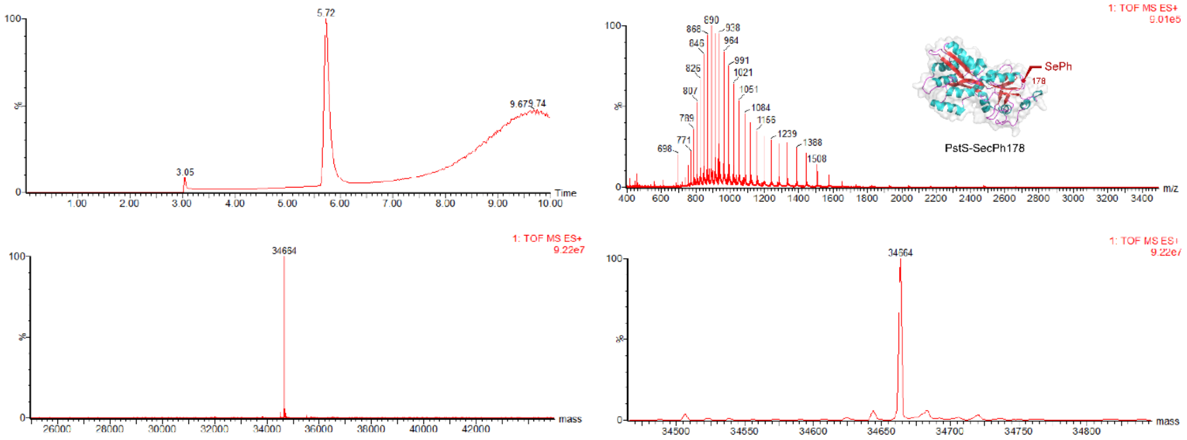

Conversion

| PstS-Fpc178 | PstS-Ala178 | PstS-SecPh178 |
|---|---|---|
| 0% | 0% | >98% conversion |

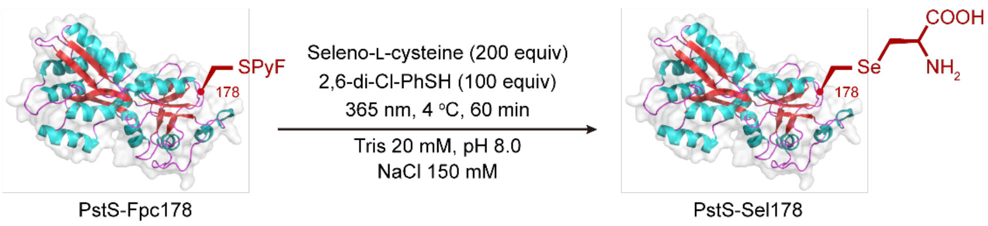

The PstS-Sel178 was prepared according to general procedure 4.

ESI-MS spectrum for the modified PstS is shown below.

Calculated mass = 34675

Observed mass = 34675

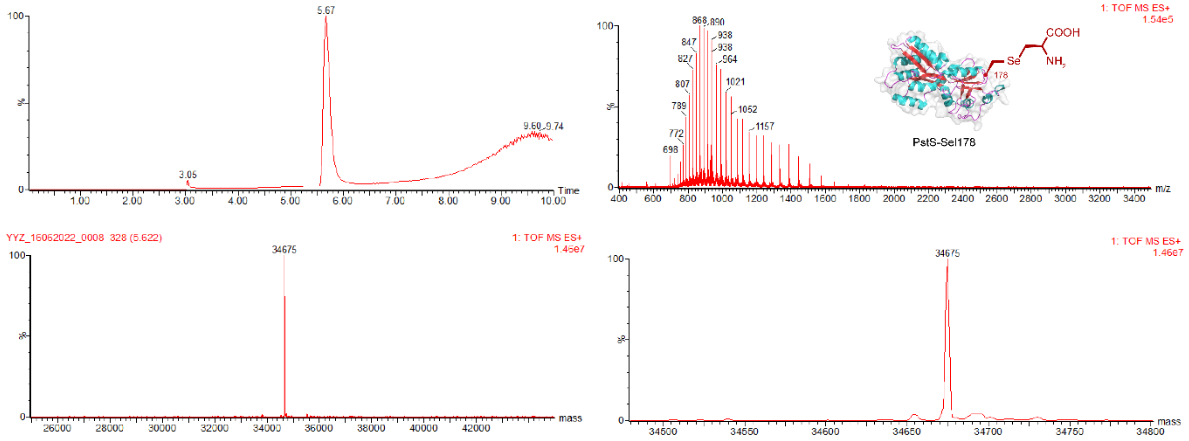

Conversion

| PstS-Fpc178 | PstS-Ala178 | PstS-SecPh178 |
|---|---|---|
| 0% | 0% | >98% conversion |

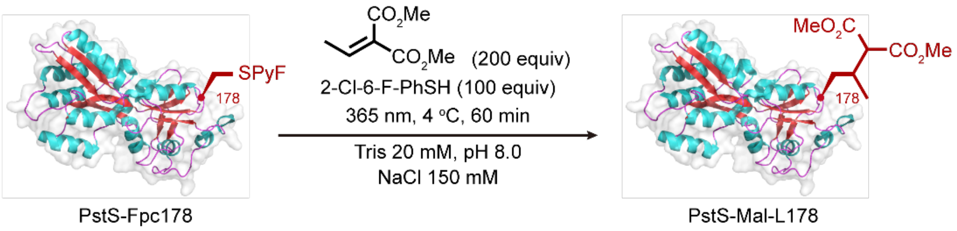

The PstS-Mal-L178 was prepared according to general procedure 5.

ESI-MS spectrum for the modified PstS is shown below.

Calculated mass = 34667

Observed mass = 34668

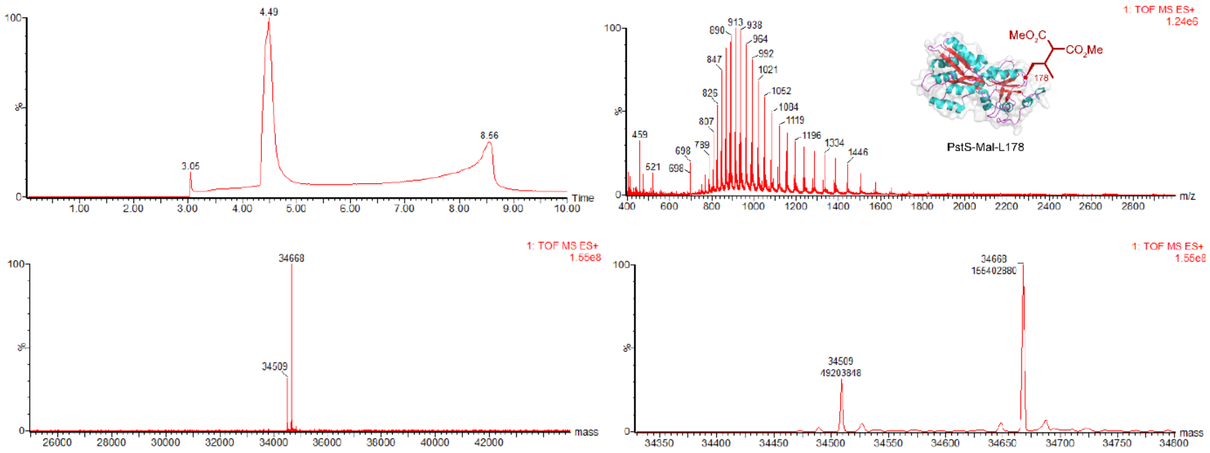

Conversion

| PstS-Fpc178 | PstS-Ala178 | PstS-Mal-L178 |
|---|---|---|
| 0% | 25% | 75% |

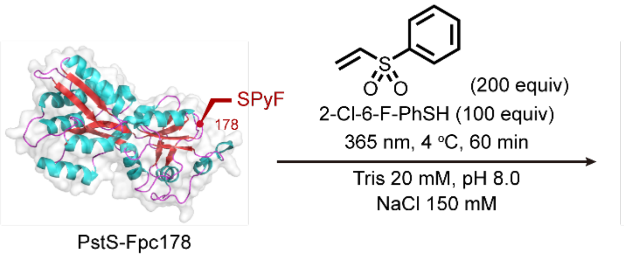

The PstS-A(Sulfone)178 was prepared according to general procedure 5.

ESI-MS spectrum for the modified PstS is shown below.

Calculated mass = 34676 (1 adduct), 34844 (2 adducts), 35012 (3 adducts)

Observed mass = 34676 (1 adduct), 34844 (2 adducts), 35013 (3 adducts)

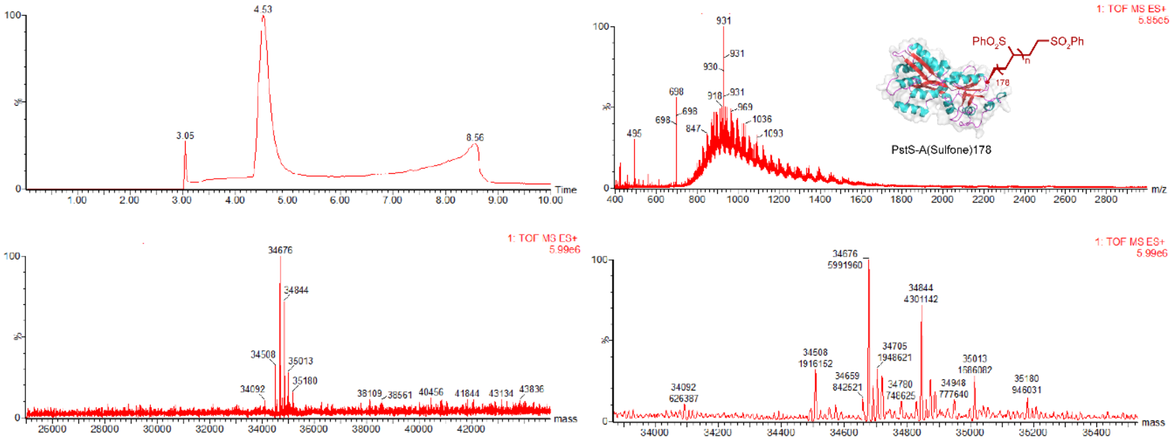

Conversion

| PstS-Fpc178 | PstS-Ala178 | PstS-A(Sulfone)178 (1-4 adducts) |
|---|---|---|
| 0% | 13% | 87% |

See also **Supplementary Figure 15**, which suggests that adducts may be non-specific.

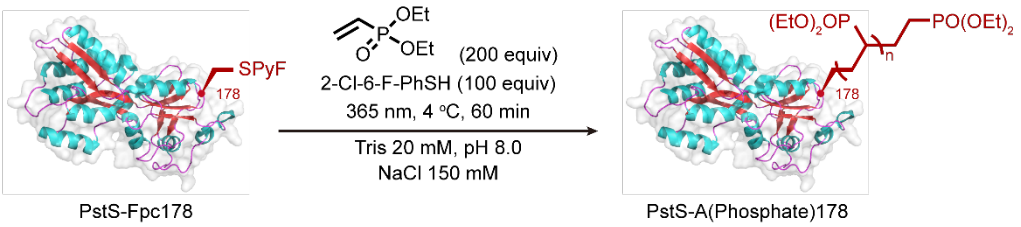

The PstS-A(Phosphate)178 was prepared according to general procedure 5.

ESI-MS spectrum for the modified PstS is shown below.

Calculated mass = 34673 (n =0), 34837 (n = 1), 35001 (n = 2), 35165 (n = 3)

Observed mass = 34672 (n =0), 34837 (n = 1), 35000 (n = 2), 35164 (n = 3)

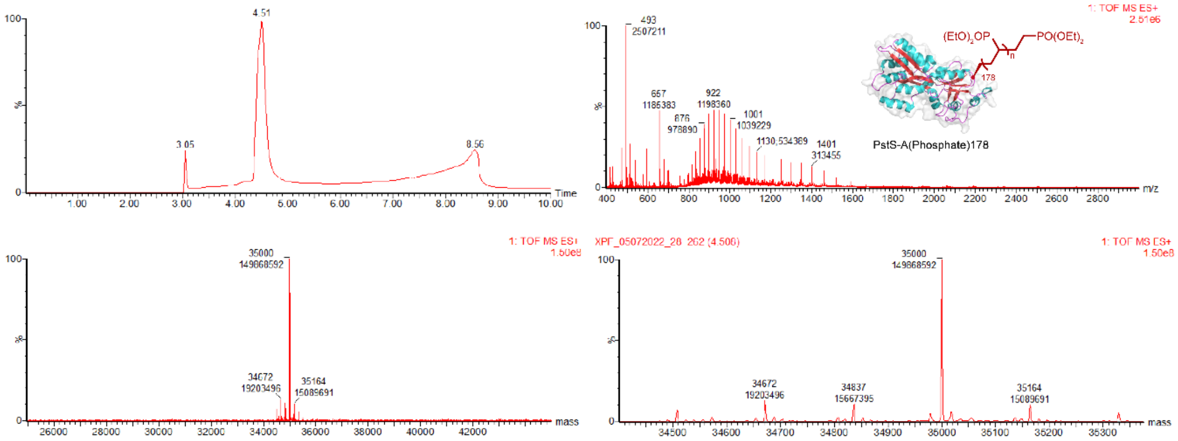

Conversion

| PstS-Fpc178 | PstS-Ala178 | PstS-A(Phosphate)178 (n = 0-3) |
|---|---|---|
| 0% | 0% | >98% conversion |

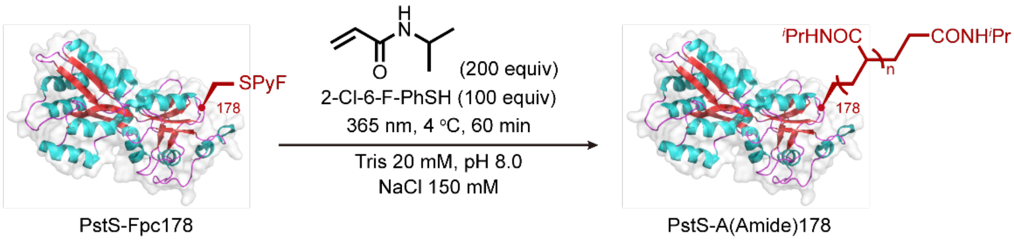

The PstS-A(Amide)178 was prepared according to general procedure 5.

ESI-MS spectrum for the modified PstS is shown below.

Calculated mass = 34847 (n = 2), 34960 (n = 3), 35073 (n = 4), 35186 (n = 5), 35299 (n = 6), 35412 (n = 7), 35525 (n = 8), 35638 (n = 9), 35751 (n = 10), 35864 (n = 11), 35977 (n = 12).

Observed mass = 34847 (n = 2), 34961 (n = 3), 35073 (n = 4), 35187 (n = 5), 35300 (n = 6), 35414 (n = 7), 35527 (n = 8), 35640 (n = 9), 35753 (n = 10), 35866 (n = 11), 35978 (n = 12).

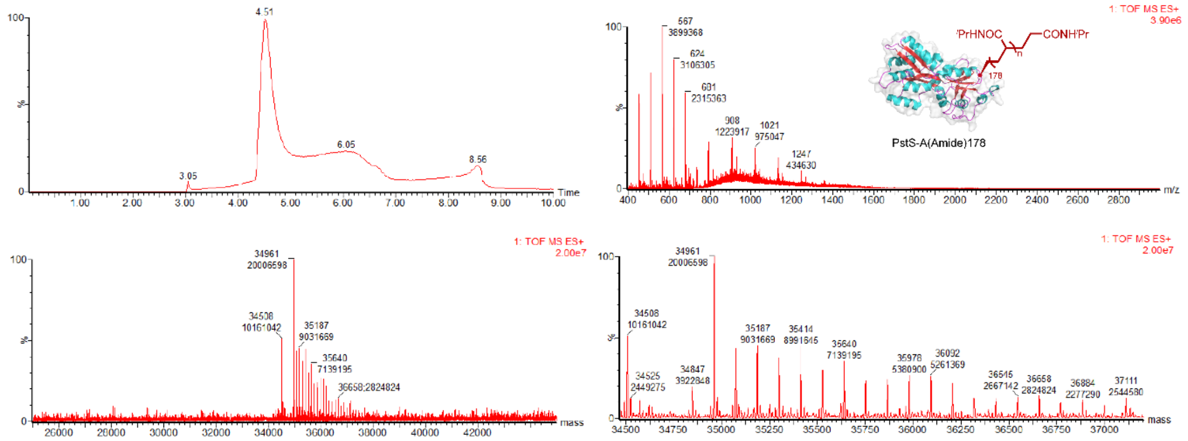

Conversion

| PstS-Fpc178 | PstS-Ala178 | PstS-A(Amide)178 (n = 2-19) |
|---|---|---|
| 0% | 9% | 91% |

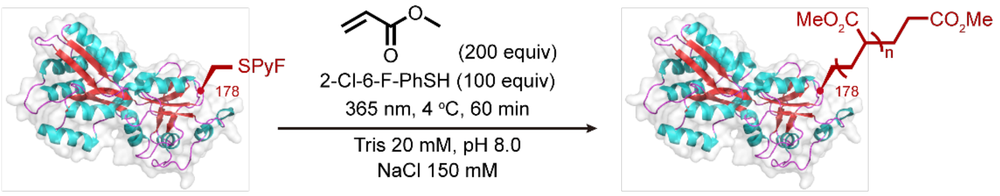

PstS-Fpc178 PstS-A(Ester)178

The PstS-A(Ester)178 was prepared according to general procedure 5.

ESI-MS spectrum for the modified PstS is shown below.

Calculated mass = 34766 (n = 2), 34852 (n = 3), 34938 (n = 4), 35024 (n = 5), 35110 (n = 6), 35196 (n = 7).

Observed mass = 34766 (n = 2), 34852 (n = 3), 35938 (n = 4), 35025 (n = 5), 35110 (n = 6), 35196 (n = 7).

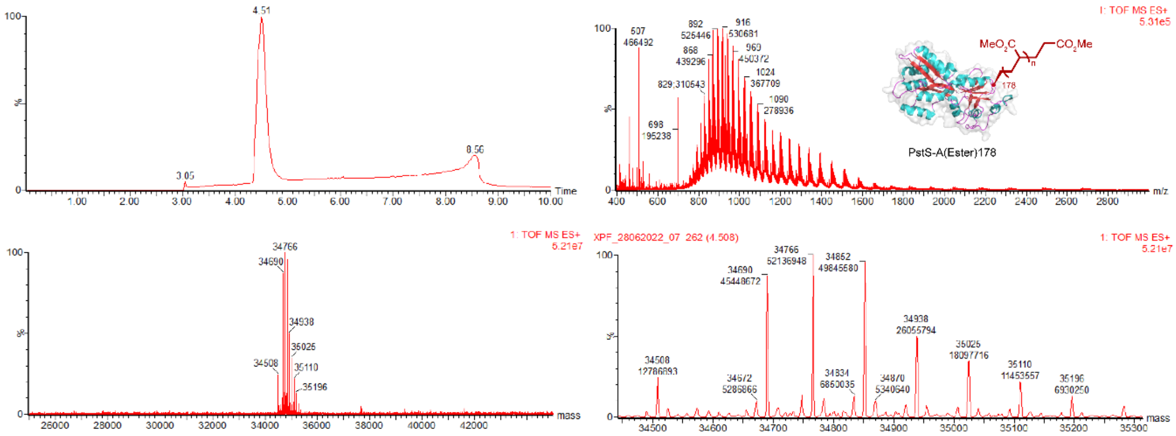

Conversion

| PstS-Fpc178 | PstS-Ala178 | PstS-A(Ester)178 (n = 2-7) |
|---|---|---|
| 20% | 6% | 74% |

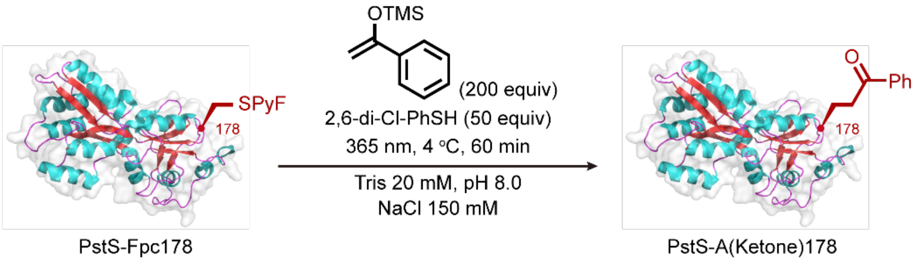

The PstS-A(Ketone)178 was prepared according to general procedure 5.

ESI-MS spectrum for the modified PstS is show below.

Calculated mass = 34627

Observed mass = 34628

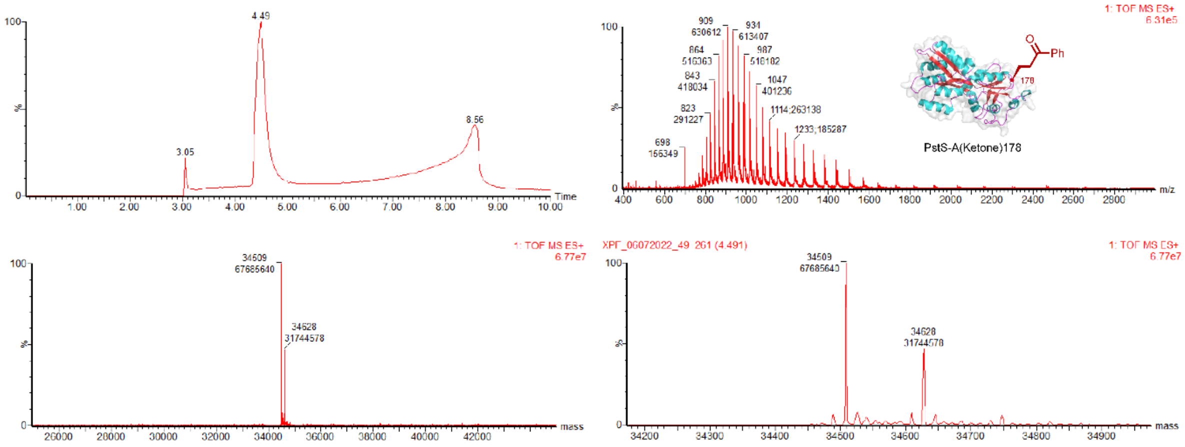

Conversion

| PstS-Fpc178 | PstS-Ala178 | PstS-A(Ketone)178 |
|---|---|---|
| 0% | 67% | 33% |

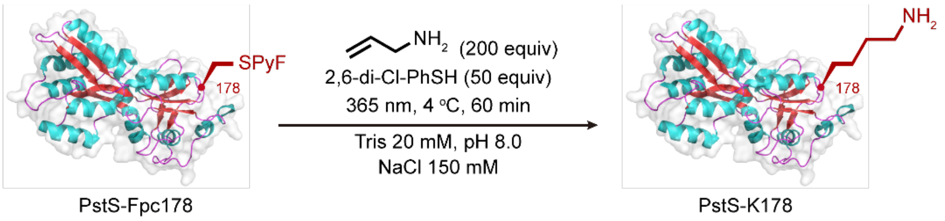

The PstS-Lys178 was prepared according to general procedure 5.

ESI-MS spectrum for the modified PstS is shown below.

Calculated mass = 34566

Observed mass = 34565

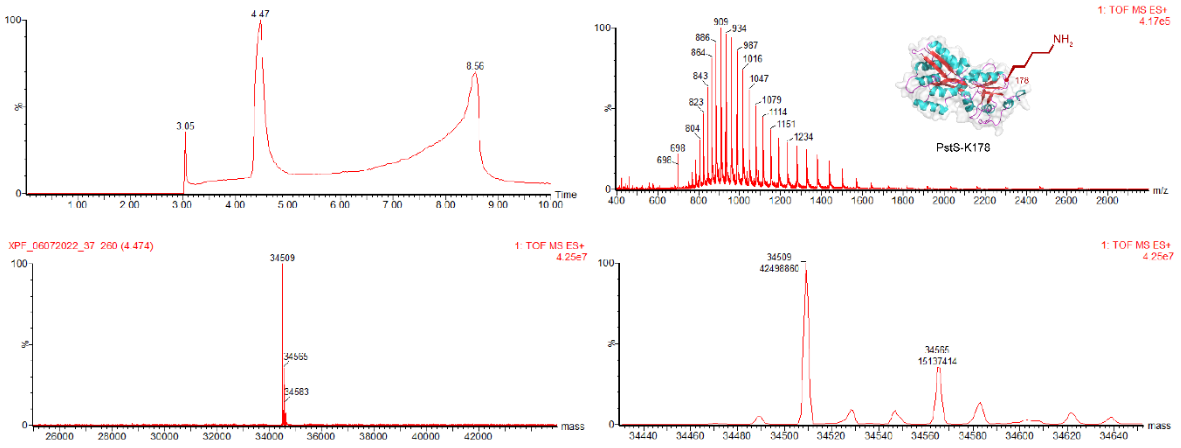

Conversion

| PstS-Fpc178 | PstS-Ala178 | PstS-Lys178 |
|---|---|---|
| 0% | 74% | 26% |

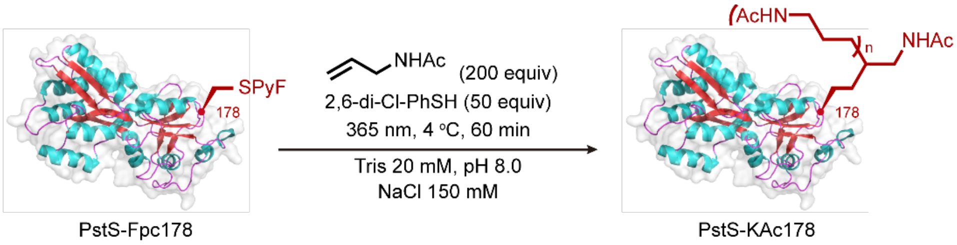

The PstS-KAc178 was prepared according to general procedure 5.

ESI-MS spectrum for the modified PstS is shown below.

Calculated mass = 34607 (n = 0), 34706 (n = 1)

Observed mass = 34607 (n = 0), 34706 (n = 1)

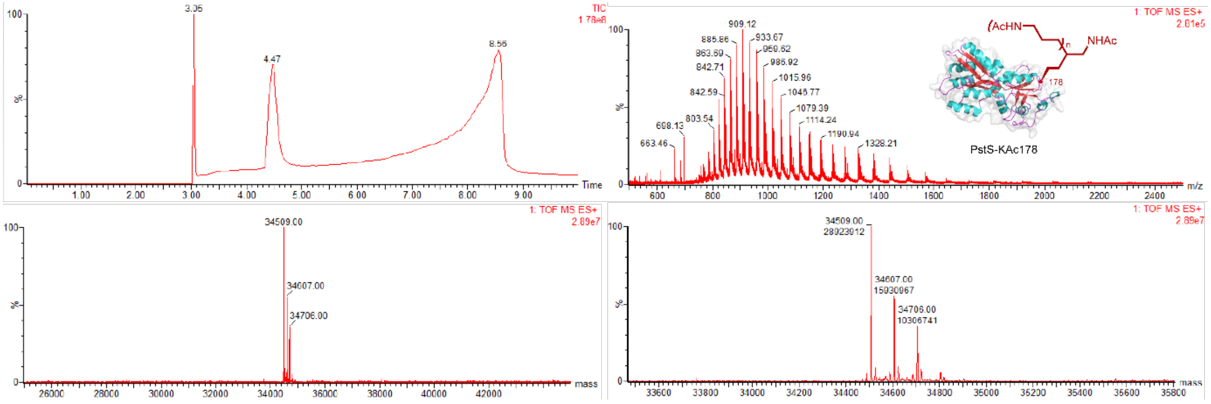

Conversion

| PstS-Fpc178 | PstS-Ala178 | PstS-KAc178 (n = 0-1) |
|---|---|---|
| 0% | 52% | 48% |

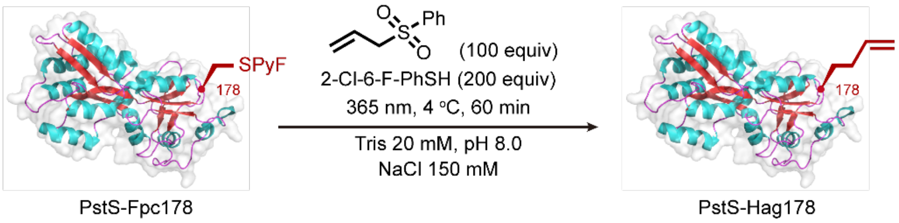

The PstS-Hag178 was prepared according to general procedure 5.

ESI-MS spectrum for the modified PstS is shown below.

Calculated mass = 34548

Observed mass = 34549

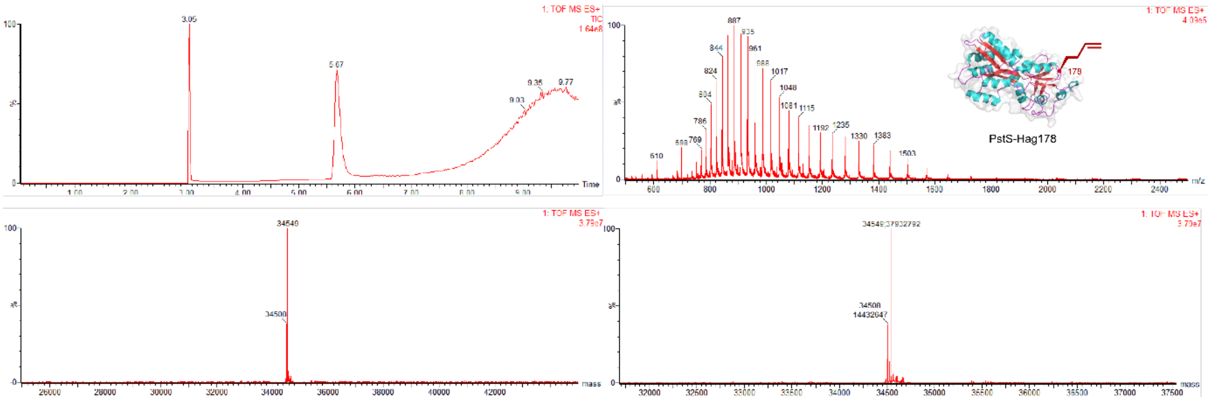

Conversion

| PstS-Fpc178 | PstS-Ala178 | PstS-Hag178 |
|---|---|---|
| 0% | 28% | 72% |

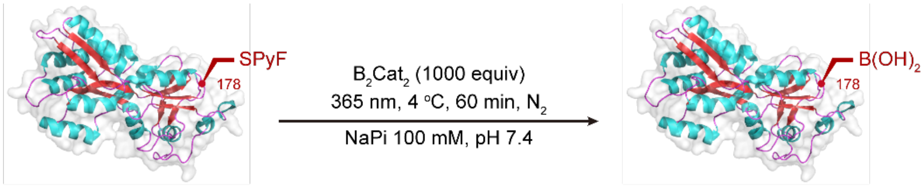

PstS-Fpc178 PstS-Bal178

The PstS-Bal178 was prepared according to general procedure 6.

ESI-MS spectrum for the modified PstS is shown below.

Calculated mass = 34553 (Bal), 34535 (Bal-$H_2O$)

Observed mass = 34552 (Bal), 34533 (Bal-$H_2O$)

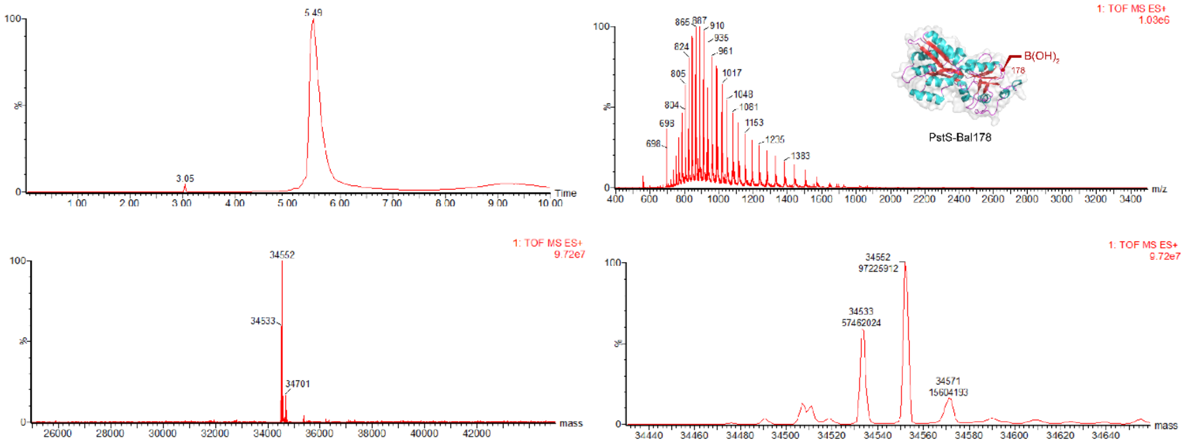

Conversion

| PstS-Fpc178 | PstS-Ala178 | PstS-Bal178 | |
|---|---|---|---|
| | | Bal | Bal-$H_2O$ |
| 0% | 9% | 57% | 34 |

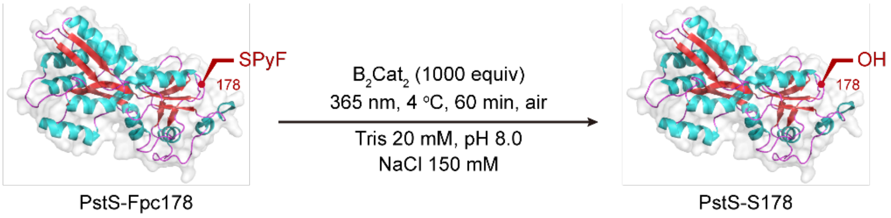

The PstS-Ser178 was prepared according to general procedure 7.

ESI-MS spectrum for the modified PstS is shown below.

Calculated mass = 34525

Observed mass = 34524

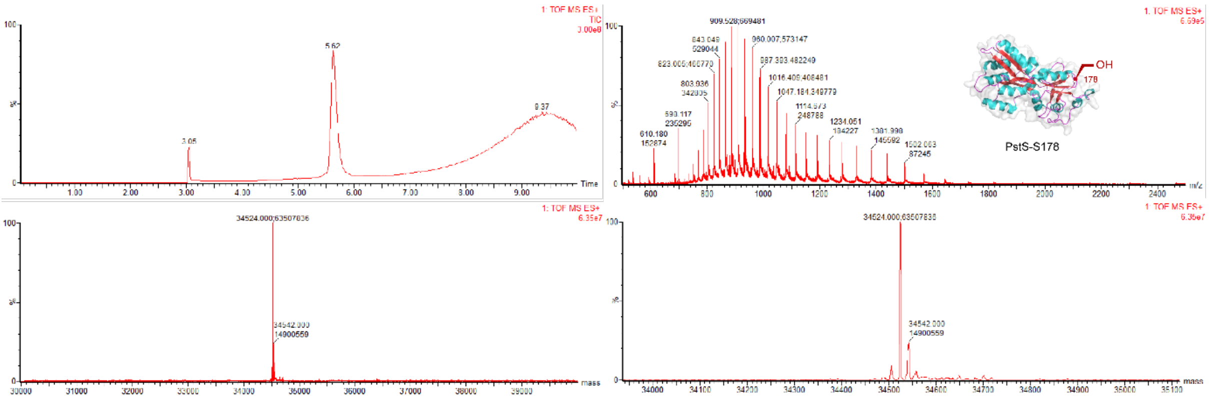

Conversion

| PstS-Fpc178 | PstS-Ala178 | PstS-Ser178 |
|---|---|---|
| 0% | 0% | 81% |

**Widescale introduction of L-boronoalanine (L-Bal) into Proteins.**

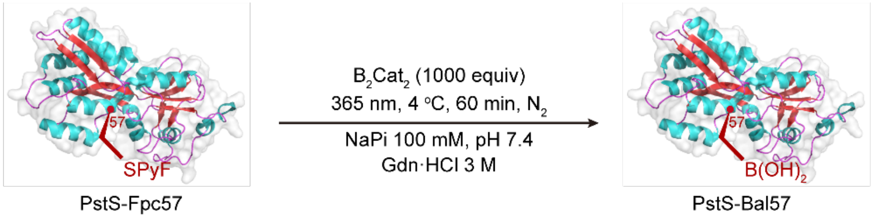

The PstS-Bal57 was prepared according to general procedure 8.

ESI-MS spectrum for the modified PstS is shown below.

Calculated mass = 34553 (Bal), 34535 (Bal-$H_2O$), 34517 (Bal-$2H_2O$)

Observed mass = 34551 (Bal), 34533 (Bal-$H_2O$), 34516 (Bal-$2H_2O$)

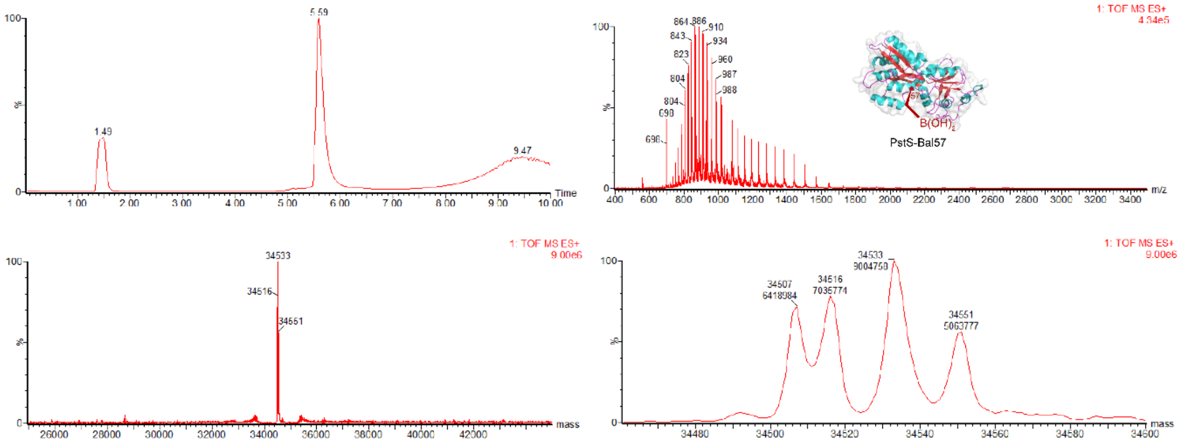

Conversion

| PstS-Fpc57 | PstS-A57 | PstS-Bal57 | | |
|---|---|---|---|---|
| | | Bal | Bal-$H_2O$ | Bal-$2H_2O$ |
| 0% | 23% | 25% | 32% | 20% |

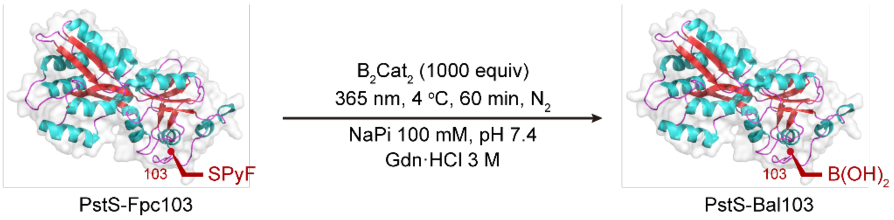

The PstS-Bal103 was prepared according to general procedure 8.

ESI-MS spectrum for the modified PstS is shown below.

Calculated mass = 34553 (Bal), 34535 (Bal-$H_2O$)

Observed mass = 34551 (Bal), 34536 (Bal-$H_2O$).

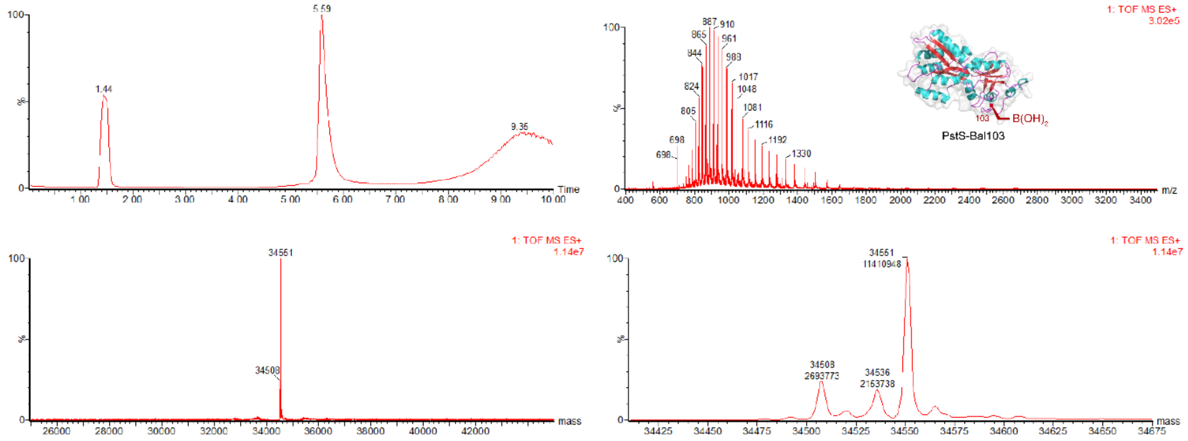

Conversion

| PstS-Fpc103 | PstS-A103 | PstS-Bal103 | | |
|---|---|---|---|---|
| | | Bal | Bal-$H_2O$ | Bal-$2H_2O$ |
| 0% | 17% | 70% | 13% | 0% |

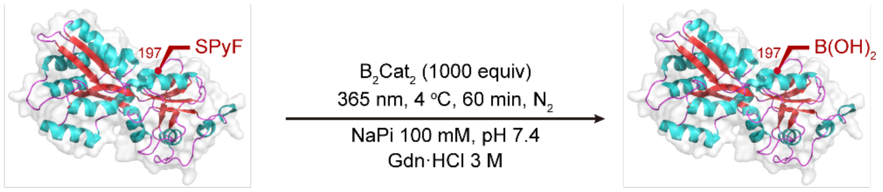

PstS-Fpc197 PstS-Bal197

The PstS-Bal197 was prepared according to general procedure 8.

ESI-MS spectrum for the modified PstS is shown below.

Calculated mass = 34597 (Bal), 34579 (Bal-$H_2O$), 34561 (Bal-$2H_2O$)

Observed mass = 34596 (Bal), 34578 (Bal-$H_2O$), 34562 (Bal-$2H_2O$)

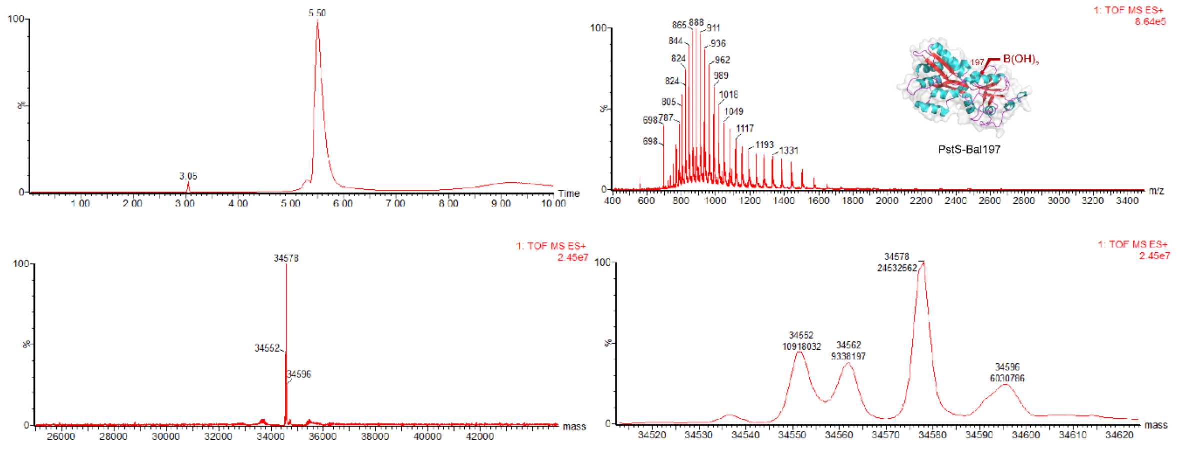

Conversion

| PstS-Fpc197 | PstS-Ala197 | PstS-Bal197 | | |
|---|---|---|---|---|
| | | Bal | Bal-$H_2O$ | Bal-$2H_2O$ |
| 0% | 22% | 11% | 48% | 18% |

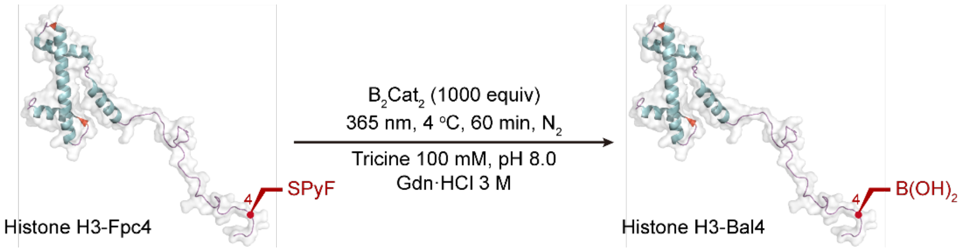

The HistoneH3-Bal4 was prepared according to general procedure 8.

ESI-MS spectrum for the modified histone is shown below.

Calculated mass = 15226 (Bal), 15208 (Bal-$H_2O$)

Observed mass = 15224 (Bal), 15207 (Bal-$H_2O$).

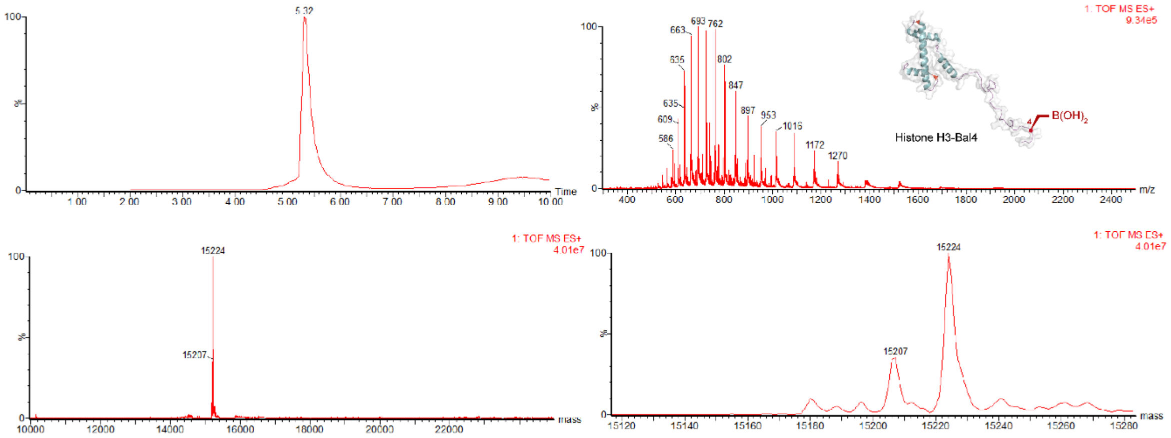

Conversion

| Histone H3-Fpc4 | Histone H3-Ala4 | Histone H3-Bal4 | | |
|---|---|---|---|---|
| | | Bal | Bal-$H_2O$ | Bal-$2H_2O$ |
| 0% | 0% | 71% | 29% | 0% |

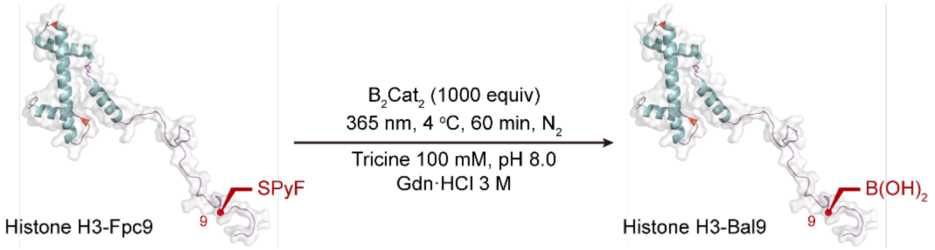

The Histone H3-Bal9 was prepared according to general procedure 8.

ESI-MS spectrum for the modified histone is shown below.

Calculated mass = 15226 (Bal)

Observed mass = 15225 (Bal)

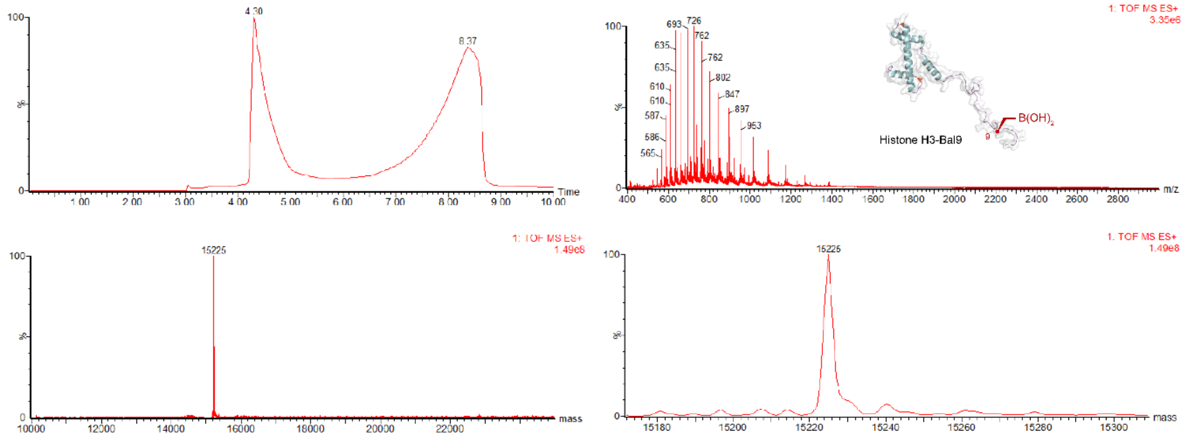

Conversion

| Histone H3-Fpc9 | Histone H3-Fpc9 | Histone H3-Bal9 | | |
|---|---|---|---|---|
| | | Bal | Bal-$H_2O$ | Bal-2$H_2O$ |
| 0% | 0% | >98% conversion | 0% | 0% |

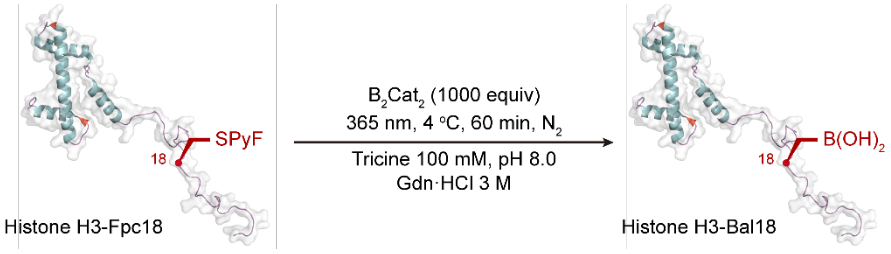

The Histone H3-Bal18 was prepared according to general procedure 8.

ESI-MS spectrum for the modified histone is shown below.

Calculated mass = 15226 (Bal)

Observed mass = 15225 (Bal)

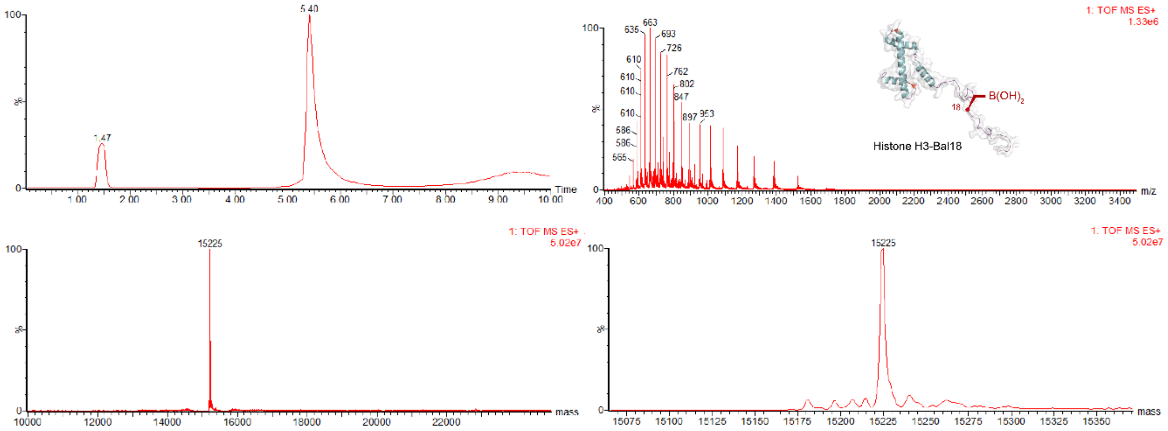

Conversion

| Histone H3-Fpc18 | Histone H3-Ala18 | Histone H3-Bal18 | | |
|---|---|---|---|---|
| | | Bal | Bal-$H_2O$ | Bal-$2H_2O$ |
| 0% | 0% | >98% conversion | 0% | 0% |

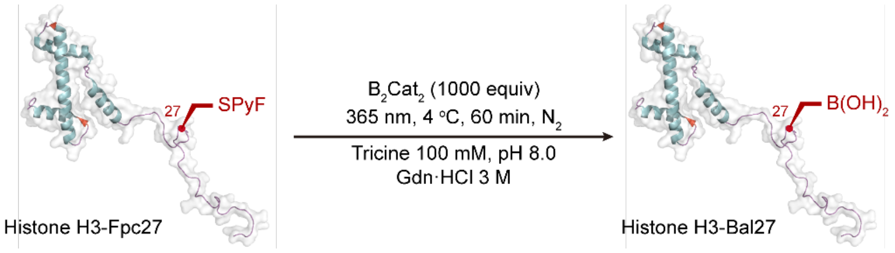

The Histone H3-Bal27 was prepared according to general procedure 8.

ESI-MS spectrum for the modified histone is shown below.

Calculated mass = 15226 (Bal)

Observed mass = 15224 (Bal)

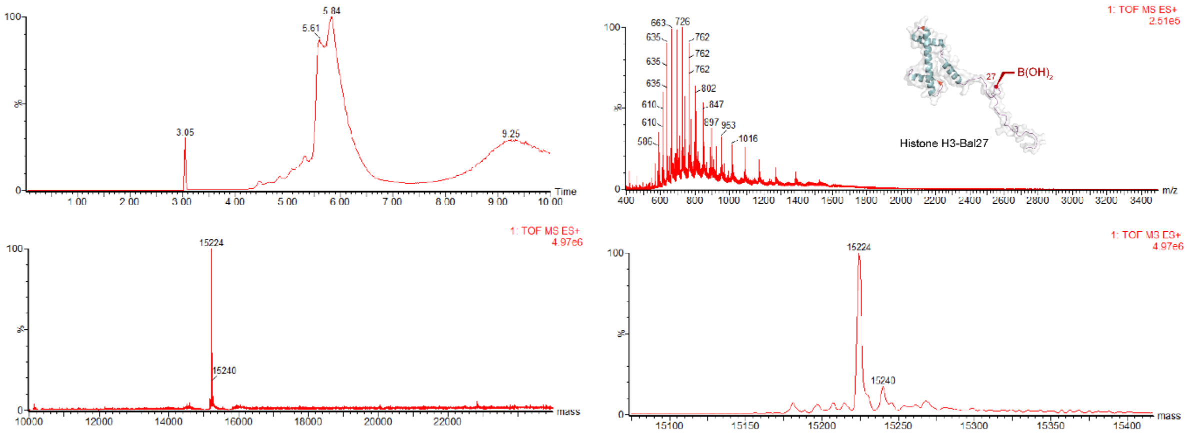

Conversion

| Histone H3-Fpc27 | Histone H3-Ala27 | Histone H3-Bal27 | | |
|---|---|---|---|---|
| | | Bal | Bal-$H_2O$ | Bal-$2H_2O$ |
| 0% | 0% | >98% conversion | 0% | 0% |

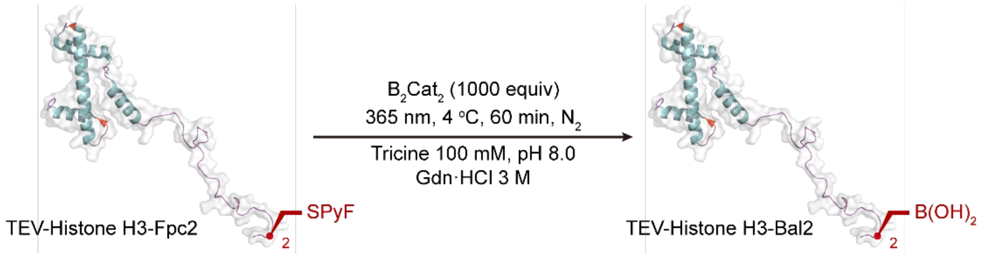

The TEV-Histone H3-Bal2 was prepared according to general procedure 8.

ESI-MS spectrum for the modified histone is shown below.

Calculated mass = 16049 (Bal), 16031 (Bal-$H_2O$), 16013 (Bal-$2H_2O$)

Observed mass = 16048 (Bal), 16029 (Bal-$H_2O$), 16012 (Bal-$2H_2O$)

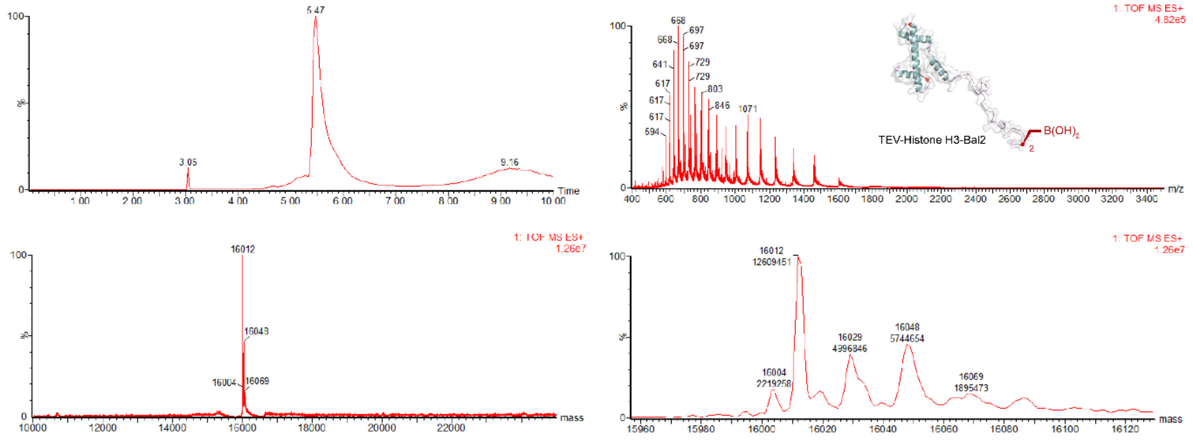

Conversion

| TEV-Histone H3-Bal2 | TEV-Histone H3-Ala2 | TEV-Histone H3-Bal2 | | |
|---|---|---|---|---|
| | | Bal | Bal-$H_2O$ | Bal-$2H_2O$ |
| 0% | 0% | 23% | 22% | 55% |

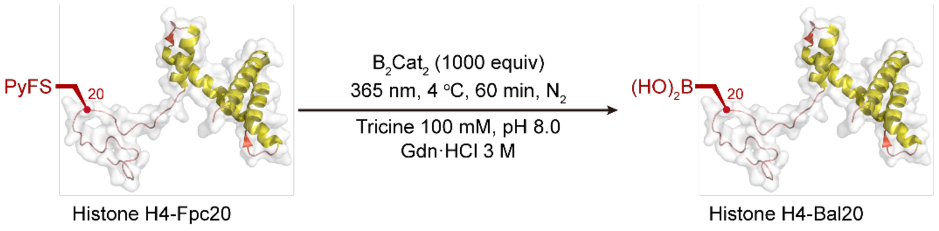

The Histone H4-Bal20 was prepared according to general procedure 8.

ESI-MS spectrum for the modified histone is shown below.

Calculated mass = 11223 (Bal), 11205 (Bal-$H_2O$)

Observed mass = 11222 (Bal), 11204 (Bal-$H_2O$)

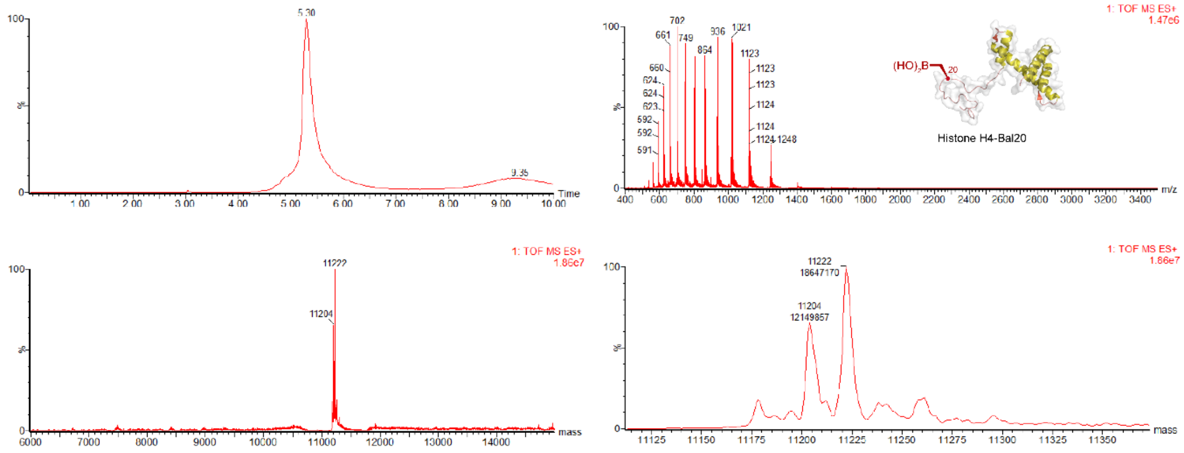

Conversion

| Histone H4-Fpc20 | Histone H4-Ala20 | Histone H4-Bal20 | | |
|---|---|---|---|---|
| | | Bal | Bal-$H_2O$ | Bal-2$H_2O$ |
| 0% | 0% | 62% | 38% | 0% |

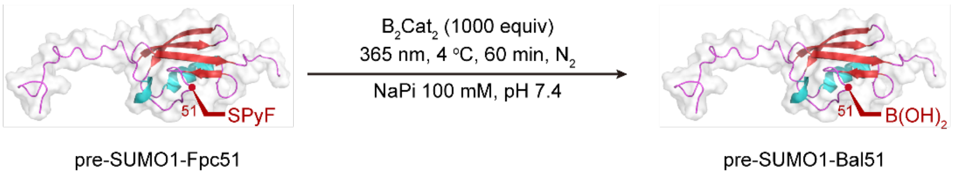

The pre-SUMO1-Bal51 was prepared according to general procedure 8.

ESI-MS spectrum for the modified pre-SUMO1 is shown below.

Calculated mass = 12487 (Bal), 12469 (Bal-$H_2O$)

Observed mass = 12486 (Bal), 12469 (Bal-$H_2O$)

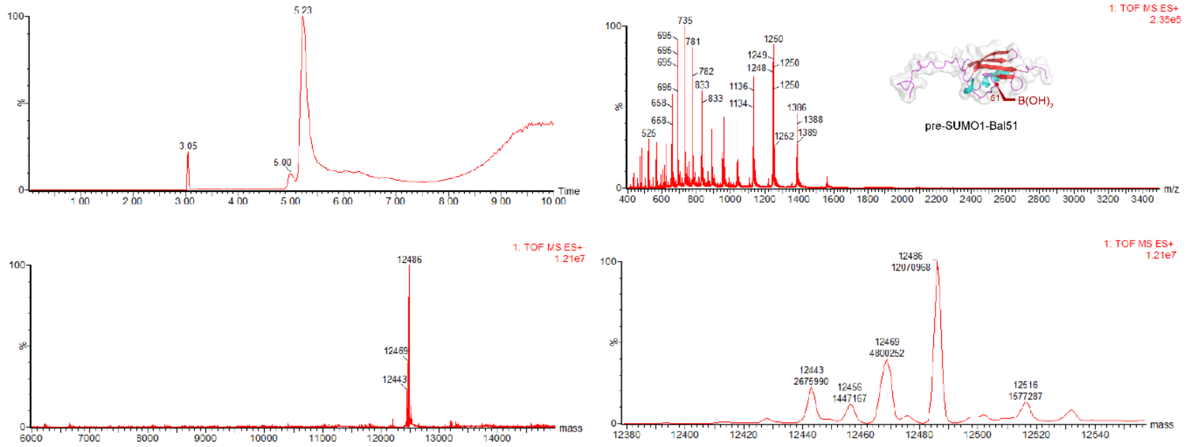

Conversion

| pre-SUMO1-Fpc51 | pre-SUMO1-A51 | pre-SUMO1-Bal51 | | |
|---|---|---|---|---|
| | | Bal | Bal-$H_2O$ | Bal-$2H_2O$ |
| 0% | 15% | 63% | 21% | 0% |

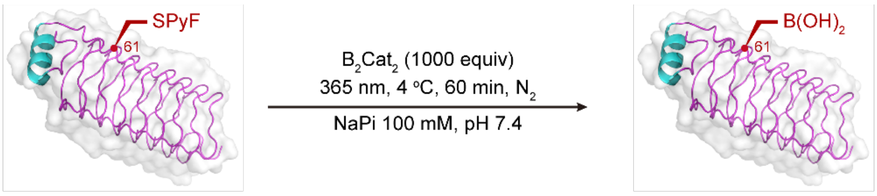

Npβ-Fpc61 Npβ-Bal61

The Npβ-Bal61 was prepared according to general procedure 8.

ESI-MS spectrum for the modified Npβ is shown below.

Calculated mass = 21043 (Bal), 21025 (Bal-$H_2O$), 21007 (Bal-$2H_2O$)

Observed mass = 21040 (Bal), 21024 (Bal-$H_2O$), 21006 (Bal-$2H_2O$)

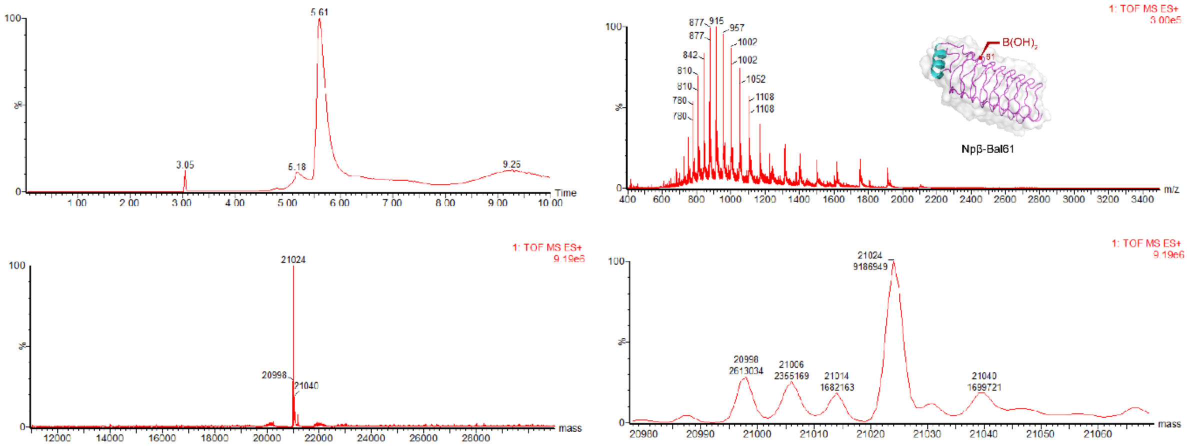

Conversion

| Npβ-Fpc61 | Npβ-Ala61 | Npβ-Bal61 | | |
|---|---|---|---|---|
| | | Bal | Bal-$H_2O$ | Bal-$2H_2O$ |
| 0% | 16% | 11% | 58% | 15% |

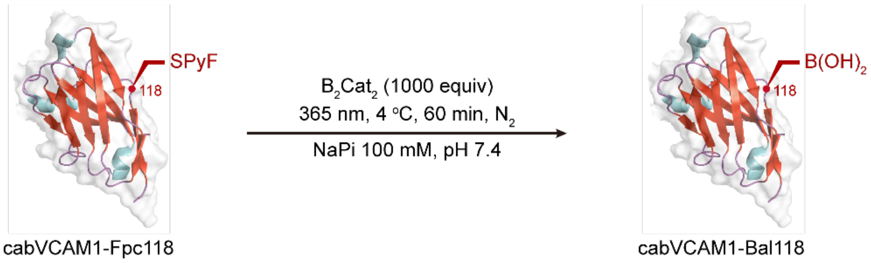

The cAbVCAM1-Bal118 was prepared according to general procedure 8.

ESI-MS spectrum for the modified cAbVCAM1 is shown below.

Calculated mass = 14631 (Bal), 14613 (Bal-$H_2O$), 14595 (Bal-$2H_2O$)

Observed mass = 14631 (Bal), 14613 (Bal-$H_2O$), 14595 (Bal-$2H_2O$)

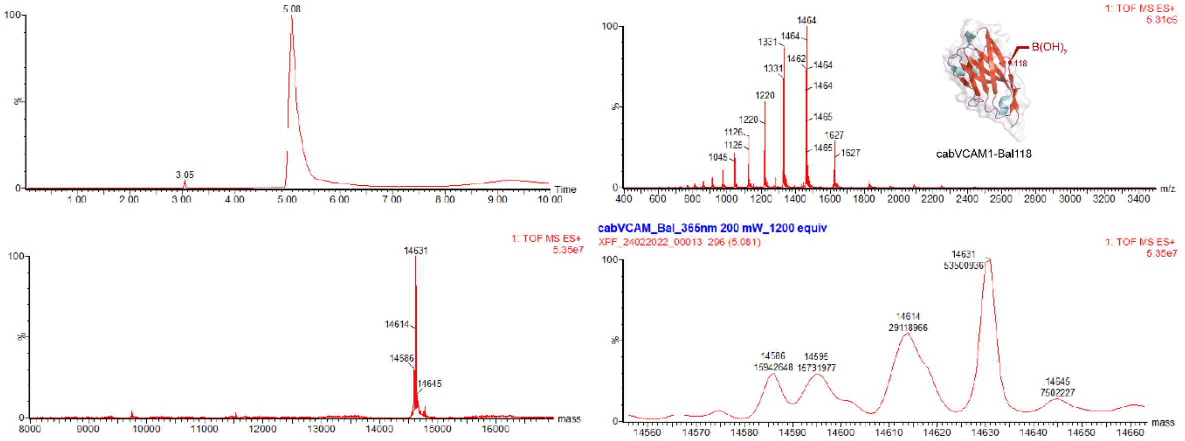

Conversion

| cAbVCAM1-Fpc118 | cAbVCAM1-Ala118 | cAbVCAM1-Bal118 | | |
|---|---|---|---|---|
| | | Bal | Bal-$H_2O$ | Bal-$2H_2O$ |
| 0% | 3% | 46% | 26% | 14% |

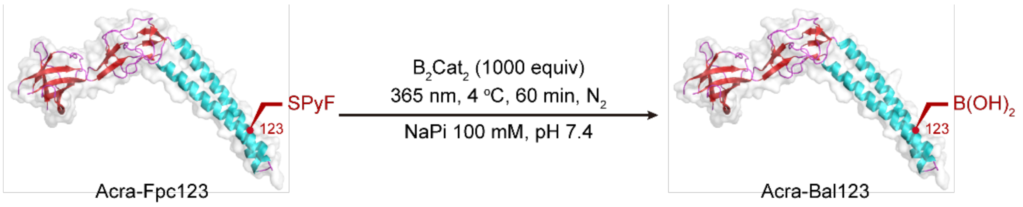

The AcrA-Bal123 was prepared according to general procedure 8.

ESI-MS spectrum for the modified AcrA is shown below.

Calculated mass = 38811 (Bal-$H_2O$)

Observed mass = 38812 (Bal-$H_2O$)

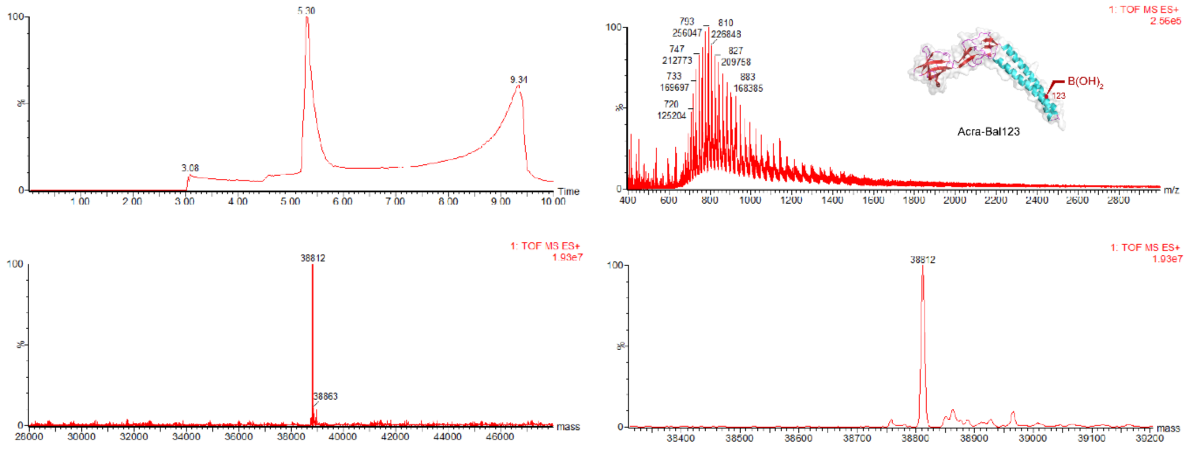

Conversion

| AcrA-Fpc123 | AcrA-Ala123 | AcrA-Bal123 | | |
|---|---|---|---|---|
| | | Bal | Bal-$H_2O$ | Bal-$2H_2O$ |
| 0% | 3% | 0% | >98% conversion | 0% |

**Characterization of the retention of native L-stereochemistry by protein $^{19}$F NMR.**

Protein $^{19}$F-NMR using shift reagent: Determination of the Histone $^{TEV}$H3-L-Bal9. Histone $^{TEV}$H3-L-Bal9 was prepared according to general procedure 8 and then it was desalted to binding buffer (40 mM NaPi, 5 M urea, pH 7.0, 10% $D_2O$) at a final concentration of 97 μM. Then 10 equiv. of chiral shift reagent were added and the sample was vortexed. The sample was transferred to a NMR tube and analyzed on a Bruker AVIII 600 MHz spectrometer equipped with a Prodigy $N_2$ broadband cryoprobe (3,500 scans, d1 = 2 s).

Protein $^{19}$F-NMR using shift reagent of reference epimeric mixture: Determination of the Histone H3-D/L-Bal9. Histone H3-D/L-Bal9 was prepared as reported[3] and then it was desalted to binding buffer (40 mM NaPi, 5 M urea, pH 7.0, 10% $D_2O$) at a final concentration of 144 μM. Then 10 equiv. of chiral shift reagent were added and the sample was vortexed. The sample was transferred to a NMR tube and analyzed on a Bruker AVIII 600 MHz spectrometer equipped with a Prodigy $N_2$ broadband cryoprobe (3,500 scans, d1 = 2 s).

Protein $^{19}$F-NMR using shift reagent with spiking of Histone H3-D/L-Bal9 in binding buffer (40 mM NaPi, 5 M urea, pH 7.0, 10% $D_2O$) was mixed with Histone H3-L-Bal9 in binding buffer (40 mM NaPi, 5 M urea, pH 7.0, 10% $D_2O$), then transferred to a NMR tube and analyzed on a Bruker AVIII 600 MHz spectrometer equipped with a Prodigy $N_2$ broadband cryoprobe (3,500 scans, d1 = 2 s).

### Creation of H3-L-KAc18 and Assessment of Enzymatic Processing as a Substrate using Sirt 2

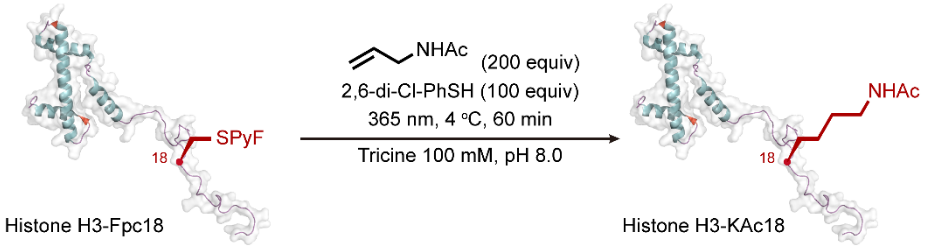

The Histone_H3_18KAc was prepared according to general procedure 5.

ESI-MS spectrum for the modified histone is shown below.

Calculated mass = 15281

Observed mass = 15280

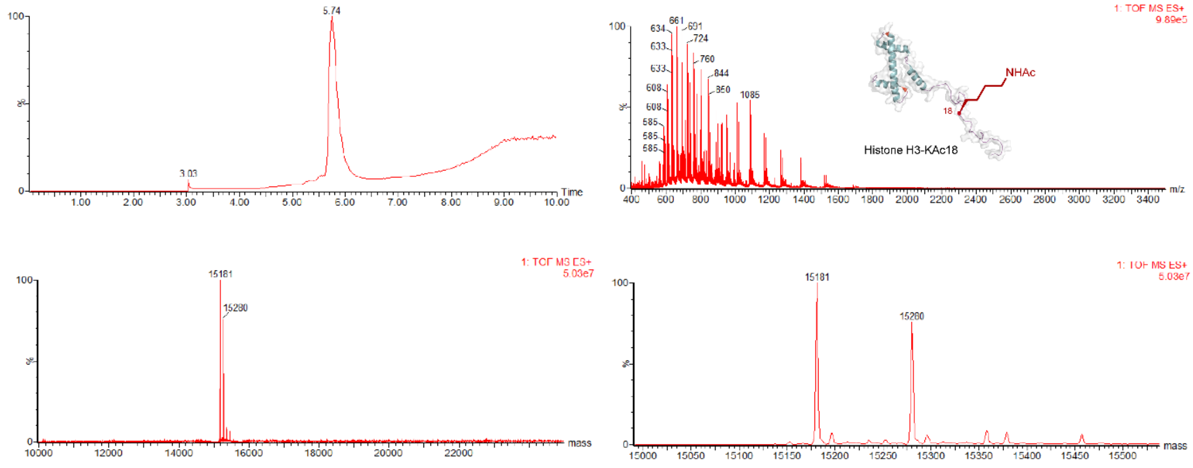

| Histone H3-Fpc18 | Histone H3-Ala18 | Histone H3-KAc18 |
|---|---|---|
| 0% | 55% | 45% |

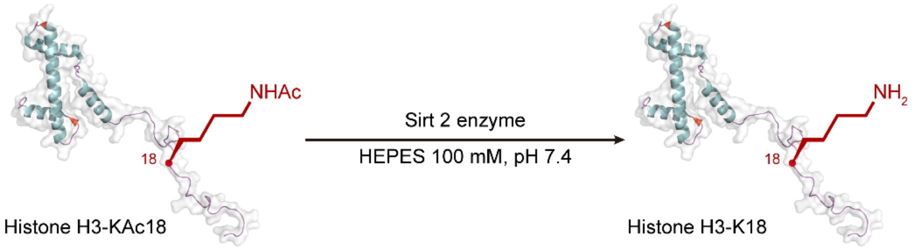

The Histone H3-K18 was prepared according to following procedure.

Histones H3-KAc18 was dialyzed thrice against HEPES buffer (20 mM, pH 7.4), twice for 2 h, once overnight at 4 °C. The solutions were pre-warmed to 37 °C, and Sirt2 (0.5 μg from a stock in the same buffer) and NAD$^+$ (150 μM final concentration from a 10x stock in buffer) were added to solutions containing either Histone H3-KAc18 (20 μM histone, 50 μL final reaction volume). The reactions were shaken at 37 °C, 600 rpm, with aliquots of the crude reaction mixture taken out at 2 time points (30 min, 15 h), diluted (1:50 in $H_2O$ + 1% formic acid) and immediately analysed via LC-MS.

ESI-MS spectrum for the modified histone is shown below.

Calculated mass = 15238

Observed mass = 15238

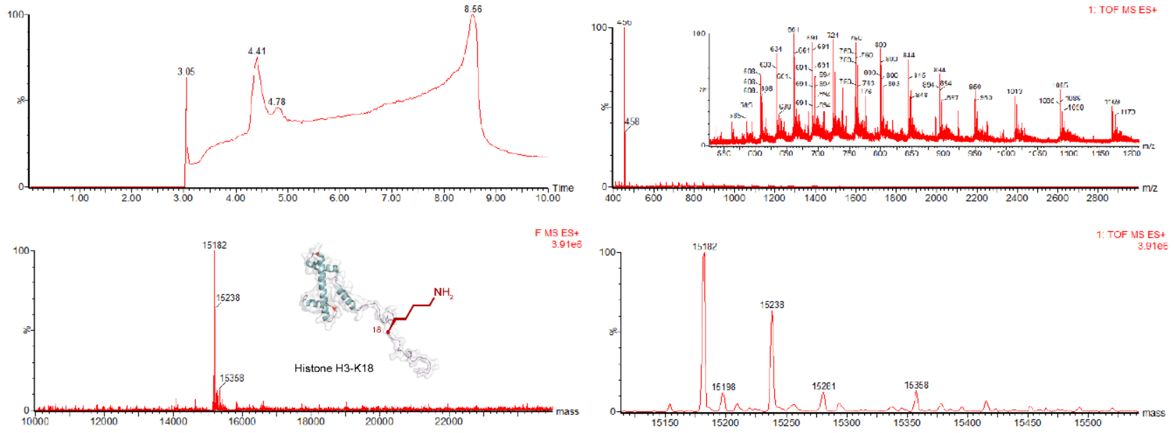

### DFT Calculations for Model PyF Containing Systems

QM calculations were performed using the basis sets shown below using GAUSSIAN 16. Molecular systems with different electron withdrawing substituents on electron acceptor were used as shown.

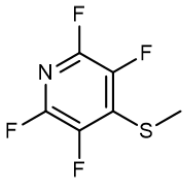

2,3,5,6-tetrafluoro-4-(methylthio)pyridine

$P_1$

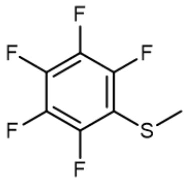

methyl(perfluorophenyl)sulfane

$P_2$

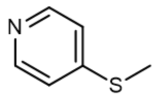

4-(methylthio)pyridine

$P_3$

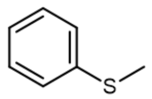

methyl(phenyl)sulfane

$P_4$

LUMO-SOMO gap (eV) for electron acceptors' radical anions:

| | **B3LYP/TZVP** | **B3LYP/631G(d,p)** | **B3LYP/6311G(d,p)** |
|---|---|---|---|
| **P1** | 3.374 | 3.488 | 3.343 |
| **P2** | 3.466 | 3.671 | 3.349 |
| **P3** | 2.620 | 2.826 | 2.731 |
| **P4** | 2.244 | 2.449 | 2.363 |

| | **B3LYP/TZVP** | **B3LYP/631G(d,p)** | **B3LYP/6311G(d,p)** |
|---|---|---|---|
| **P1** | -2845319.162 | -2844573.742 | -2845179.181 |
| **P2** | -3063909.714 | -3063059.803 | -3063739.618 |
| **P3** | -1801974.043 | -1801696.223 | -1801958.628 |
| **P4** | -1759773.025 | -1759518.792 | -1759774.610 |

Optimized energy (kJ/mol) for electron acceptors' radical anions:

S-C(sp3) bond length (Å) comparison for electron acceptors: neutral vs radical anions:

| | **B3LYP/TZVP** | | **B3LYP/631G(d,p)** | | **B3LYP/6311G(d,p)** | |
|---|---|---|---|---|---|---|
| | **Neutral** | **Radical Anion** | **Neutral** | **Radical Anion** | **Neutral** | **Radical Anion** |
| **P1** | 1.83792 | 1.87942 | 1.83225 | 1.89356 | 1.83315 | 1.89074 |
| **P2** | 1.83938 | 1.84511 | 1.83489 | 1.84455 | 1.83458 | 1.84248 |
| **P3** | 1.826 | 1.93578 | 1.82269 | 1.92979 | 1.82122 | 1.92908 |
| **P4** | 1.8388 | 2.0158 | 1.83608 | 1.99276 | 1.83542 | 1.99583 |

S-C(Ar) bond length (Å) comparison for electron acceptors: neutral vs radical anions:

| | **B3LYP/TZVP** | | **B3LYP/631G(d,p)** | | **B3LYP/6311G(d,p)** | |
|---|---|---|---|---|---|---|
| | **Neutral** | **Radical Anion** | **Neutral** | **Radical Anion** | **Neutral** | **Radical Anion** |
| **P1** | 1.77548 | 1.77987 | 1.76877 | 1.74407 | 1.77121 | 1.75031 |
| **P2** | 1.78676 | 1.78593 | 1.78047 | 1.77401 | 1.78101 | 1.77767 |
| **P3** | 1.7778 | 1.74722 | 1.77506 | 1.74119 | 1.77432 | 1.74197 |
| **P4** | 1.80412 | 1.74297 | 1.79954 | 1.73881 | 1.79989 | 1.74001 |

| | **B3LYP/TZVP** | | **B3LYP/631G(d,p)** | | **B3LYP/6311G(d,p)** | |
|---|---|---|---|---|---|---|
| | **HOMO (eV)** | **LUMO (eV)** | **HOMO (eV)** | **LUMO (eV)** | **HOMO (eV)** | **LUMO (eV)** |
| **P1** | -7.198 | -1.984 | -6.835 | -1.439 | -7.121 | -1.862 |
| **P2** | -6.886 | -1.583 | -6.535 | -1.117 | -6.811 | -1.468 |
| **P3** | -6.274 | -0.833 | -6.229 | -0.583 | -6.411 | -0.858 |
| **P4** | -6.441 | -0.959 | -6.095 | -0.476 | -6.298 | -0.781 |

Optimized energy levels for neutral electron acceptors:
